# Using landscape genomics to infer genomic regions involved in environmental adaptation of soybean genebank accessions

**DOI:** 10.1101/2022.02.18.480989

**Authors:** Max Haupt, Karl Schmid

## Abstract

Understanding how crops adapt to specific environmental conditions is becoming increasingly important in the face of accelerating climate change, but the genetics of local adaptation remain little understood for many crops. Landscape genomics can reveal patterns of genetic variation that indicate adaptive diversification during crop evolution and dispersal. Here, we examine genetic differentiation and association signatures with environmental gradients in soybean (Glycine max) germplasm groups from China that were inferred from the USDA Soybean Germplasm Collection (*N* = 17,019 accessions) based on population structure and passport information. We recover genes previously known to be involved in soybean environmental adaptation and report numerous new candidate genes in selection signatures implicated by genomic resources such as the genome annotation and gene expression datasets to function in flowering regulation, photoperiodism and stress reaction cascades. Linkage disequilibrium network analysis suggested functional relationships between genomic regions with selection signatures, consistent with the polygenic nature of environmental adaptation. We tested whether haplotypes associated with environmental adaptation in China were present in 843 North American and 160 European soybean cultivars and found that haplotypes in major genes for early maturity have been selected during breeding, but also that a large number of haplotypes exhibiting putative adaptive variation for cold regions at high latitudes are underrepresented in modern cultivars. Our results demonstrate the value of landscape genomics analysis in genebank germplasm as a starting point for the study of crop environmental adaptation and have the potential to inform future research efforts focused on improved soybean adaptation. Functional validation of candidate genes will support understanding of their adaptive roles and likely enable the transfer of beneficial adaptive variation into modern breeding germplasm.

## Introduction

The environmental adaptation of crop varieties is of central importance in plant breeding and a key objective for successful crop production. Modern plant breeding meets this requirement by classifying crop growing regions into homogeneous mega-environments, which are defined by the spatial distribution of biotic and abiotic stresses in the target population of environments (Gauch and Zobel, 1997). This approach allowed for continuous and efficient development of breeding germplasm with region-specific adaptation. The ongoing climate change and an associated increase in environmental instability results in higher frequency and severity of stressors (Aguirre-Liguori et al., 2021), which demands a higher resilience of modern varieties to future conditions while maintaining high yield levels.In addition, more precise adaptation of specialized cultivars to suitable environments at finer resolution than current mega-environments can increase yields per unit of agricultural land (Resende et al., 2021). These objectives require a detailed understanding of the genetic architecture of environmental adaptation and stress resistance to develop highly adapted varieties for specific target production environments, which is challenging because of the highly polygenic nature of adaptive traits (Snowdon et al., 2021). In plant breeding, functional studies of adaptation are frequently limited to stress experiments conducted under controlled conditions in which few stress factors and their interactions are evaluated with a small number of genotypes. Breeding of new, climate-resistant varieties requires the identification of new sources of beneficial genetic variation for polygenic traits by screening large sets of genetic resources, and a detailed characterisation of the genetic architecture of adaptation.

Landscape genomics is a useful approach to investigate the genetic basis of environmental adaptation without the need for extensive stress experiments because it uses evidence from natural experiments that have shaped genetic diversity during plant evolution. The approach was orginally developed to study the evolution of natural populations by identifying signatures of natural selection in the genome to understand mechanisms of local adaptation (Hoban et al., 2016). Key methods of landscape genomics are outlier tests, which detect differences in allele frequencies between populations, and genotype-environment association tests, which detect correlations between environmental variables and allele frequencies, to identify genomic regions and candidate genes involved in local adaptation (Storfer et al., 2018). The expansion of crop species from their center of domestication and a century-long cultivation under different agroecological conditions resulted in the origin of locally adapted and genetically differentiated landrace cultivars that are products of both natural and human mediated selection (Cortinovis et al., 2020). The application of landscape genomics methods to such landraces therefore allows to investigate the genetic basis of adaptive diversification, supports the discovery of beneficial variation, and provides strategic insights for modern breeding to enhance stress tolerance.

Soybean (*Glycine max*) is a a crop whose area of cultivation has expanded far beyond its center of origin. It was domesticated about 3,000 years ago in China, where subsequent cultivation by ancient farmers led to the selection of more than 20,000 landraces (Carter et al., 2004). By the 15th century AD, soybean was cultivated throughout Asia, and during the 1930s, soybean became a major crop in the Americas. Today, soybean is one of the most economically important crops worldwide.For this expansion, photoperiodic adaptation to narrow latitudinal ranges was an essential trait because it allowed the cultivation of soybean in new environments in which the length of growing seasons differed from the native environment. The genetic basis of photoperiodic adaptation of soybean was therefore of considerable research interest, which led to the identification of *E*-series loci as major maturity genes. Several sets of germplasm have been characterized for the mapped maturity loci *E1 - E4* (e.g., Kurasch et al., 2017; Jiang et al., 2014). This work revealed that the allelic composition of *E* loci shows a geographic distribution that reflects adaptation to different growing regions (Liu et al., 2020a; Goettel and Yong-qiang, 2017). The study of soybean photoperiodic adaptation continues to rely on phenotypic evaluations and regularly confirms the distribution of involved alleles along geographic and environmental gradients (Liu et al., 2021; Zimmer et al., 2021; Wang et al., 2020). These loci are thus also prime candidates for discovery in genome scans of diverse germplasm using landscape genomics methods. Further work on the genetic basis of local adaptation in soybean identified correlations with environmental gradients in East Asia that include humidity, precipitation, temperature, and soil properties. These gradients are thought to influence traits like pod dehiscence (Zhang and Singh, 2020), root development, and tolerance to drought and other abiotic factors (Bandillo et al., 2017).

In this study, we expand previous work on soybean adaptation by focusing on differences in genetic variation among germplasm groups native to China and associations of population allele frequencies with environmental gradients to identify the genetic basis of local environmental adaptation. By combining passport data of the global USDA Soybean Germplasm Collection with the inference of population structure of this material we identified subpopulations and assigned them to their original cultivation environments. Based on the population and environmental classification, we then conducted landscape genomic analyses to identify genomic regions with signatures of adaptive differences between germplasm groups cultivated in different environments. Analysis of linkage disequilibrium, genome annotation and gene expression data was used to elucidate the genomic architecture of adaptation and to elucidate putative functions of candidate genes located in genomic regions with signatures of adaptation. Our results demonstrate the value of landscape genomic analysis of genebank germplasm for studying crop environmental adaptation and its potential to aid soybean adaptation to new environmental conditions.

## Materials & Methods

### Plant Materials

The soybean material analyzed in this study mainly consists of accessions from the USDA Soybean Germplasm Collection (Nelson, 2011). It includes accessions from maturity groups (MG) 000 (early) to MG X (late) and represents a global germplasm sample (*N*=17,019) with emphasis on origin from Asia. It also includes old U.S. cultivars (*N*=208, cultivar release ca. 1895 - 1940s) and a sample of modern cultivars from the U.S. and Canada resulting from public and private breeding programs (*N*=635, cultivar release ca. 1947 - 2016). In addition, we included 160 modern European varieties from maturity groups 000 to II, representing the range of soybean germplasm currently grown in Europe.

### Genotyping Data

SoySNP50K SNP array genotyping data of the USDA Soybean Germplasm Collection (Song et al., 2015, 2013) were obtained from SoyBase (https://soybase.org/snps/). A set of 160 modern European varieties were genotyped with the the same array on an Illumina iScan system at TraitGenetics, Germany. For genotyping, DNA was extracted from dried leaf tissue of a single plant of each variety using the Genomic Micro AX Blood Gravity Kit from A&A Biotechnology and quality and quantity of the DNA were checked on a 3% agarose gel. The USDA and our genotyping data were merged into a single VCF file with VCFtools (Danecek et al., 2011). Non-chromosomal SNPs were removed from the genotype data set, as were markers that were only available for the 160 European varieties. Remaining missing data were imputed using BEAGLE version 5.1 with default settings (Browning and Browning, 2016). Before imputation, a randomly selected subset of 10% marker data set was masked to assess imputation accuracy. The final SoySNP50K dataset included 41,084 SNP markers and imputation accuracy was assessed as 98.30%.

High-density SNP genotypes for accessions from the USDA Soybean Germplasm Collection were obtained from the soybean haplotype map (GmHapMap) resource and from imputations based on SoySNP50K genotypes and resequencing data (Torkamaneh et al., 2021). GmHapMap genotypes for 1,508 accessions that were assigned to one of four subpopulations from China in our analysis of population structure were extracted from the full dataset using VCFtools (Danecek et al., 2011), and 1,088,360 SNP markers with a minor allele frequency ≥ 0.01% in these accessions were retained. This dataset was used for detecting selection signatures in germplasm from China. A second dataset containing the same set of SNP markers was created for varieties in the GmHapMap resource that originated from the USA and Canada, and was used for comparison to germplasm from China for genomic regions with selection signatures. To allow the same comparison with modern European cultivars, the SoySNP50K genotypes for 160 European cultivars were merged with the SoySNP50K and the GmHapMap genotypes for the 1,508 accessions from China. BEAGLE version 5.1 (Browning and Browning, 2016) was used with default settings to impute the GmHapMap markers that were missing for the 160 varieties in this dataset. Imputation accuracy was estimated based on a subset of the masked markers as described above and was 98.55%. The GmHapMap genotypes for the 160 European cultivars were then extracted from this data set.

### Linkage Disequilibrium and LD Network Analysis

LD was analyzed based on SoySNP50K genotypes in the entire set of genebank accessions and within accessions of the *G. max* collection, within old US cultivars, within modern North American cultivars, and within European cultivars. Only SNPs with minor allele frequency ≥ 5% (*N* = 34,358 *SNPs*) were considered for LD calculations. Pairwise LD (*r*^2^) between SNPs was calculated using PLINK (Purcell 2007) via the pairwiseLD function of the synbreed R package (Wimmer et al., 2012). The level of background LD was estimated from LD between physically unlinked markers following Breseghello and Sorrels (2006). After detection of selection signatures, pairwise *r*^2^ values were also calculated with the GmHapMap SNP data for ‘peak’ markers from genomic regions with strong statistical support for population differentiation or association signatures using the LD function of the gaston R package (Perdry and Dandine-Roulland, 2019). In addition, LD was calculated for 10 samples of 1,000 randomly selected SNPs each to approximate the level of LD under a neutral expectation. Only nonpericentromeric SNPs were considered for the sample because observed selection signatures also occurred almost exclusively in non-pericentromeric regions. LD estimates were cubic root transformed to approximate a normal distribution, and permutation t-tests were performed with 1,000 simulations using the twotPermutation function of the DAAD R package (Maindonald and Braun, 2020) to compare the level of average LD between selection signatures with the neutral expectation. LD network analysis (Kemppainen et al., 2015) was performed based on matrices of pairwise LD values between selection signature peak markers by applying three times background LD as threshold to identify distinct clusters in LD networks which may indicate functional gene networks (Boyrie et al., 2021). LD clusters that were located on different chromosomes but nevertheless showed increased LD among selection signatures were then analysed for putative functional relationships between their genes by reviewing their annotation.

### Inference of Population Structure

ADMIXTURE v1.23 (Alexander et al., 2009) was used for the model-based estimation of ancestry in the global germplasm sample using the SoySNP50K genotypes for markers with a minor allele frequency >= 0.05 (*N* = 34,358 SNPs). Since ADMIXTURE assumes approximate linkage equilibrium between markers, the dataset was pruned by removing markers with associations to neighbouring markers that exceeded the level of background LD in sliding windows of 500 kb and 10 markers using the function snpgdsLDpruning from the SNPRelate R package (Zheng et al., 2012), resulting in a marker set of 5,283 SNPs. Ten independent runs of ADMIXTURE were performed for different model scenarios that varied with regards to the assumed number of ancestral populations (*K*). Models from *K* = 2 to *K* = 20 were explored and 10-fold cross-validation was used to quantify model accuracy. CLUMPP v1.1.2 (Jakobsson and Rosenberg, 2007) was used to align and summarize the results of the independent runs for each *K*. Population structure was further summarized by a principal component analysis (PCA) of the full SoySNP50K dataset using the dudi.pca function of the ade4 R package (Dray and Dufour, 2007).

### Inference of Environmental Association and Detection of Selection Signatures

Characterization of traditional cultivation environments is a prerequisite for inferring environmental adaptation in germplasm. To approximate the geographic extent of soybean cultivation in China, a EARTHSTAT dataset of soybean harvested area with a resolution of ≈ 10 km^2^ was used (Ray et al., 2012). These data were overlaid with monthly climate data from the WorldClim v2 (Fick and Hijmans, 2017) database, which provides a resolution of ≈ 1 km^2^. In combination, this allowed extraction of climate data specific to soybean growing areas (Figures 1D and S1). The environmental variables *mean daily temperature, cumulative temperature ≥ 15°C, average daily temperature range, precipitation amount*, and *average daily solar radiation* were calculated from monthly WorldClim mean values for the vegetative and generative phases of soybean cultivation in six growing areas and their latitudinal mean values (Tables 1 and 2). The resulting dataset was summarized by principal component analysis using the dudi.pca function of the ade4 R package (Dray and Dufour, 2007). The definition of the geographic extent of the growing regions and the region-specific soybean growing seasons were taken from the literature (Jamet and Chaumet, 2016; Song et al., 2016; Li et al., 2008; Wang and Gai, 2002). The assignment of the germplasm to these regions was based on the population structure (Figure 1) and passport information provided by the USDA Germplasm Resources Information Network (GRIN) database (www.ars-grin.gov), especially collection site information and maturity group designation.

**Figure 1.**
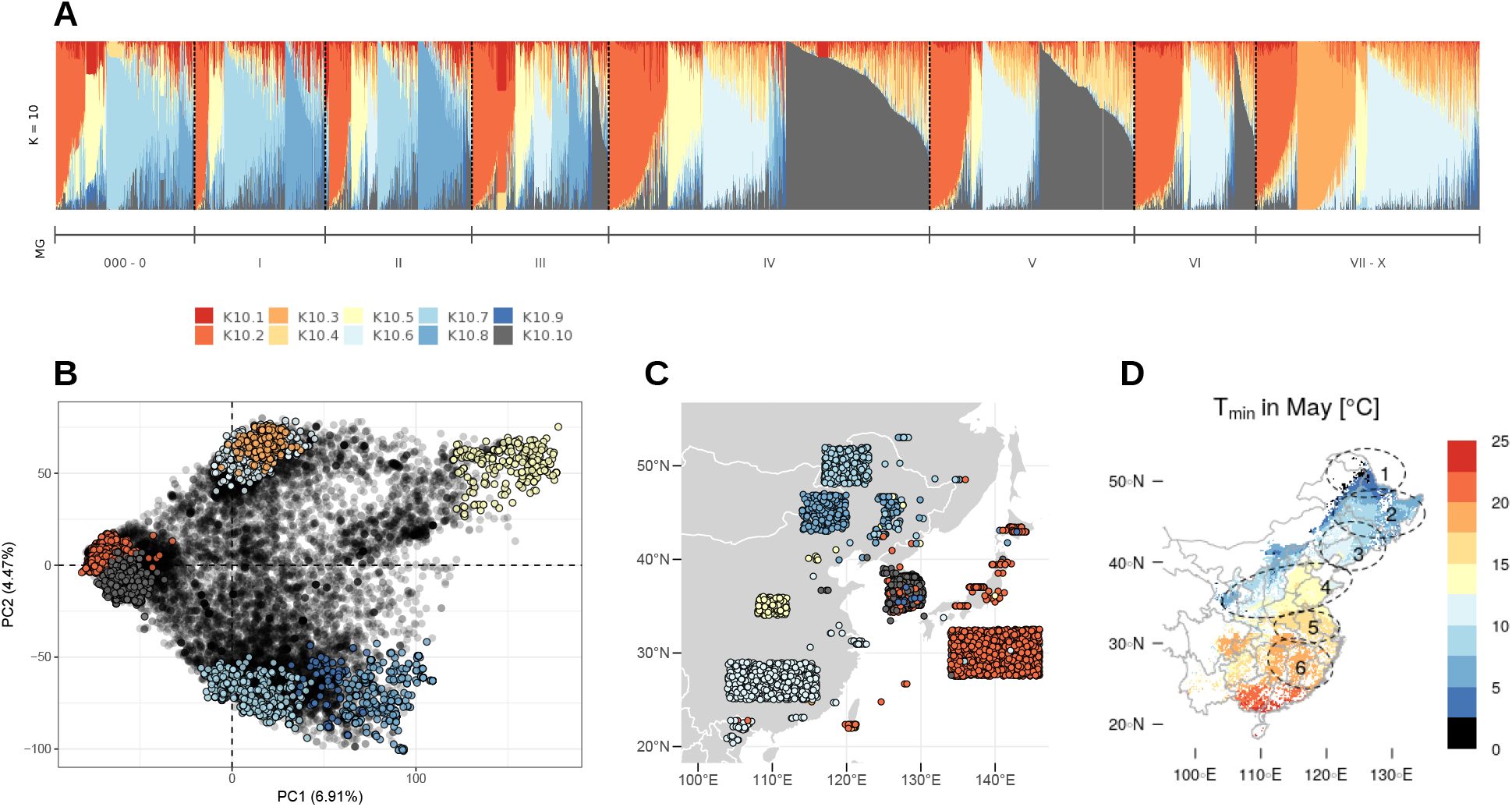
Population structure and origin of soybean germplasm in the USDA Soybean Germplasm Collection. (A) Ancestry proportions as inferred by ADMIXTURE for 17,019 genebank accessions for K = 10. Each accession is represented by a single vertical line and the length of the coloured segments corresponds to the proportion of ancestry from the inferred ancestral populations. (B) Principal Component Analysis of the 17,019 accessions. Conserved accessions with ancestry fractions >= 80% with one of the inferred ancestral groups (K10.1 – K10.10) are highlighted. (C) Geographic origin of conserved accessions from East Asia according to georeference information. Accessions for which georeference information were not available are pooled in blocks and relied on information on the origin at country and province level. (D) Geographic extend of soybean cultivation in China with ellipses 1 - 6 representing growing areas to which germplasm was allocated as detailed in Table 1. Average minimum temperatures for May are shown for soybean growing areas (Figure S1) as an example to illustrate the environmental range.

**Table 1.** Allocation of germplasm groups to growing regions in China based on fractions of ancestry ≥80% with one of the inferred ancestral populations and based on passport information. Regions 1 - 6 correspond to the growing regions given in Figure 1D and Table 2. Subdivision of the K10.7 and K10.8 groups was based on maturity group ratings and group K10.6 was divided according to provincial origin. Two independent genome scans that differed by the included germplasm groups from North-Eastern China were performed: Scenario A including K10.7 and scenario B including K10.8, respectively. NE: North-Eastern China, YR: Yellow River valley, MLRY: Middle to Lower Reaches of Yangtze River region, LSR: Lower Subtropical Region, RP_R1-R8_: Soybean reproductive period from begin of flowering to full maturity.

| Region | Germplasm | MG | N | Sowing Time | $RP_{R1-R8}$ | Scenario |
| --- | --- | --- | --- | --- | --- | --- |
| 1: NE | K10.7 | 000 – 0 | 223 | May 1 | Jul 1 – Sep 30 | A |
| 2: NE | K10.7 | I | 145 | May 1 | Jul 1 – Sep 30 | A |
|  | K10.8 | I | 123 | May 1 | Jul 1 – Sep 30 | B |
| 3: NE | K10.7 | II – III | 152 | May 1 | Jul 1 – Sep 30 | A |
|  | K10.8 | II – III | 250 | May 1 | Jul 1 – Sep 30 | B |
| 4: YR | K10.5 | II – VII | 245 | May 1 | Jul 1 – Sep 30 | A B |
| 5: MLRY | K10.6 | IV – VIII | 318 | Jun 15 | Aug 1 – Sep 30 | A B |
| 6: LSR | K10.6 | IV – VIII | 52 | Jun 15 | Aug 1 – Sep 30 | A B |

**Table 2.**
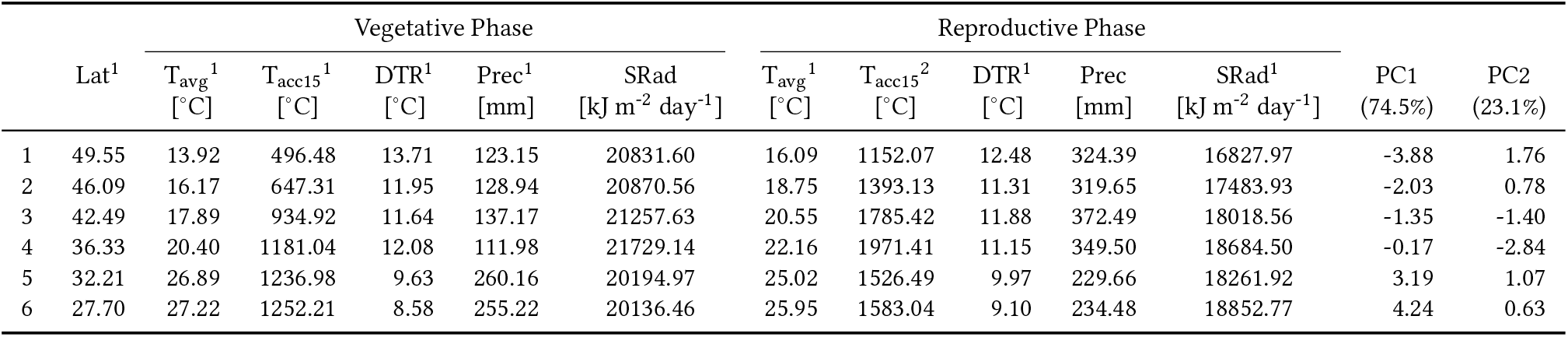
Environmental conditions in six soybean growing regions in China during the vegetative and reproductive phases of soybean development. Lat: latitude, T_avg_: *daily average temperature*, T_acc15_: *accumulated temperature ≥ 15°C*, DTR: *average diurnal temperature range*, Prec: *precipitation amount*, SRad: *average daily solar radiation*, PC1/PC2: position of the growing regions on the first two principal components from a summary of the environmental data by principal component analysis, ^1/2^: significant correlation with with PC1 or PC2 (p < 0.05), Pearson correlation coefficients are given in Figure S4.

The standard covariate model of BAYPASS (Gautier, 2015) was used to identify genetically differentiated regions among soybean populations from the six growing regions and to identify markers associated with populationspecific covariates. Since the population structure analysis identified two genetically distinct groups that both originate from northeastern China and show highly overlapping distribution ranges, BAYPASS was run separately with each of these groups in combination with the other groups to allow detection of convergent and lineage-specific adaptation strategies to the environment in northeastern China. In the following, the two scenarios will be called scenario A and B, respectively (Table 1). For both scenarios, the first two principal components from environmental characterization were provided as population covariate inputs (Table 2) and population allele counts for GmHapMap loci with minor allele frequencies ≥0.01 as genotypic inputs (Supplementary Data Worksheets 2 to 5).

BAYPASS calculated the *XtX* statistic (which corresponds to a SNP-specific *F_ST_* value) for runs that compared genome-wide genetic differentiation between five and six germplasm groups and estimated SNP-wise associations between population allele frequencies and gradients of environmental variables. The *XtX* statistic accounts for the variance-covariance structure between populations to correct for population structure (Günther and Coop, 2013), and the significance of observed *XtX* signals was evaluated with respect to pseudo-observed data sets (POD) of 100,000 SNPs obtained with the R function simulate. baypass as in Gautier (2015). With respect to marker - covariate associations, BAYPASS compares models with and without associations and summarizes model support in terms of Bayes factors (BFs) in deciban units, with a BF > 10 indicating “strong evidence” in favor of a marker - covariate association (and “very strong evidence” at 15 < BF < 20; “decisive evidence” at BF > 20). Five independent runs of BAYPASS were performed for each scenario, and results were summarized as SNP-related means of these five runs of *XtX* statistics and BFs for environmental associations with principal components. SNPs that exceeded the 99% quantile of *XtX* PODs were considered to indicate significant adaptive differentiation between populations, and SNPs with BFs >10 were assumed to be associated with an environmental gradient. SNPs that exceeded these thresholds and were less than 50 kb apart were concatenated and considered a selection signal. For these regions, the signal peaks, start and end positions (±5 kb) were recorded.

### Functional Annotation of Selection Signatures and Gene Ontology Enrichment Analysis

The soybean genome annotation (Wm82.a2.v1) was queried and gene models located within regions with selection signatures were assigned to selection signals. ShinyGO v0.61 (Ge et al., 2020) was used for gene ontology enrichment analyses of gene sets located in genomic regions with differentiation and association signatures and to group genes into functional categories defined by high-level GO terms. Gene ontology enrichment for gene sets belonging to different LD clusters as determined by LD network analysis was also analyzed. Annotation of variants and prediction of functional effects for GmHapMap SNP markers was performed using SnpEff (Cingolani et al., 2012) to categorize the functional effects of the marker set used. To do this, we built our own SnpEff database based on the soybean reference genome and genome annotation GFF (Wm82.a2.v1) available at https://genomevolution.org/coge/GenomeInfo.pl?gid=24686.

### Gene Co-Expression Analysis and Evidence from Differential Gene Expression Studies

To further determine possible functional relationships between signal regions, a transcriptome dataset of 61 US milestone varieties was obtained from the NCBI database (GEO accession GSE99758) to enable the comparison between gene coexpression networks and observed LD clusters. Fragments Per Kilobase Million (FPKM) values were converted to Transcripts Per Million (TPM) values (Kim et al., 2018). The 53,040 x 61 matrix of TPM values was transformed by log_2_(TPM+1), and gene coexpression analysis was performed using the R package CEMi-Tool (Russo et al., 2018). The implemented variance stabilization transformation and signed-network method were used, and a *p*-value threshold of 0.1 was applied to filter correlations of expression data. Gene ontology enrichment analyses for the gene sets obtained were performed using ShinyGO v0.61 (Ge et al., 2020) as described above, and connectivity measures were used to define hub genes in the co-expression network.

We also scanned datasets for differentially expressed genes between vegetative and generative soybean tissues (young leaf versus flower and vice versa, young leaf versus seed 28 days after flower and vice versa, https://soybase.org/soyseq/heatmap/index.php) and in response to dehydration, salt, and drought stress (Rodrigues et al., 2015; Belamkar et al., 2014) to detect possible involvement of candidate genes in developmental and adaptive processes.

### Inference of Haplotype Blocks

A subset of the GmHapMap SNP dataset including modern European and North American cultivars and the germplasm groups from China was used to enable comparisons between these groups at genomic regions with selection signatures at the haplotype level. Haplotype blocks were derived using the block_calculation function of the R HaploBlocker package (Pook et al., 2019) to obtain nonoverlapping blocks, and the results were then queried at genomic positions with selection signatures to determine the amount and length of haplotype blocks and their composition in groups of genebank accessions from China and modern Western breeding material. The same was done for 10 samples of 1,000 randomly selected non-pericentromeric SNPs for comparison.

## Results

### Population Structure and Reconstruction of Germplasm Origin

As a first step, we conducted a population structure analysis to re-establish the link between genebank accessions and their geographic origin via the detection of subpopulations in the USDA soybean collection. This was necessary as a starting point for the downstream landscape genomics analysis because the majority of the recorded collection locations do not necessarily correspond to the primary geographic origin of accessions (Figure 1C).

Model-based ancestry estimation of genebank accessions based on SoySNP50K markers did not provide an unambigous criterion for deciding on the number of ancestral populations (*K*) in our sample of 17,017 accessions. Instead, cross-validation with ADMIXTURE revealed decreasing errors for modeling scenarios with higher *K* values (Figure S2). Models with *K* > 10 showed little improvement in model accuracy over simpler models, which is why we chose *K* = 10 as the value for further analyses. The majority of accessions showed a high degree of genetic admixture (Figure 1A). Consistent with previous work (Xavier et al., 2018; Bandillo et al., 2015), local subpopulations could be identified whose composition largely corresponded to region of origin and photoperiod sensitivity (i.e., maturity group) (Figure 1A-C). To determine the overall provenance of subpopulations, we focused on accessions that shared ancestry values ≥ 80% with one of the derived ancestral groups (Figure S3):With this cutoff, K10.2 formed the largest group (*N*=1,922) and was composed predominantly of accessions from Japan. This group also appears as a tight cluster in the PCA (Figure 1B) and showed a close genetic relationship to K10.10 (*N*=1,322), which consists mainly of germplasm from Korea. Accessions in the Korean group were predominantly classified into MGs IV and V, whereas the group of accessions from Japan ranged widely from MG 000 to VIII. Both groups consisted of large-grained accessions, which is a recognized characteristic of soybean landraces from Japan and Korea (Sedivy et al., 2017), and they were previously reported as distinct clusters (Xavier et al., 2018; Bandillo et al., 2015). Furthermore, we identified four groups originating from China that were not previously described in such detail in the USDA genebank but are consistent with the population structure of fully domesticated germplasm from China (Li et al., 2020; Han et al., 2015): Two groups consisted of accessions collected predominantly in northeastern China (K10.7; *N*=525 and K10.8; *N*=379), one group consisted of accessions from the Huang-Huai Valley (Yellow River Valley, central China) (K10.5; *N*=269), and another group was composed of accessions from southern China (K10.6; *N*=705). The latter has a genetic similarity to an additional group of accessions originating from India and Indonesia (K10.3; *N*=439). The smallest group of accessions was K10.9 (*N*=97) and included mainly U.S. soybean varieties. It should not be considered an ancestral population and is more representative of recent US breeding efforts in the 20th century. The origins of two additional derived ancestral groups (K10.1 and K10.4) remained unidentifiable because no accessions with respective ancestry values ≥ 80% were preserved in the germplasm sample.

For the landscape genomics analysis, we focused on the subpopulations from China because this material represents four well-separated genetic clusters (Figure 1B). They cover the diverse agroclimatic range of soybean cultivation in this region, with subpopulations occupying specific subareas (Figure 1C-D and Table 2). These subpopulations still exhibited considerable intra-group variation in their MG classification, indicating the broad adaptation of germplasm within these subranges. To provide a more detailed correspondence between accessions and their likely original agroenvironments, we used existing maturity group classifications to divide the two northeastern clusters (K10.7 and K10.8) into the MG 000-0, MG I, and MG II-III subgroups (Table 1) and assigned them to latitudinal ranges (Figure 1D). This approach was inspired by the latitudinal segmentation of soybean germplasm according to photoperiod sensitivity observed in North America and Europe (Goettel and Yong-qiang, 2017; Kurasch et al., 2017). For K10.5 and K10.6, we refrained from a clustering by MG because in central and southern China crop rotations and climate allow deviations from standard northern spring cropping practices and require varieties with different maturation behaviors. Instead, we divided the southern Chinese cluster (K10.6) based on provincial origin into groups consistent with previously published ecotypes (Song et al., 2016; Li et al., 2008; Wang and Gai, 2002): the Middle to Lower Reaches of Yangtze River (MLRY) group includes germplasm from Anhui, Henan, Hubei, Jiangsu, and Shanghai provinces, and the Lower Subtropical Region (LSR) group includes material from Fujian, Hunan, Jiangxi, and Zhejiang provinces. In total, we thus obtained eight germplasm groups attributable to six defined growing regions (Tables 1 and 2). These groups were used to identify genomic regions involved in agroclimatic adaptation.

### Differentiation and Adaptation Signatures in Germplasm originating from China

We used genetic differentiation among different germplasm groups from China and associations of local population allele frequencies with environmental gradients to investigate the genetic architecture of soybean environmental adaptation. Based on GmHapMap SNPs, two scenarios were considered: Scenario A included six germplasm groups of northeastern to southern Chinese origin and scenario B included five germplasm groups that originated along the same north-south transect. Both scenarios differed in the included northeastern groups, which were assigned to genetic cluster K10.7 in the case of scenario A and K10.8 in the case of scenario B (Figure 1B-C, Table 1). Since K10.7 and K10.8 overlap in their distribution ranges, they were examined in separate scenarios to recognize mutual and unique adaptation strategies. All germplasm groups, as well as their geographical assignment to cultivation regions, are consistent with previous characterisations of landraces from China (Han et al., 2015; Li et al., 2020, 2008; Song et al., 2016; Wang and Gai, 2002).

Signatures of adaptive genetic differentiation were found in both scenarios and on all 20 chromosomes (Figures 2A, 3A-G and S5). They include genomic regions up to 826 kb in length, encompassing a total of 6,314 annotated genes. 90% of these signals were restricted to regions smaller than 150 kb and included fewer than five candidate genes (Figure 2B). Scenario A resulted in 330 genomic regions with significant genetic differentiation among germplasm groups and scenario B in 288 regions with an overlap of 100 regions (Figure 3E-G). Genomic regions associated with environmental gradients (Table 2) occurred in the same order of magnitude and also showed selection signatures that were mutually detected in both scenarios, as well as genomic regions that were identified in only one of the two scenarios. Between 52 to 92 significantly differentiated genomic regions (based on *XtX*) overlapped with genomic regions identified in the environmental association analysis. Genomic regions identified by both methods included genes known to be involved in environmental adaptation of soybean (Figure 3A-D).

**Figure 2.**
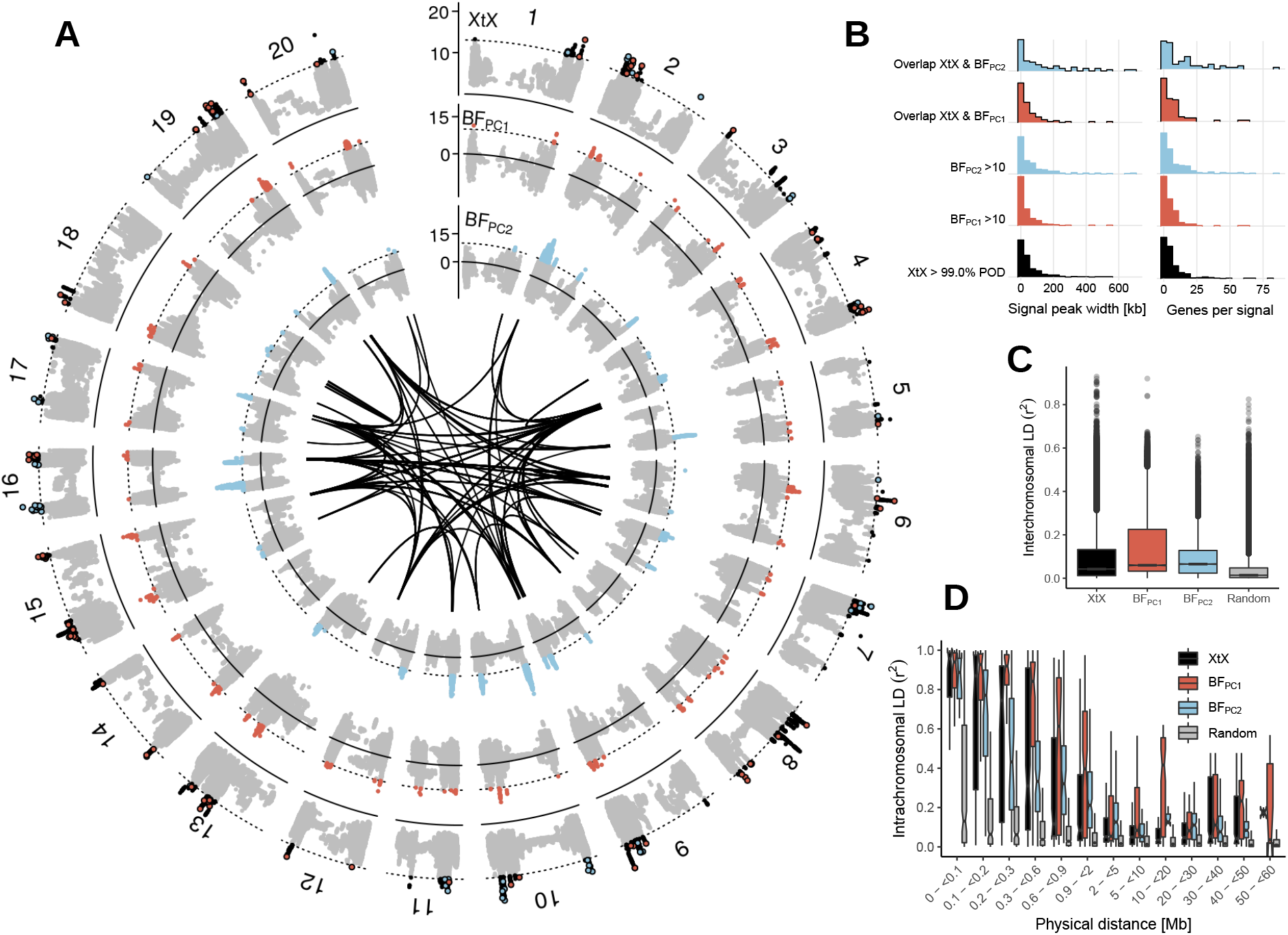
Result summary of the genome-wide scan for selection signatures. (A) Manhattan plot of the *XtX* statistic and of genotype-environment associations with the first two environmental principal components in Bayes factors for all 20 chromosomes in scenario A (for scenario B see Figure S5 and for larger representations of each chromosome see Figures S8-S47). Red and blue points in the *XtX* track represent overlaps between genetic differentiation and association signals with the first and second environmental principal components. The dotted horizontal lines represent the 1% POD significance threshold of the *XtX* statistic and the threshold of BF = 10 deciban. Black lines in the center indicate regions with elevated LD between selection signatures exceeding the level of background LD 3-fold and a minimum physical distance of 5Mb. (B) Width of genomic regions with selection signatures and number of candidate genes. (C) LD between regions with selection signatures from different chromosomes in genetic differentiation and association signals and between randomly selected SNPs. (D) Decay of LD with physical distance between regions with selection signatures and between randomly selected SNPs.

**Figure 3.**
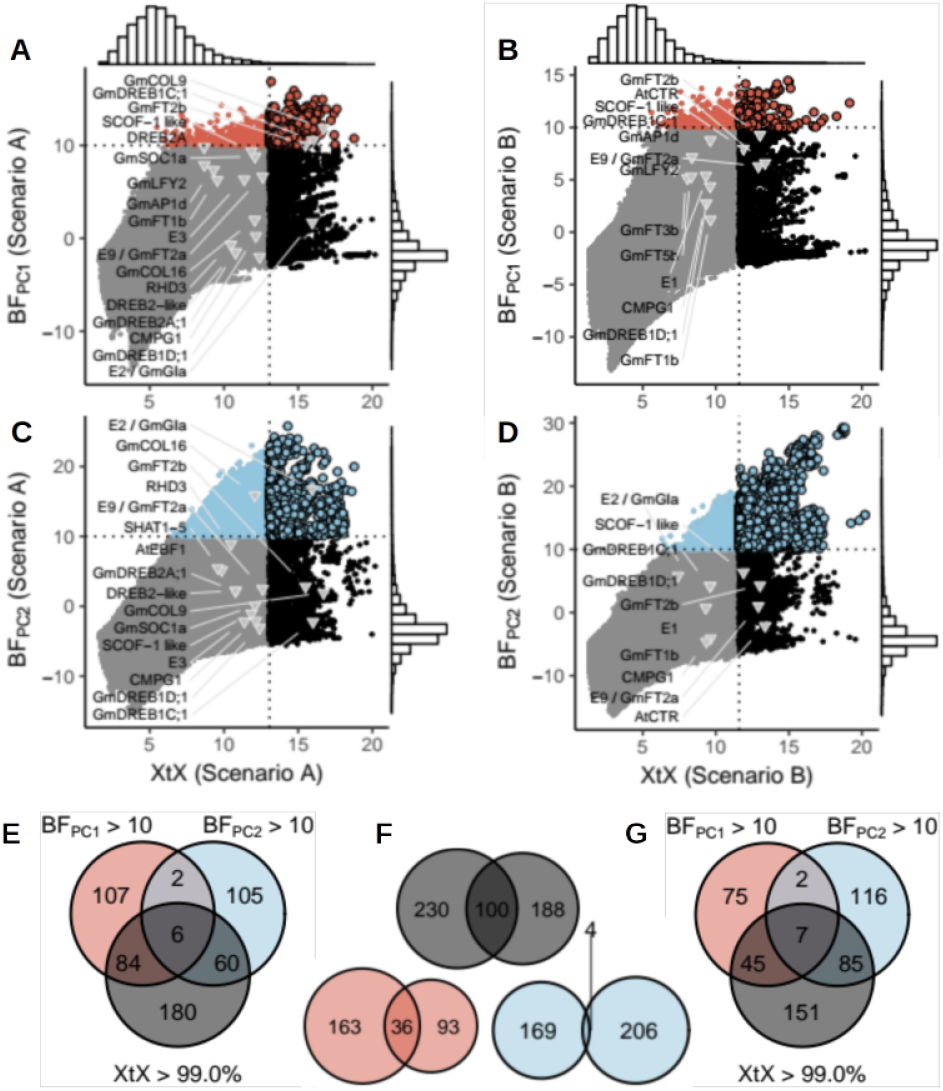
Set relationships of selection signatures among genetic differentiation and association statistics and among scenarios. (A-D) Comparison of genome-wide differentiation estimates (*XtX*) and support for genotype-environment associations (BF) with the first (A-B) and the second (C-D) environmental principal component for scenario A (A,C) and scenario B (B,D). The dotted vertical lines represent the 1% POD significance threshold of the *XtX* statistic, the dotted horizontal lines represent the threshold of BF = 10 deciban. Several soybean genes with agronomic relevance to environmental adaptation that exceeded the 5% *XtX* POD significance threshold and/or exhibited Bayes factors > 5 decibans are labeled. (E-G) Venn diagrams showing the number of genomic regions that displayed differentiation or association signatures and the number of overlaps in scenario A and B, respectively.

A gene ontology (GO) enrichment analysis (FDR < 0.05) detected between one and twenty significantly enriched GO terms for genes located in highly differentiated genomic regions or with a signal of environmental association. These included GO terms related to flowering, photoperiodism and responses to abiotic and biotic stresses (Table S1), supporting a putative involvement of these genes in local environmental adaptation. An inspection of the genome annotation revealed additional evidence for a role in local adaptation, as outlined in the following for chromosome 8. It harbors > 20 genomic regions with selection signatures (Figures 4 and S6), which harbor 64 annotated genes with homologs in other species. Based on their sequence homology, many of these genes are are involved in sensory, regulatory or responsive functions related to environmental adaptation, including the regulation of flowering, abiotic stress responses and nutrient acquisition (Figure 4A, Table 3).

**Figure 4.**
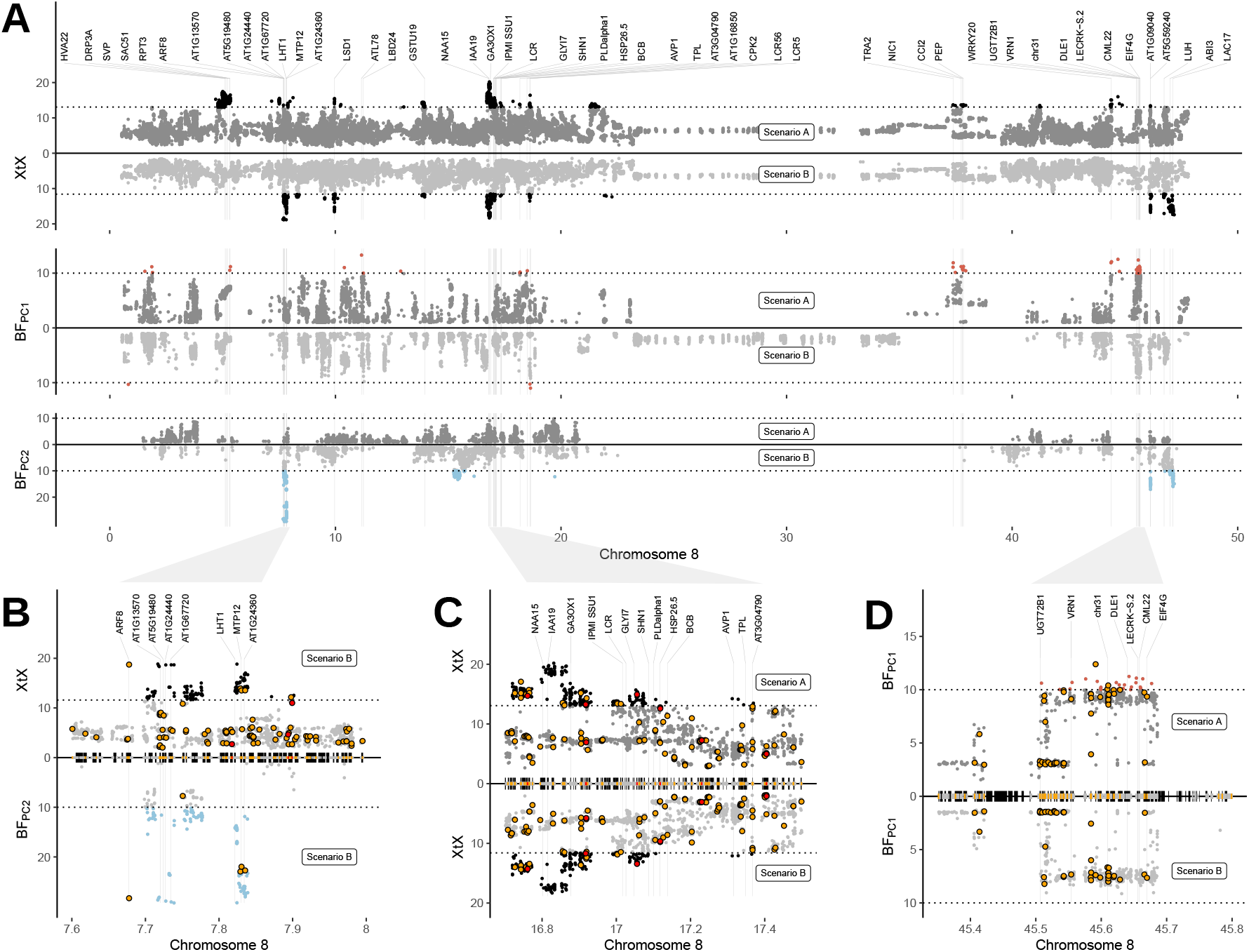
Identification of candidate genes with a putative role in environmental adaptation on chromosome 8. (A) Manhattan plot of the *XtX* statistic and of genotype-environment associations with the first two environmental principal components in Bayes factors for scenario A and scenario B for the complete chromosome 8. Negative Bayes factors are omitted and dotted horizontal lines are analogous to Figure 2A. Labeled candidate genes are listed in Tab 3. The unit of the x-axis is Megabases (Mb). (B-D) Detailed view of genomic regions with selection signatures. Black blocks on the x-axis indicate the positions of predicted gene models. Red and yellow blocks and points indicate non-synonymous variants with high and moderate impacts on protein function. Further close-ups of regions with candidate genes on chromosome 8 are included in Fig. S6.

**Table 3.**
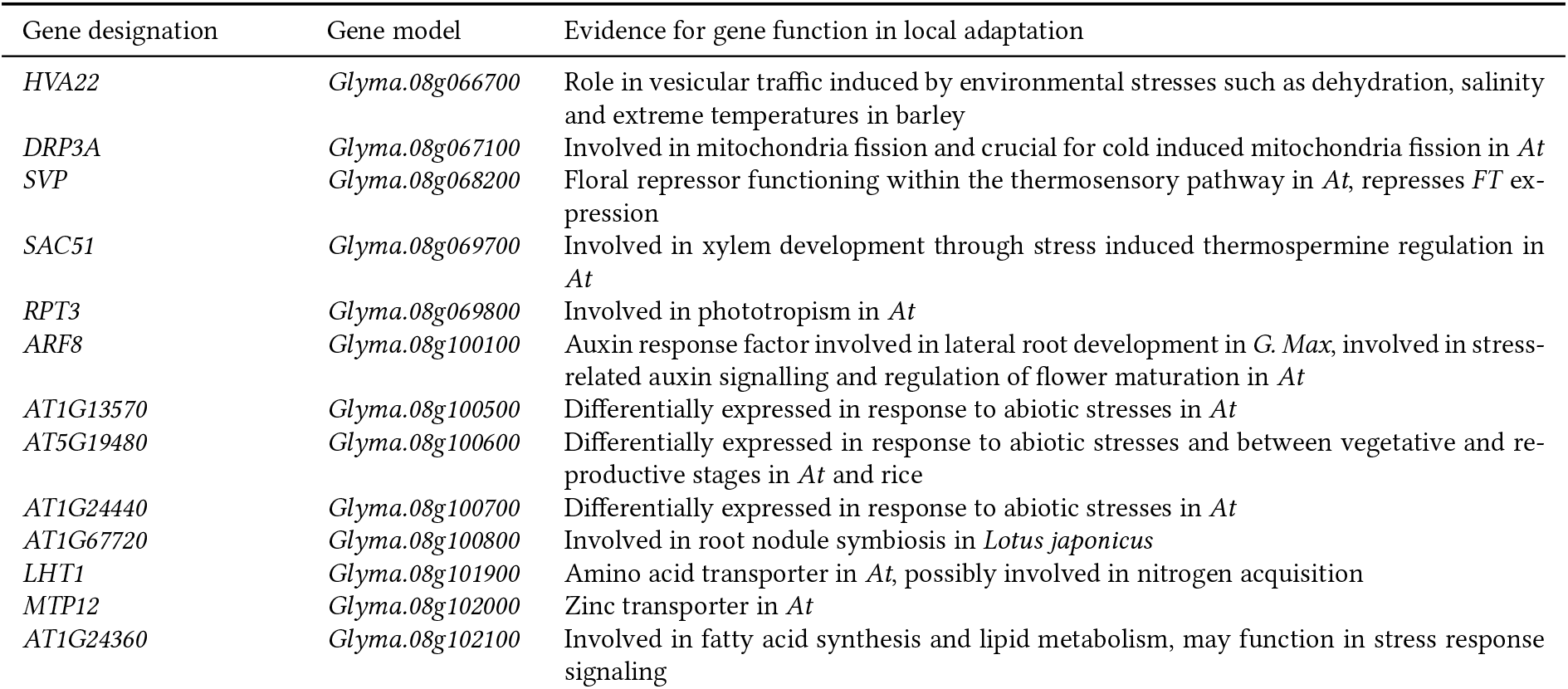
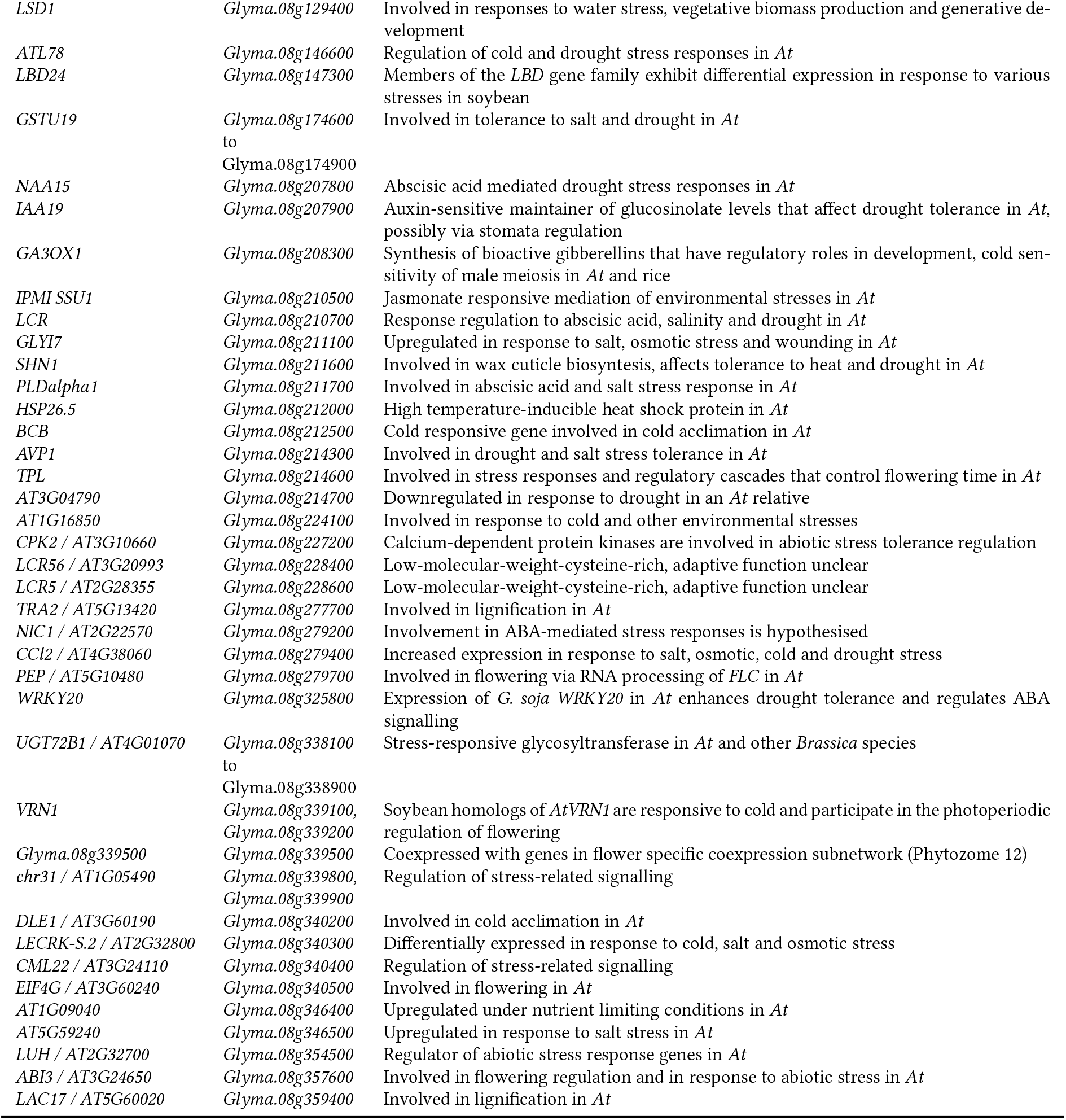
Overview of candidate genes located in genomic regions with selection signatures on chromosome 8 (as detailed in Figure 4) and evidence for their putative involvement in environmental adaptation. The relationship between gene designation, gene model and evidence for gene function was obtained from SoyBase. Information on the physical position of the gene models is included in Worksheet 6 in the Supplementary Data. Gene designation corresponds to gene labels in Figure 4. *At: Arabidopsis thaliana*.

Most genomic regions with selection signals included promising candidate genes (Figures 4B-D and S6) suggesting that further functional validation is required. Environmental adaptation may involve complex interactions of small effect loci (Hoban et al., 2016), which raises the hypothesis of selection signatures that result from synergistic fitness effects of linked genes which may facilitate detection of environmental adaptation in selection scans.

To prioritise candidate genes, we predicted potential functional effects of non-synonymous variants using SnpEff. A small proportion of SNPs (0.3%) were classified to have a strong functional effect (e.g., loss of function) and 5.7% of SNPs to have moderate impacts such as a change in protein effectiveness (Cingolani et al., 2012). This information may allow to prioritize candidate genes with non-synonymous variants over neighboring genes in selected regions and may even include causative variants responsible for selection signatures (e.g. Figures 4B and S6), but causal inference especially of closely linked genes (e.g. Figure 4C-D) remains speculative without further validation. Many genomic regions with selection signatures were also devoid of non-synonymous variants among SNPs with strong statistical support for selection (e.g. Figure S6), suggesting that causal variants were not included, but linked to markers in our SNP set.

Since a substantial proportion of genomic regions appeared in more than one selection test, they may constitute robust signals (Hoban et al., 2016). On chromosome 8 we identified genomic regions with significant differentiation in both scenarios A and B (e.g. Figure 4C) as well as regions with strong genetic differentiation and association to environmental variables (e.g. Figures 4B and S6).

Selection signatures on other chromosomes did not differ substantially from chromosome 8 (Figures 2A, S5 and S8-S47. Worksheet 6 in the Supplementary Data contains all 6,314 genes within genomic regions with selection signatures. A comparison with previous studies revealed that several known adaptation genes did not exceed our significance thresholds (Figure 3A-D) suggesting that our approach is conservative. For example, maturity loci *E2* (Figure S7) and *E9* (Figure 5) show strong selection signatures, whereas *E3* and *E1* showed differentiation between germplasm groups (Figure 3A-D), but did not achieve statistical significance. The limited differentiation at the major maturation locus *E1* in germplasm from China is well known (Zhou et al., 2015) and likely reflects a low frequency of the *e1* allele, which is rare even in landraces from northeastern China where the allele is supposed to be beneficial (Langewisch et al., 2017). These observations show that results from landscape genomic studies are population specific and highlight the limitations of genome scans because polygenic and functionally redundant trait architectures may remain undetected (Hoban et al., 2016).

**Figure 5.**
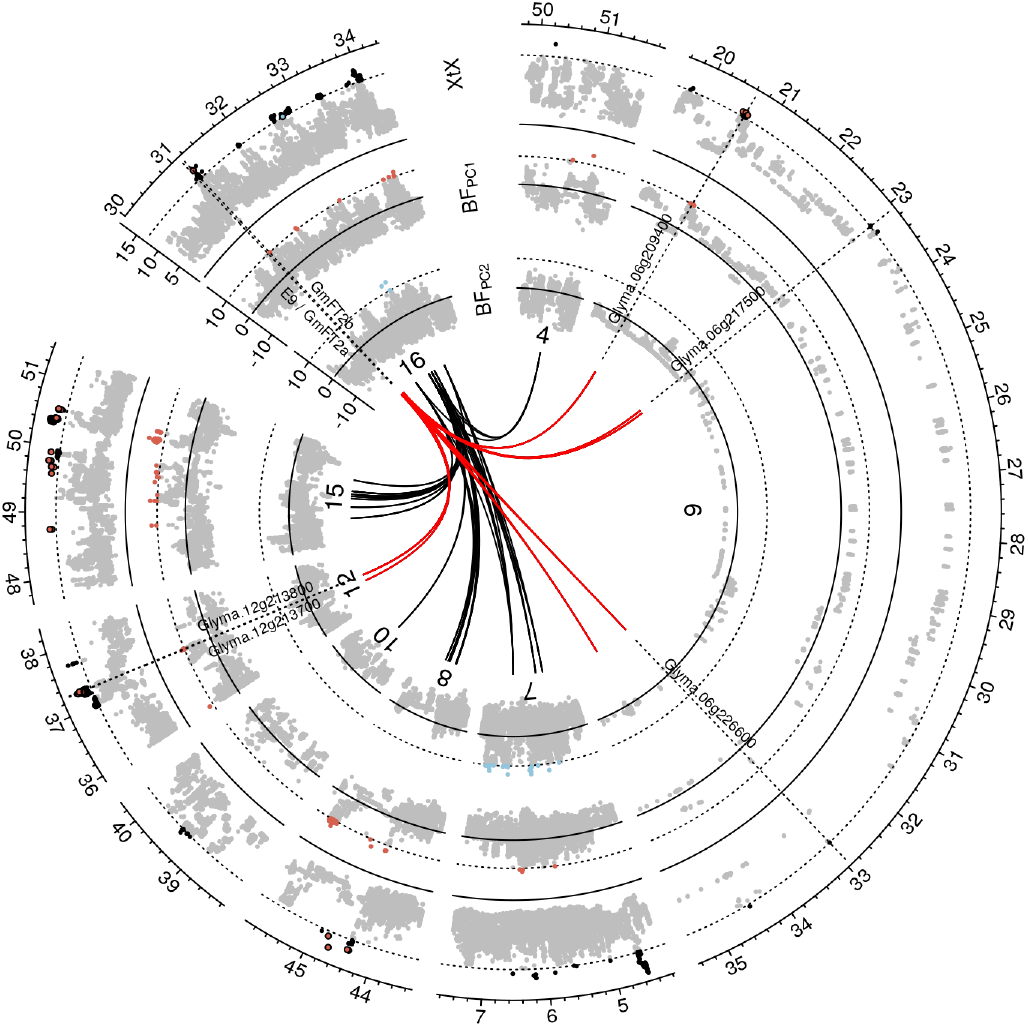
Linkage disequilibrium networks among genomic regions with selection signatures. Detailed view of a Manhattan plot corresponding to Figure 2A for genomic regions that display LD > 3-fold background LD with a region on chromosome 16 that includes the genes *E9*/ *GmFT2a* and *GmFT2b* which are involved in the regulation of flowering. Only outgoing LD connections (from chromosome 16) are displayed in form of black and red lines. Red lines indicate putative members of a network involved in flowering regulation.

### Linkage Disequilibrium among Genomic Regions with Selection Signatures

Many genomic regions with differentiation and/or association signatures showed increased mean pairwise LD (*r*^2^) between SNPs with highest values in differentiation or association statistics within each region (peak markers). In addition, pairwise *r*^2^ values of peak markers from different genomic regions with selection signals were consistently higher than between pairs of markers randomly sampled from the whole genome (p < 0.001; Two-sample permutation t-test). We observed this pattern for comparisons of putatively selected genomic regions both within and between chromosomes (Figure 2C-D). The average background LD between unlinked markers located on different chromosomes was *r*^2^ = 0.21, and this value was exceeded two to five times more frequently between markers with selection signatures on different chromosomes than between randomly selected markers. For higher *r*^2^ values this ratio increased by a factor of five in scenario B and 38 in scenario A (Table S3). Such strong disequilibrium levels between unlinked genomic regions (Figures 2A and S5) suggest epistatic selection of polygenically inherited traits that may contribute to local environmental adaptation (Boyrie et al., 2021) To further investigate this hypothesis, we performed LD network analysis for genomic regions with selection signatures by applying 3-fold background LD as a threshold to identify discrete clusters that might indicate functional associations. LD clusters were identified in both scenarios, but individual clusters mostly consisted of signals localized in proximity of each other within individual chromosomes (Figures 6A-B, S48 and S49). Since LD clusters may originate from selective sweeps or inversions that inhibit local recombination (Kemppainen et al., 2015) or from neutral variation in recombination rate across the genome (Booker et al., 2020), we focused on LD clusters that are distant in the genome (minimum distance of 5Mb) and include interchromosomal LD connections because they may represent functional epistatic interactions. In scenario A, two such instances of interchromosomal LD clusters with selection signatures (Figures 6A-B and S48) and in scenario B, three such clusters were observed (Figure S49). Gene Ontology enrichment analysis for these LD clusters did not reveal significantly enriched GO terms for the genes located within regions with selection signatures (data not shown). One cluster in scenario A included a soybean homolog of *AtFT2b* (Chen et al., 2020) and the flowering and maturity locus *E9*, which is a soybean homolog of *AtFT2a* (Zhao et al., 2016). Both are located on chromosome 16 in close proximity and this region was in high LD with regions on chromosomes 6 and 12 (Figure 5). These regions also show selection signatures and contain candidate genes with a putative role in flowering regulation. For example, the gene model *Glyma.06g209400* was assigned the GO term *regulation of flower development* and the *Arabidopsis* homolog of *Glyma.12 g213700* (*AT4G21200*) is involved in flower initiation (Mateos et al., 2015). Other candidates of this cluster might be related to the regulation of the heat stress response during flowering: The homolog of *Glyma.06g226600* (*AT3G14200*) is a stress-responsive gene involved in pollen tube growth in *Arabidopsis* (Cartagena et al., 2008), while *Glyma.12 g213800* (*AT3G01140*) is a regulator of cuticle biosynthesis inducible in petals by heat stress (Chen et al., 2019).

**Figure 6.**
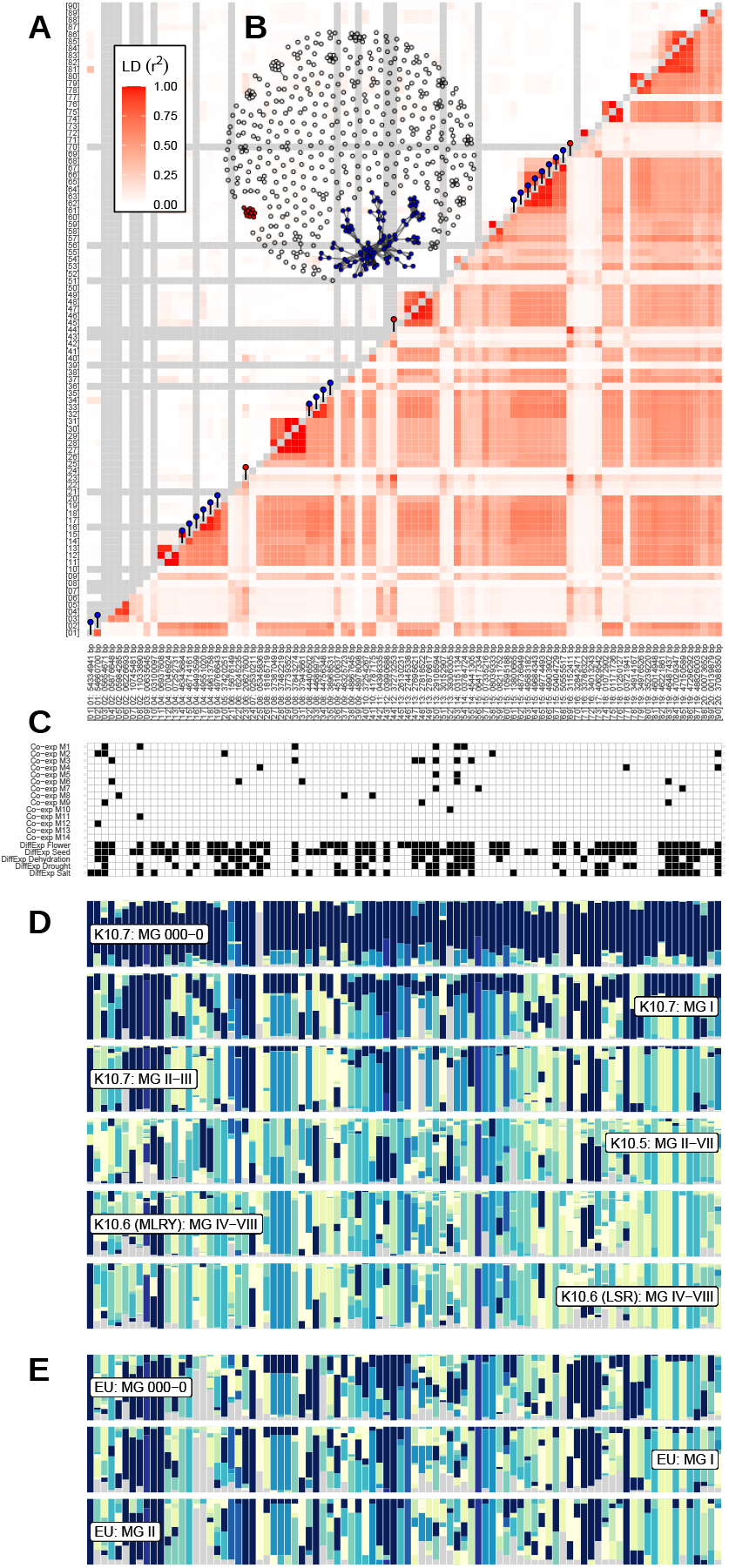
Comparison of haplotype block proportions and of LD between genomic regions with selection signatures in germplasm from China and in modern European varieties. (A) LD estimates among 90 genomic regions with overlapping genetic differentiation and genotype-environment association signatures (PC1) observed in scenario A for germplasm groups from China (below diagonal) and in modern European soybean varieties (above diagonal). Red and blue pins on the diagnonal indicate regions that clustered in LD network analysis. For a complete overview of LD among all selection signatures in scenarios A and B see Figures S48 and S49. (B) LD network of 544 genomic regions with selection signatures in scenario A for germplasm groups included in scenario A. Two clusters (92 and 10 regions, respectively) comprising LD connections exceeding the level of background LD 3-fold and a minimum physical distance of 5Mb are highlighted in blue and red. (C) Occurrences of genes from 14 co-expression modules and of genes that have been found to be differentially expressed in development and in response to abiotic stresses is indicated in black for the displayed 90 genomic regions. (D) Haplotype block proportions in germplasm groups from China in the 90 genomic regions mirror a latitudinal gradient. (E) Haplotype block proportions in modern European varieties in the 90 genomic regions reveal an underrepresentation of haplotype blocks of North-Eastern Chinese descent that could harbour adaptive variation to high-latitude cold regions.

To further investigate possible functional connections between genes in LD clusters, we analyzed gene coexpression networks. In the RNA expression profiles of 61 US milestone cultivars that we used as proxies for our germplasm from China 4,593 out of 53,040 predicted genes were present and included in the construction of a coexpression network with the CEMiTool R package. The resulting network included 14 modules containing between 59 and 941 member genes. Twelve coexpression modules contained a wide variety of significantly enriched GO terms (Table S2). A total of 463 coexpressed genes were located in genomic regions with selection signatures (e.g. Figure 6C). The average inter- and intramodular connectivity measures of these putative targets of selection were significantly higher than network averages (p = 0.0032 and p = 0.0001; Welch’s two sample t-test) suggesting a functional role in local adaptation (Chateigner et al., 2020). On the other hand, in genomic regions with selection signatures, genes belonging to a LD cluster overlapped little with genes contained in a coexpression module (e.g. Figure 6A and C) suggesting that there is no simple relationship between both entities. However, this observation does not reject the hypothesis of a functional association between LD cluster genes because the transcriptome data cover only a small part of the gene expression variation. In soybean, gene expression is variable among different tissues and developmental stages (Severin et al., 2010). Since the available expression profiles covered gene expression only at a very early vegetative stage of soybean development, and expression of adaptation genes can be context-dependent (Barrera-Redondo et al., 2020) and population-specific (Blanc et al., 2020), the available data from US milestone cultivars show a low power to detect such a relationship. We also screened additional datasets with genes that are differentially expressed between vegetative and generative tissues of soybean or in reaction to abiotic stresses (Rodrigues et al., 2015; Belamkar et al., 2014) for overlaps with the gene content in our selection signatures. Numerous differentially expressed genes are located in genomic regions with selection signatures (e.g. Figure 6C), but they failed to specifically elucidate the potential functions of gene LD networks. This data suffers the same limitations that were mentioned above for the co-expression data. Another caveat is the observation that LD clusters identified in genebank accessions from China are not observed in US and European cultivars (Figures 6A, S48 and S49) suggesting that adaptive variation or epistatic interactions in the former may be depleted in the latter group. Taken together, these analyses show that LD network analysis identifies clusters of genes among genomic regions with selection signatures on different chromosomes. However, additional evidence for functional interactions of genes located within distant genomic regions with strong disequilibrium remains anecdotal and plausible only for the genetic network involving maturity locus *E9* and further genes involved in flowering time control.

### Frequency of putative adaptive haplotypes in U.S. and European cultivars

To utilize genomic regions with selection signals and putative adaptive genetic variation in genetic resources for breeding of elite varieties, we identified haplotype blocks that are shared between germplasm from China and modern breeding material from North America and Europe. In China, variation of haplotype blocks in genomic regions with selection signatures showed a latitudinal gradient (e.g. Figures 6D and S50-S53). Canadian and Central European soybean breeding programs aim for improved adaptation to cooler regions at higher latitudes, for which little genetic variation is currently available (Saleem et al., 2021; Hahn and Würschum, 2014; Cober et al., 2013). New variation for cold adaptation may therefore be found in haplotype blocks in landraces that are adapted to the far north of northeastern China and adjacent regions in Russia (Haupt and Schmid, 2020; Jia et al., 2014). We therefore investigated whether haplotype blocks in genomic regions with selection signals in germplasm from China are present in modern cultivars from North America and Central Europe, and found variable proportions of haplotype blocks with ancestry from northeast China in these cultivars (e.g. Figures 6E and S50-S53). For example, the genomic region harboring the maturity locus *E9* in the earliest maturing modern North American and European cultivars (MG 000-0) was nearly fixed for the haplotype block prevalent in material from northeastern China, whereas haplotype blocks with ancestry in material from Southern China rapidly increases in frequency in later maturity group cultivars. Similar patterns of variation in haplotype frequency were observed in other genomic regions and appear to reflect selection for improved adapation to specific latitudes. This interpretation is consistent with an increase of haplotype blocks originating from northeastern China in modern early-maturing cultivars when compared to older cultivars (e.g., Figures S50-S53). Overall, haplotype blocks of northeastern ancestry with signatures of selection were generally more common in early-maturing cultivars than in later-maturing cultivars, but they are still underrepresented in many genomic regions. These observations suggest a strong potential for further introgression of potentially adaptive alleles for cultivation in cool high latitude regions. Haplotype block profiles of modern early-maturing North American and European cultivars (e.g. Figures S54 and S55) show that such introgression could be achieved within the modern germplasm pool by targeted crosses of varieties with complementary haplotype block profiles and subsequent marker-assisted selection to develop breeding material enriched in haplotype blocks with ancestry from northeastern China.

## Discussion

We applied landscape genomics analyses within soybean germplasm that originates from China and which is part of the USDA Soybean Germplasm Collection (Nelson, 2011). Like similar studies in other crop species (Lei et al., 2019; Romero Navarro et al., 2017; Russell et al., 2016; Lasky et al., 2015), we identified hundreds of candidate loci with selection signatures (e.g. Figure 2A, Figure 3). These regions included genes which control flowering, abiotic stress response and nutrient acquisition. The latter function may indicate adaptation to variable nutrient supply in soils of different climatic environments (Liu et al., 2020b; Geng et al., 2017). The timing of flowering is a key diversification trait that determines the adaptation of soybean populations to different environments depending on their photoperiodic sensitivity (Li et al., 2020). A function of the candidate genes in abiotic stress responses has been suggested by differences in their expression in different experiments under stress conditions such as drought, salt, dehydration and cold, as well as by functional studies of their role in abiotic stress response cascades (e.g. Figure 6C and Table 3). In total, 6,314 genes were detected by our selection scans and may play a role in local adaptation. Many of these were characterised both by high genetic differentiation and a significant association with an environmental variable (Figure 3 E-G, Supplementary Data Worksheet 6). Such selection signals can be considered robust (Hoban et al., 2016) and genes located in these regions may be prioritized for further functional studies or marker-assisted selection in breeding programs.

For example, soybean breeding programs in Canada and Central Europe require adaptation to high latitude cold regions and may benefit most from selection signatures in scenario A because it included germplasm of the northernmost origin in China and Russia (Table 2).

Selection scans are susceptible to confounding factors that mimic selection signatures, leading to false positives, or mask selection signals and cause false negatives. Confounding factors include population structure, demographic history, lack of genomic and environmental information, and statistical limitations (Hoban et al., 2016). In the following, we therefore discuss the extent to which these factors may have influenced our results and highlight current limitations and possible solutions for functional validation of selection signatures in soybean adaptation.

### Geographic Origin of Genebank Germplasm

The application of landscape genomics to domesticated crops faces the challenge that the germplasm used for analysis often originates from *ex situ* collections. These collections were established during or after the Green Revolution to preserve landraces that were replaced by modern cultivars. Collection expeditions at that time focused primarily on obtaining germplasm and neglected the collection of environmental and geographic parameters, which are required to understand the environmental condition that influenced the evolution of landraces (Curry, 2017). This lack of information also applies to the USDA soybean genebank collection (Nelson, 2011). Material from this collection was used in a previous landscape genomics analysis of landraces from China, Korea and Japan (Bandillo et al., 2017). Our approach differs from this study in two important ways. First, we excluded accessions from Korea and Japan as they represent different germplasm groups (Figure 1A-C). These groups may be influenced by geographical and ethnocultural boundaries or reflect independent domestication events that confound selection scans. Therefore, we focused on germplasm groups from China, which we considered sufficient to identify signatures of local adaptations due to the broad environmental range in which soybean is grown in China (Figures 1D and S1). Secondly, we did not rely on the georeference information contained in the passport data, as it is often inaccurate or accessions were grouped into regional provenances with imprecise geographic coordinates (Haupt and Schmid, 2020). In addition, geo-references were available for only a fraction of the accessions in the USDA soybean germplasm collection. To avoid bias due to misallocation and uneven representation, we grouped genebank accessions into groups that reflected their likely regional origin based on their population structure, classification into maturity groups and provincial origin. Therefore, we did not examine individual accessions but differences in allele frequencies of genetically homogeneous groups without admixed genotypes to infer genetic differentiation and genotype-environment associations. We assumed that population comparisons would allow both robust and sensitive landscape genomic analysis. It should be noted that the exclusion of admixed individuals reflects a trade-off between a comprehensive representation of the germplasm and the reliability of their geographical origin, which may inflate genomic differences between different germplasm groups if they were geographically evenly distributed and adaptive mixing of neighbouring groups occurred in ecological transition zones (Bandillo et al., 2017, 2015; Li et al., 2008). An extension of current passport information, e.g. by incorporating information from the Chinese soybean germplasm collection (Qiu et al., 2011), could allow the future inclusion of admixed germplasm if genotyping allows linkage of germplasm from different collections and subsequent combination of passport information. In addition, new collections from core areas where landraces are still cultivated may lead to high quality georeferenced collections to anchor existing accessions from previous collections.

### Selective Forces Shaping Soybean Germplasm

To investigate selective forces that have shaped genetic diversity in soybean-growing regions of China we used EARTHSTAT (Ray et al., 2012) and WorldClim (Fick and Hijmans, 2017) datasets and assumed that they adequately represented the geographical extent and environmental conditions at the time of soybean landrace adaptation. We considered a limited number of environmental parameters (Table 2) because the available data were based on only three environmental qualities: temperature, precipitation and solar radiation. Therefore, our environmental characterisation is not complete, but considering previous research on soybean ecotypes (Song et al., 2016; Wang and Gai, 2002), it can be assumed that our approximation has adequately captured the main selective gradients. Since we focused on comparisons between germplasm groups with large-scale provenances within groups (Figure 1D), the spatial resolution of environmental data was adequate. The temporal resolution was limited to monthly averages, which excludes the detection of selective forces such as sudden spring temperature drops or other shortterm extreme weather conditions that are not captured by monthly summaries (Hoban et al., 2016; Rellstab et al., 2015). The climate data exhibited a high degree of multicollinearity, making it difficult to identify the true causes of selection among measured (and unmeasured) variables. To account for multicollinearity, we combined environmental descriptors in a multivariate principal components analysis and used only the first two principal components, which together explained 97.6% of the environmental variation, as covariates for genotype-environment associations (Table 2). Despite these limitations we obtained significant genotype-environment associations for about 50% of genomic regions with strong genetic differentiation (high *XtX* values) between germplasm groups (Figure 3E-G). This result is very promising for future studies to identify causal genes for local adaptation in these genomic regions.

In addition to natural selection, native soybean germplasm was also subjected to multilayered artificial selection after domestication, which included aesthetic and culinary preferences such as colour, texture and flavour, as well as agronomic traits that were determined by local cultivation practices, crop rotations and traditional uses. These factors are difficult to capture as they are missing from the passport information, although they have influenced the geographical distribution of locally conserved landraces as well as their genomic diversity. It remains a challenge to distinguish selection signatures caused by local environmental adaptation from agronomic selecton, which requires a cautious interpretation of our results.

### Population Structure and Demography

The high average genetic differentiation between germplasm groups increases the variance of differentiation statistics like *XtX* and only loci with large effects caused by selection are detected as significant outliers in the distribution of *XtX* values or have high Bayes Factors in the environmental association analysis (Hoban et al., 2016; Hodgins and Yeaman, 2019). Our approach is therefore biased towards detecting loci with large effects with a high false-negative rate whereas loci with small and medium effects on local adaptation likely remain undetected. Correspondingly, not all genes previously know to play a role in adaptation loci were identified by our selection tests (Figure 3A-D). In addition, neutral population structure and demographic events such as migration, dispersal, and population bottlenecks may generate signals that mimic selection and inflate selection scans with false positives. For a robust differentiation between selection and neutrality, we used BAYPASS utility functions and the estimated covariance of allele frequencies between populations to generate a null distribution of *XtX* statistics and reduce the proportion of false positives (Gautier, 2015). Alternatively, the demographic history of soybean in China could have been modeled and used to simulate the *XtX* statistic. However, this approach is highly susceptible to deviations from the true demographic history (Hoban et al., 2016), which likely is complex for soybean due to human influence, repeated introgression of wild *G. soja*, and multiple domestication events (Sedivy et al., 2017).

### Genetic Architecture of Environmental Adaptation

It is frequently assumed that locally adaptive phenotypes are controlled by many genes with small effects (Hoban et al., 2016). Adaptation may predominantly result from small and correlated allele frequency changes at many loci (Stetter et al., 2018). Under such a polygenic mode of adaptation, most loci involved are expected to lack signatures of strong selection and are not identified in selection scans. As noted above, several genes previously reported in the context of environmental adaptation in modern soybean populations showed increased differentiation and/or environmental association but did not exceed significance thresholds (Figure 3A-D), consistent with the hypothesis of polygenic adaptation. Allelic heterogeneity (multiple adaptive alleles at a locus), conditional neutrality, pleiotropy and epistasis also may mask signals of adaptation in selection scans (Hoban et al., 2016). The ability to identify adaptive variation in soybean may be reduced by a high level of self-fertilization, because simulations suggest that such a mode of reproduction leads to genetic adaptation by multiple genes that are dispersed throughout the genome and have smaller average effect sizes (Hodgins and Yeaman, 2019). Nevertheless, our study revealed that many distinct genomic regions include genes previously implicated in soybean adaptation (Figure 3A-D) and selection signatures were enriched for gene functions like flowering, photoperiodism, and responses to abiotic and biotic stresses (Table S1). Some extended genomic regions may represent adaptation hotspots because they combine multiple loci with small effects (e.g. Figure 4B-D), although long-range signatures of selection may also be caused by inversions that suppress recombination locally and may be neutral. Inversions frequently play a role in adaptation if they preserve favourable allele combinations (Huang and Rieseberg, 2020). Several regions with selection signatures formed LD networks despite being located on different chromosomes and may indicate epistatic fitness interactions (Figures 6B, S48 and S49). One network may be involved in regulation of flowering (Figure 5), but the functional involvement of the other networks remains unknown because numerous genes are located in these regions which makes it difficult to differentiate targets of selection and linked genes. Since gene ontology enrichment analysis also did not detect significantly enriched GO terms for these networks, they may consist of artefacts resulting from population structure or population mixing (Kemppainen et al., 2015), i.e. introgressions of alleles from wild *G. soja* populations into East Asian landrace populations (Sedivy et al., 2017) that may have contributed to local adaptation during soybean domestication and expansion. Epistatic interactions can also be detected in the form of gene coexpression clusters (Blanc et al., 2020), and we therefore conducted a coexpression analysis with published transcriptome data of 61 US milestone varieties to examine co-occurrence of gene expression variation and LD patterns. Although coexpression networks did not reflect LD networks (Figure 6), the hypothesis of long-range epistatic interactions should not be rejected at this stage. Future analyses should include transcriptome data of native germplasm from China, which are becoming available (Liu et al., 2020b) and will allow population-specific expression studies of adaptation genes.

### Future perspectives for functional analysis and breeding

Our identification of selection signatures was based on genetic variation at the SNP level. The soybean haplotype map resource, GmHapMap, (Torkamaneh et al., 2021) provided a high marker density and allowed the exploration of genetic differentiation and association signals at gene-level resolution, thereby exceeding the capability of the underlying SoySNP50K genotypes (Song et al., 2015). In addition to SNPs, also structural variants such as copy number variations or inversions are recognized to be critical agents of adaptation (Wellenreuther et al., 2019). Since long range genomic regions with selection signatures and high local LD (Figure 2, S48 and S49) may resemble inversions, such patterns need to be validated using long-read sequencing technology. Recent advances in soybean pan-genome genomics (Liu et al., 2020b) revealed an enrichment of abiotic and biotic response genes in the dispensable part of the soybean genome, indicating the importance that presence-absence variation may play for local adaptation. Therefore, it is important to note that our analysis is currently limited to adaptive variation represented in SNP genotypes and short read sequencing data and does not allow the identification of more complex types of adaptive variants. However, recent genome sequencing efforts of wild and landrace germplasm collections from East Asia (Sedivy et al., 2017) and new reference genomes (Bayer et al., 2021; Liu et al., 2020b) will result in a high quality pangenome that makes the full spectrum of genetic variation availble for more precise landscape genomics analyses.

Despite the limitations of our genomic data we identified numerous candidate genes for soybean local adaptation, which now require functional validation. Validation should include comparative genomics to select the most promising candidate genes based on evolutionary conservation and predicted phenotypic effects. Candidate genes can then be genome edited to produce knock-out or knock-in mutations, or modify gene product dosage by targeting regulatory elements (Fernandez i Marti and Dodd, 2018). For example, multiplexed CRISPR/Cas9-mediated knock-outs of four functional homologs of the *Arabidopsis thaliana* gene *AtLNK2* in soybean accelerated flowering under long-day conditions (Li et al., 2021), demonstrating the feasibility of adaptive modification in soybean for experimental validation and practical application. Bai et al. (2020) reported on a CRISPR/Cas9 platform targeting a total of 102 genes and the creation of a soybean mutagenesis population with successful multiplexing of up to eight target genes in one plant. Similar populations in which putative adaptation loci are mutated will allow their functional characterization and the assessment of their value for crop improvement.

Given that adaptive traits are likely polygenic, marker-assisted selection of progeny from crosses among modern varieties that complement each other in haplotype blocks with putative adaptive alleles can be used to enrich genomes with favorable alleles for cultivation in new target production environments in cooler and higher latitudes in North America and Europe. Such a genomics-based allele stacking approach among modern cultivars can be accommodated in elite breeding programs while reducing linkage drag that is frequently observed during the introgression of exotic genetic resources. The integration of landscape genomic approaches into breeding programs using genomic prediction of adaptive phenotypes provides an attractive avenue for adapting crops to future climates.

## Acknowledgements

The authors thank the United States Department of Agriculture (USDA) and the USDA Soybean Germplasm Collection for their open data and free distribution of germplasm policies that made this work possible. This work was funded by BMEL Project Sojagenopath (FKZ 2814EPS012). Seeds of European soybean varieties were provided by the State Breeding Institute, University of Hohenheim.

## Author Contribution Statement

MH and KS designed the study, MH performed the analyses and wrote the first version of the manuscript. MH and KS wrote the final version of the manuscript.

## Data Accessibility

SoySNP50K genotype data for 160 modern European soybean varieties are available at http://doi.org/10.5281/zenodo.6126368. SoySNP50K genotypes of the USDA Soybean Germplasm Collection (Song et al., 2015, 2013) are available on SoyBase (https://soybase.org/snps/). GmHapMap genotypes of the USDA Soybean Germplasm Collection (Torkamaneh et al., 2021) are available on *figshare* (https://figshare.com/articles/dataset/USDA_20K_samples_Imputed_from_GmHapMap/7388750). Supplementary Data that include ancestry proportions inferred with ADMIXTURE and selected passport information, allele counts for soybean populations from China as inputs for BAYPASS, mean and standard deviation of marker specific *XtX* estimates, Bayes factors of environmental associations and a list of all candidate genes within genomic regions with signatures of selection are available at http://doi.org/10.5281/zenodo.6126368.

