## Supplementary Tables and Figures for "Using landscape genomics to infer genomic regions involved in environmental adaptation of soybean genebank accessions"

Max Haupt and Karl Schmid
University of Hohenheim, Stuttgart, Germany

#### Contents

**1 Supplementary Tables** **2**

**2 Supplementary Figures** **8**

### 1 Supplementary Tables

**Table S 1:** Results of the Gene Ontology Enrichment Analysis conducted with ShinyGO v0.61 (http://bioinformatics.sdstate.edu/go/) for genes located in selection signatures.

| Signal Type and Scenario | Enrichment FDR | Genes in list | Total genes | Functional Category |
|---|---|---|---|---|
| XtX A | 3.00E-03 | 18 | 127 | Response to reactive oxygen species |
| | 2.60E-02 | 71 | 1125 | Cellular component assembly |
| | 2.60E-02 | 11 | 69 | Response to hydrogen peroxide |
| | 2.60E-02 | 18 | 164 | Response to osmotic stress |
| | 2.60E-02 | 10 | 60 | Protein complex oligomerization |
| | 3.20E-02 | 17 | 157 | Response to salt stress |
| | 3.20E-02 | 6 | 23 | Phloem development |
| | 3.20E-02 | 4 | 9 | Long-day photoperiodism |
| | 3.20E-02 | 4 | 9 | Long-day photoperiodism, flowering |
| | 3.20E-02 | 4 | 9 | Regulation of long-day photoperiodism, flowering |
| | 4.00E-02 | 56 | 888 | Protein-containing complex assembly |
| $BF_{PC1}$ A | 2.30E-02 | 6 | 40 | Jasmonic acid metabolic process |
| | 2.30E-02 | 6 | 35 | Salicylic acid metabolic process |
| | 2.30E-02 | 5 | 23 | Phloem development |
| | 2.30E-02 | 6 | 41 | Phenol-containing compound metabolic process |
| | 2.90E-02 | 7 | 63 | Benzene-containing compound metabolic process |
| | 3.00E-02 | 6 | 47 | Phloem or xylem histogenesis |
| | 3.00E-02 | 61 | 2059 | Cellular catabolic process |
| | 3.00E-02 | 63 | 2140 | Organic substance catabolic process |
| | 3.00E-02 | 7 | 67 | Aspartate family amino acid biosynthetic process |
| | 3.00E-02 | 2 | 2 | Gamma-aminobutyric acid catabolic process |
| | 3.20E-02 | 6 | 53 | Stomatal movement |
| | 3.20E-02 | 6 | 53 | Regulation of stomatal movement |
| | 3.40E-02 | 5 | 36 | Methionine biosynthetic process |
| $BF_{PC2}$ A | 2.70E-04 | 9 | 36 | Ionotropic glutamate receptor signaling pathway |
| | 2.70E-04 | 9 | 36 | Glutamate receptor signaling pathway |
| | 4.50E-02 | 10 | 87 | Monosaccharide transmembrane transport |
| | 4.50E-02 | 10 | 87 | Hexose transmembrane transport |
| XtX and $BF_{PC1}$ A | 8.10E-04 | 5 | 23 | Phloem development |
| | 6.40E-03 | 29 | 1385 | Macromolecule catabolic process |
| | 1.00E-02 | 5 | 47 | Phloem or xylem histogenesis |
| | 1.30E-02 | 9 | 211 | RNA phosphodiester bond hydrolysis, endonucleolytic |
| | 1.30E-02 | 37 | 2140 | Organic substance catabolic process |
| | 1.60E-02 | 4 | 35 | Salicylic acid metabolic process |
| | 1.60E-02 | 35 | 2059 | Cellular catabolic process |
| | 1.60E-02 | 5 | 60 | Protein complex oligomerization |
| | 2.00E-02 | 4 | 40 | Jasmonic acid metabolic process |
| | 2.00E-02 | 4 | 41 | Phenol-containing compound metabolic process |
| | 2.00E-02 | 5 | 69 | Response to hydrogen peroxide |
| | 2.00E-02 | 7 | 155 | RNA catabolic process |
| | 2.00E-02 | 39 | 2450 | Catabolic process |
| | 2.00E-02 | 7 | 157 | Response to salt stress |
| | 2.40E-02 | 7 | 164 | Response to osmotic stress |
| | 2.70E-02 | 9 | 274 | Polysaccharide catabolic process |
| | 3.00E-02 | 9 | 282 | RNA phosphodiester bond hydrolysis |
| | 3.00E-02 | 6 | 127 | Response to reactive oxygen species |
| | 3.10E-02 | 15 | 670 | Polysaccharide metabolic process |
| | 4.90E-02 | 9 | 308 | Organic hydroxy compound metabolic process |
| XtX and $BF_{PC2}$ A | 2.00E-03 | 5 | 36 | Ionotropic glutamate receptor signaling pathway |
| | 2.00E-03 | 5 | 36 | Glutamate receptor signaling pathway |
| | 7.30E-03 | 6 | 87 | Monosaccharide transmembrane transport |
| | 7.30E-03 | 6 | 87 | Hexose transmembrane transport |
| XtX B | 2.10E-05 | 13 | 48 | Systemic acquired resistance |
| | 2.10E-05 | 13 | 50 | Defense response, incompatible interaction |
| | 8.50E-03 | 15 | 115 | Immune system process |
| | 8.50E-03 | 15 | 113 | Innate immune response |
| | 8.50E-03 | 15 | 113 | Immune response |
| | 3.00E-02 | 8 | 41 | Galactose metabolic process |
| $BF_{PC1}$ B | 3.00E-04 | 6 | 40 | Jasmonic acid metabolic process |
| | 3.00E-04 | 6 | 35 | Salicylic acid metabolic process |
| | 3.00E-04 | 6 | 41 | Phenol-containing compound metabolic process |
| | 2.90E-03 | 6 | 63 | Benzene-containing compound metabolic process |
| $BF_{PC2}$ B | 1.60E-02 | 114 | 2450 | Catabolic process |
| XtX and $BF_{PC1}$ B | 3.00E-02 | 3 | 40 | Jasmonic acid metabolic process |
| | 3.00E-02 | 3 | 35 | Salicylic acid metabolic process |
| | 3.00E-02 | 3 | 41 | Phenol-containing compound metabolic process |
| | 4.50E-02 | 31 | 4978 | Protein modification process |
| | 4.50E-02 | 3 | 63 | Benzene-containing compound metabolic process |
| | 4.50E-02 | 2 | 15 | Chloroplast rRNA processing |
| | 4.50E-02 | 31 | 4978 | Cellular protein modification process |
| XtX and $BF_{PC2}$ B | 1.10E-02 | 62 | 2140 | Organic substance catabolic process |
| | 1.10E-02 | 68 | 2450 | Catabolic process |
| | 1.60E-02 | 6 | 41 | Galactose metabolic process |

**Table S 2:** Results of the Gene Ontology Enrichment Analysis conducted with ShinyGO v0.61 (http://bioinformatics.sdstate.edu/go/) for genes in co-expression modules.

| Module | Genes in Module | Genes in Signals | Enrichment FDR | Genes in list | Total genes | Functional Category |
|---|---|---|---|---|---|---|
| M1 | 941 | 98 | 0.024 | 9 | 103 | Phenylpropanoid metabolic process |
| | | | 0.024 | 7 | 67 | Lignin metabolic process |
| | | | 0.024 | 6 | 47 | Phenylpropanoid catabolic process |
| | | | 0.024 | 6 | 47 | Lignin catabolic process |
| M2 | 647 | 55 | 3.8E-07 | 12 | 100 | Chromatin assembly |
| | | | 3.8E-07 | 13 | 117 | DNA packaging |
| | | | 3.8E-07 | 12 | 98 | Nucleosome assembly |
| | | | 7E-07 | 12 | 108 | Chromatin assembly or disassembly |
| | | | 1E-06 | 12 | 114 | Nucleosome organization |
| | | | 4.5E-06 | 14 | 188 | Protein-DNA complex assembly |
| | | | 1.5E-05 | 14 | 210 | Protein-DNA complex subunit organization |
| | | | 0.013 | 16 | 482 | DNA conformation change |
| | | | 0.018 | 16 | 502 | Chromatin organization |
| | | | 0.048 | 24 | 1027 | Chromosome organization |
| | | | 0.048 | 5 | 65 | Response to water deprivation |
| | | | 0.048 | 5 | 67 | Response to water |
| | | | 0.049 | 15 | 543 | Cell cycle process |
| | | | 0.049 | 2 | 6 | DGTP metabolic process |
| | | | 0.049 | 2 | 6 | Purine nucleotide catabolic process |
| | | | 0.049 | 2 | 6 | DGTP catabolic process |
| | | | 0.049 | 2 | 6 | Purine nucleoside triphosphate catabolic process |
| | | | 0.049 | 2 | 6 | Purine deoxyribonucleotide metabolic process |
| | | | 0.049 | 2 | 6 | Purine deoxyribonucleotide catabolic process |
| | | | 0.049 | 2 | 6 | Purine deoxyribonucleoside triphosphate metabolic process |
| | | | 0.049 | 2 | 6 | Purine deoxyribonucleoside triphosphate catabolic process |
| M3 | 598 | 60 | 1.2E-05 | 6 | 23 | Removal of superoxide radicals |
| | | | 1.2E-05 | 6 | 23 | Cellular response to oxygen radical |
| | | | 1.2E-05 | 6 | 23 | Cellular response to superoxide |
| | | | 1.2E-05 | 6 | 23 | Response to superoxide |
| | | | 1.2E-05 | 6 | 23 | Response to oxygen radical |
| | | | 1.2E-05 | 6 | 23 | Superoxide metabolic process |
| | | | 0.00038 | 6 | 41 | Cellular response to reactive oxygen species |
| | | | 0.0047 | 4 | 21 | Cutin biosynthetic process |
| | | | 0.015 | 6 | 81 | Cellular response to oxidative stress |
| | | | 0.026 | 7 | 127 | Response to reactive oxygen species |
| | | | 0.031 | 3 | 16 | TRNA 5 -leader removal |
| M4 | 441 | 47 | 0.00081 | 12 | 270 | Response to biotic stimulus |
| | | | 0.0037 | 5 | 43 | Regulation of protein serine/threonine phosphatase activity |
| | | | 0.0037 | 3 | 8 | Asparagine biosynthetic process |
| | | | 0.0037 | 5 | 53 | Negative regulation of phosphatase activity |
| | | | 0.0037 | 5 | 52 | Negative regulation of phosphoprotein phosphatase activity |
| | | | 0.0037 | 5 | 53 | Negative regulation of dephosphorylation |
| | | | 0.0037 | 5 | 52 | Negative regulation of protein dephosphorylation |
| | | | 0.0047 | 4 | 31 | Plant-type ovary development |
| | | | 0.0047 | 4 | 31 | Plant ovule development |
| | | | 0.0049 | 4 | 33 | Carpel development |
| | | | 0.0049 | 4 | 33 | Gynoecium development |
| | | | 0.0052 | 12 | 420 | Defense response |
| | | | 0.0053 | 3 | 14 | Asparagine metabolic process |
| | | | 0.0053 | 8 | 196 | Defense response to other organism |
| | | | 0.0085 | 5 | 74 | Negative regulation of protein modification process |
| | | | 0.01 | 5 | 80 | Negative regulation of phosphorus metabolic process |
| | | | 0.01 | 8 | 226 | Response to external biotic stimulus |
| | | | 0.01 | 5 | 80 | Negative regulation of phosphate metabolic process |
| | | | 0.01 | 8 | 226 | Response to other organism |
| | | | 0.01 | 4 | 47 | Floral whorl development |
| | | | 0.01 | 4 | 47 | Defense response to fungus |
| | | | 0.012 | 5 | 86 | Aspartate family amino acid metabolic process |
| | | | 0.014 | 5 | 90 | Glutathione metabolic process |
| | | | 0.014 | 5 | 92 | Regulation of protein dephosphorylation |
| | | | 0.014 | 5 | 92 | Regulation of phosphoprotein phosphatase activity |
| | | | 0.015 | 5 | 95 | Regulation of phosphatase activity |
| | | | 0.016 | 7 | 200 | Response to abscisic acid |
| | | | 0.016 | 5 | 97 | Regulation of dephosphorylation |
| | | | 0.016 | 7 | 200 | Response to alcohol |
| | | | 0.018 | 6 | 150 | Negative regulation of hydrolase activity |
| M5 | 428 | 38 | 4.5E-09 | 10 | 60 | Jasmonic acid mediated signaling pathway |
| | | | 4.5E-09 | 10 | 60 | Cellular response to jasmonic acid stimulus |
| | | | 4.5E-09 | 9 | 40 | Regulation of jasmonic acid mediated signaling pathway |
| | | | 1.4E-08 | 10 | 69 | Response to jasmonic acid |
| | | | 8E-07 | 8 | 55 | Response to wounding |
| | | | 6E-05 | 13 | 306 | Cellular response to acid chemical |
| | | | 6E-05 | 8 | 99 | Regulation of defense response |
| | | | 9.4E-05 | 11 | 233 | Regulation of signal transduction |
| | | | 9.4E-05 | 11 | 234 | Regulation of cell communication |
| | | | 9.4E-05 | 11 | 233 | Regulation of signaling |
| | | | 0.00012 | 15 | 459 | Response to acid chemical |
| | | | 0.00019 | 14 | 420 | Defense response |
| | | | 0.00087 | 10 | 251 | Cell wall biogenesis |
| | | | 0.00093 | 13 | 428 | Cellular response to oxygen-containing compound |

| Module | Genes in Module | Genes in Signals | Enrichment FDR | Genes in list | Total genes | Functional Category |
|---|---|---|---|---|---|---|
| | | | 0.0012 | 8 | 164 | Regulation of response to stress |
| | | | 0.002 | 19 | 905 | Response to hormone |
| | | | 0.002 | 13 | 473 | Hormone-mediated signaling pathway |
| | | | 0.002 | 32 | 2002 | Signal transduction |
| | | | 0.002 | 32 | 2010 | Signaling |
| | | | 0.0021 | 19 | 923 | Response to endogenous stimulus |
| | | | 0.0021 | 6 | 97 | Xyloglucan metabolic process |
| | | | 0.0021 | 34 | 2210 | Cell communication |
| | | | 0.0026 | 11 | 367 | Regulation of response to stimulus |
| | | | 0.0034 | 7 | 155 | Hemicellulose metabolic process |
| | | | 0.0034 | 13 | 514 | Cellular response to hormone stimulus |
| | | | 0.0043 | 13 | 532 | Cellular response to endogenous stimulus |
| | | | 0.0045 | 3 | 18 | Diterpenoid catabolic process |
| | | | 0.0045 | 3 | 18 | Terpenoid catabolic process |
| | | | 0.0045 | 8 | 218 | Cell wall macromolecule metabolic process |
| | | | 0.0045 | 3 | 18 | Gibberellin catabolic process |
| M6 | 356 | 42 | 0.00055 | 5 | 32 | Response to chitin |
| | | | 0.0008 | 13 | 433 | Response to auxin |
| | | | 0.0046 | 20 | 1126 | Response to organic substance |
| M7 | 230 | 29 | 1.2E-46 | 27 | 60 | Protein complex oligomerization |
| | | | 1.2E-46 | 36 | 196 | Response to heat |
| | | | 4.9E-45 | 27 | 69 | Response to hydrogen peroxide |
| | | | 8E-43 | 36 | 251 | Response to temperature stimulus |
| | | | 2.7E-39 | 27 | 106 | Response to antibiotic |
| | | | 5E-39 | 38 | 386 | Protein folding |
| | | | 6E-39 | 28 | 127 | Response to reactive oxygen species |
| | | | 2E-34 | 27 | 157 | Response to salt stress |
| | | | 6.3E-34 | 27 | 164 | Response to osmotic stress |
| | | | 1.8E-29 | 29 | 302 | Response to inorganic substance |
| | | | 3.3E-27 | 36 | 697 | Response to abiotic stimulus |
| | | | 4.1E-24 | 28 | 419 | Response to toxic substance |
| | | | 7.2E-24 | 27 | 384 | Response to drug |
| | | | 9.1E-24 | 28 | 434 | Response to oxidative stress |
| | | | 7.5E-19 | 29 | 728 | Response to oxygen-containing compound |
| | | | 1.4E-16 | 29 | 888 | Protein-containing complex assembly |
| | | | 8.1E-15 | 30 | 1125 | Cellular component assembly |
| | | | 8.7E-15 | 29 | 1045 | Protein-containing complex subunit organization |
| | | | 5.4E-14 | 39 | 2127 | Response to stress |
| | | | 7.6E-12 | 34 | 1907 | Response to chemical |
| | | | 6.8E-11 | 31 | 1726 | Cellular component biogenesis |
| | | | 1.8E-09 | 9 | 93 | Cellular response to heat |
| | | | 2.8E-06 | 7 | 107 | Chaperone-mediated protein folding |
| | | | 3.3E-05 | 42 | 4891 | Response to stimulus |
| | | | 0.00016 | 5 | 80 | de novo posttranslational protein folding |
| | | | 0.00016 | 5 | 80 | Chaperone cofactor-dependent protein refolding |
| | | | 0.0002 | 4 | 42 | Cellular response to unfolded protein |
| | | | 0.00021 | 4 | 43 | Response to unfolded protein |
| | | | 0.00041 | 5 | 100 | de novo protein folding |
| | | | 0.0005 | 32 | 3758 | Cellular component organization |
| M8 | 214 | 18 | NA | | | No significant enrichment found |
| M9 | 194 | 21 | 0.016 | 2 | 6 | Triterpenoid biosynthetic process |
| | | | 0.016 | 2 | 6 | Triterpenoid metabolic process |
| M10 | 132 | 14 | 2.4E-06 | 5 | 30 | Response to cadmium ion |
| | | | 1.1E-05 | 6 | 95 | Response to metal ion |
| | | | 1.1E-05 | 6 | 97 | Xyloglucan metabolic process |
| | | | 1.1E-05 | 8 | 251 | Cell wall biogenesis |
| | | | 2.3E-05 | 3 | 7 | Inositol catabolic process |
| | | | 7.5E-05 | 6 | 155 | Hemicellulose metabolic process |
| | | | 7.5E-05 | 10 | 599 | Cellular carbohydrate metabolic process |
| | | | 0.00011 | 4 | 43 | Regulation of protein serine/threonine phosphatase activity |
| | | | 0.00013 | 6 | 177 | Cell wall polysaccharide metabolic process |
| | | | 0.00016 | 4 | 53 | Negative regulation of phosphatase activity |
| | | | 0.00016 | 4 | 52 | Negative regulation of phosphoprotein phosphatase activity |
| | | | 0.00016 | 4 | 53 | Negative regulation of dephosphorylation |
| | | | 0.00016 | 4 | 52 | Negative regulation of protein dephosphorylation |
| | | | 0.00027 | 6 | 218 | Cell wall macromolecule metabolic process |
| | | | 0.00044 | 3 | 24 | Polyol catabolic process |
| | | | 0.00049 | 4 | 74 | Negative regulation of protein modification process |
| | | | 0.00049 | 3 | 26 | Alcohol catabolic process |
| | | | 0.00056 | 4 | 80 | Negative regulation of phosphorus metabolic process |
| | | | 0.00056 | 4 | 80 | Negative regulation of phosphate metabolic process |
| | | | 0.00065 | 3 | 30 | Organic hydroxy compound catabolic process |
| | | | 0.00075 | 3 | 32 | Inositol metabolic process |
| | | | 0.00077 | 4 | 92 | Regulation of protein dephosphorylation |
| | | | 0.00077 | 4 | 92 | Regulation of phosphoprotein phosphatase activity |
| | | | 0.00077 | 6 | 290 | Cellular glucan metabolic process |
| | | | 0.00079 | 6 | 302 | Response to inorganic substance |
| | | | 0.00079 | 4 | 95 | Regulation of phosphatase activity |
| | | | 0.00079 | 4 | 97 | Regulation of dephosphorylation |
| | | | 0.00079 | 6 | 300 | Glucan metabolic process |
| | | | 0.00079 | 9 | 748 | Cell wall organization or biogenesis |
| | | | 0.0014 | 7 | 485 | Carbohydrate catabolic process |
| M11 | 126 | 17 | 3.2E-05 | 8 | 266 | Cell redox homeostasis |
| | | | 9.3E-05 | 9 | 470 | Cellular homeostasis |
| | | | 9.3E-05 | 12 | 939 | Homeostatic process |
| | | | 0.0013 | 13 | 1474 | Regulation of biological quality |
| | | | 0.0014 | 7 | 405 | Electron transport chain |
| | | | 0.002 | 9 | 770 | Generation of precursor metabolites and energy |
| | | | 0.0035 | 2 | 8 | Positive regulation of reactive oxygen species metabolic process |

| Module | Genes in Module | Genes in Signals | Enrichment FDR | Genes in list | Total genes | Functional Category |
|---|---|---|---|---|---|---|
| | | | 0.011 | 2 | 15 | Regulation of reactive oxygen species metabolic process |
| | | | 0.013 | 2 | 18 | Cadmium ion transport |
| | | | 0.013 | 2 | 18 | Cadmium ion transmembrane transport |
| | | | 0.03 | 2 | 40 | Response to brassinosteroid |
| | | | 0.03 | 2 | 38 | Brassinosteroid mediated signaling pathway |
| | | | 0.03 | 2 | 31 | Photosynthesis |
| | | | 0.03 | 2 | 38 | Steroid hormone mediated signaling pathway |
| | | | 0.03 | 2 | 38 | Response to steroid hormone |
| | | | 0.03 | 3 | 137 | Metal ion homeostasis |
| | | | 0.03 | 16 | 3223 | Oxidation-reduction process |
| | | | 0.03 | 2 | 38 | Cellular response to brassinosteroid stimulus |
| | | | 0.03 | 2 | 38 | Cellular response to steroid hormone stimulus |
| | | | 0.03 | 2 | 39 | Manganese ion transmembrane transport |
| | | | 0.03 | 4 | 232 | Reactive oxygen species metabolic process |
| | | | 0.03 | 2 | 39 | Manganese ion transport |
| | | | 0.049 | 2 | 53 | Photosynthesis |
| M12 | 121 | 13 | 1.1E-13 | 8 | 29 | Sequestering of metal ion |
| | | | 1.1E-13 | 8 | 29 | Sequestering of iron ion |
| | | | 1.1E-13 | 8 | 29 | Intracellular sequestering of iron ion |
| | | | 1.2E-12 | 8 | 39 | Iron ion transport |
| | | | 2.1E-11 | 8 | 56 | Cellular iron ion homeostasis |
| | | | 6E-11 | 8 | 68 | Cellular transition metal ion homeostasis |
| | | | 6E-11 | 8 | 68 | Maintenance of location in cell |
| | | | 6E-11 | 8 | 67 | Iron ion homeostasis |
| | | | 6E-11 | 9 | 106 | Transition metal ion transport |
| | | | 2.6E-10 | 8 | 82 | Transition metal ion homeostasis |
| | | | 9.2E-10 | 8 | 97 | Maintenance of location |
| | | | 2.7E-09 | 9 | 170 | Cellular cation homeostasis |
| | | | 2.9E-09 | 8 | 114 | Cellular metal ion homeostasis |
| | | | 3.7E-09 | 9 | 179 | Cellular ion homeostasis |
| | | | 9.2E-09 | 9 | 200 | Cellular chemical homeostasis |
| | | | 1E-08 | 8 | 137 | Metal ion homeostasis |
| | | | 1.9E-08 | 9 | 222 | Cation homeostasis |
| | | | 1.9E-08 | 9 | 222 | Inorganic ion homeostasis |
| | | | 2.6E-08 | 9 | 231 | Ion homeostasis |
| | | | 7.2E-07 | 9 | 342 | Chemical homeostasis |
| | | | 8.1E-07 | 4 | 19 | Cellular manganese ion homeostasis |
| | | | 8.6E-07 | 10 | 470 | Cellular homeostasis |
| | | | 9.2E-07 | 4 | 20 | Manganese ion homeostasis |
| | | | 1.8E-06 | 10 | 515 | Metal ion transport |
| | | | 3E-06 | 4 | 27 | Iron ion transmembrane transport |
| | | | 6.1E-06 | 5 | 77 | Response to cytokinin |
| | | | 6.1E-06 | 15 | 1474 | Regulation of biological quality |
| | | | 7.9E-06 | 12 | 939 | Homeostatic process |
| | | | 1.2E-05 | 4 | 39 | Manganese ion transmembrane transport |
| | | | 1.2E-05 | 4 | 39 | Manganese ion transport |
| M13 | 99 | 6 | 7.7E-24 | 20 | 295 | Photosynthesis |
| | | | 2.2E-21 | 20 | 405 | Electron transport chain |
| | | | 4.9E-20 | 23 | 770 | Generation of precursor metabolites and energy |
| | | | 2.3E-15 | 33 | 3223 | Oxidation-reduction process |
| | | | 9.7E-14 | 11 | 146 | Photosynthesis |
| | | | 3.9E-13 | 11 | 168 | Cellular respiration |
| | | | 7.1E-13 | 9 | 81 | Respiratory electron transport chain |
| | | | 1.4E-12 | 12 | 265 | ATP metabolic process |
| | | | 2.3E-12 | 11 | 207 | Energy derivation by oxidation of organic compounds |
| | | | 2.3E-12 | 12 | 281 | Purine ribonucleoside triphosphate metabolic process |
| | | | 2.7E-12 | 12 | 287 | Purine nucleoside triphosphate metabolic process |
| | | | 3.7E-12 | 12 | 297 | Ribonucleoside triphosphate metabolic process |
| | | | 4E-12 | 8 | 64 | ATP synthesis coupled electron transport |
| | | | 5.5E-12 | 12 | 313 | Purine nucleoside monophosphate metabolic process |
| | | | 5.5E-12 | 12 | 313 | Purine ribonucleoside monophosphate metabolic process |
| | | | 5.8E-12 | 8 | 69 | Oxidative phosphorylation |
| | | | 5.8E-12 | 12 | 318 | Nucleoside triphosphate metabolic process |
| | | | 1.1E-11 | 12 | 338 | Ribonucleoside monophosphate metabolic process |
| | | | 3.9E-11 | 7 | 50 | Photosynthetic electron transport chain |
| | | | 4.5E-11 | 12 | 384 | Nucleoside monophosphate metabolic process |
| | | | 7.7E-11 | 12 | 404 | Purine ribonucleotide metabolic process |
| | | | 1.2E-10 | 12 | 421 | Purine nucleotide metabolic process |
| | | | 2.3E-10 | 12 | 447 | Ribonucleotide metabolic process |
| | | | 4.8E-10 | 12 | 479 | Purine-containing compound metabolic process |
| | | | 8.6E-10 | 12 | 506 | Ribose phosphate metabolic process |
| | | | 4.4E-09 | 4 | 8 | Photosynthetic electron transport in photosystem II |
| | | | 1E-08 | 12 | 634 | Nucleotide metabolic process |
| | | | 1.2E-08 | 12 | 647 | Nucleoside phosphate metabolic process |
| | | | 8.5E-08 | 14 | 1132 | Drug metabolic process |
| | | | 1.1E-07 | 12 | 791 | Nucleobase-containing small molecule metabolic process |
| M14 | 59 | 5 | NA | | | No significant enrichment found |

**Table S 3:** Proportions of selection signature peak markers and of random markers with interchromosomal LD that surpasses LD thresholds. Only interchromosomal LD estimates were considered. Peak marker refers to the SNP with the local maximum in the differentiation or association statistic in the respective selection signal.

| Scenario | LD threshold | All Signals | XtX | $BF_{PC1}$ | $BF_{PC2}$ | Random |
|---|---|---|---|---|---|---|
| A | LD > 0.21 | 0.138629205 | 0.184688471 | 0.254898803 | 0.100837277 | 2.82E-02 |
| A | LD > 0.63 | 0.001706657 | 0.004281441 | 0.001878993 | 0.0001456134 | 4.45E-05 |
| B | LD > 0.21 | 0.0504926335 | 0.108799295 | 0.03711448 | 0.0413032055 | 0.0283923241 |
| B | LD > 0.63 | 0.0009526912 | 0.001865575 | 0.001568217 | 0.0009458749 | 0.0001759548 |

#### 2 Supplementary Figures

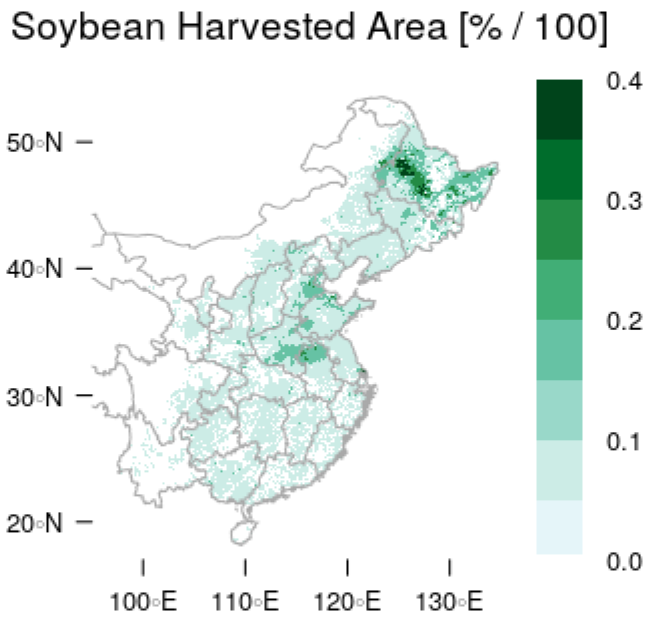


**Figure S 1:** Geographic extent of soybean cultivation in China (Ray et al. 2012).

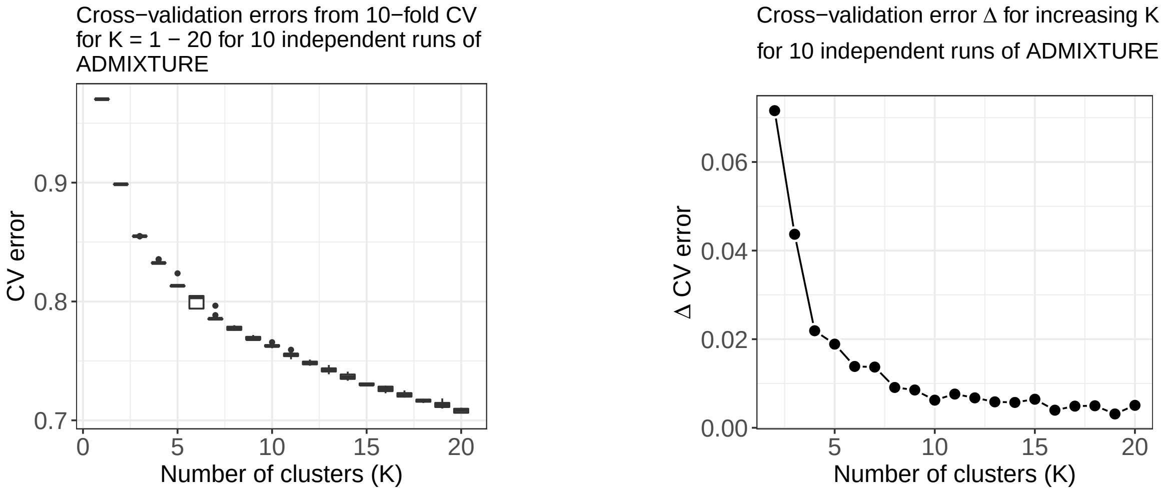


**Figure S 2:** Cross-validation error in the ADMIXTURE analysis for K varying from 1-20.

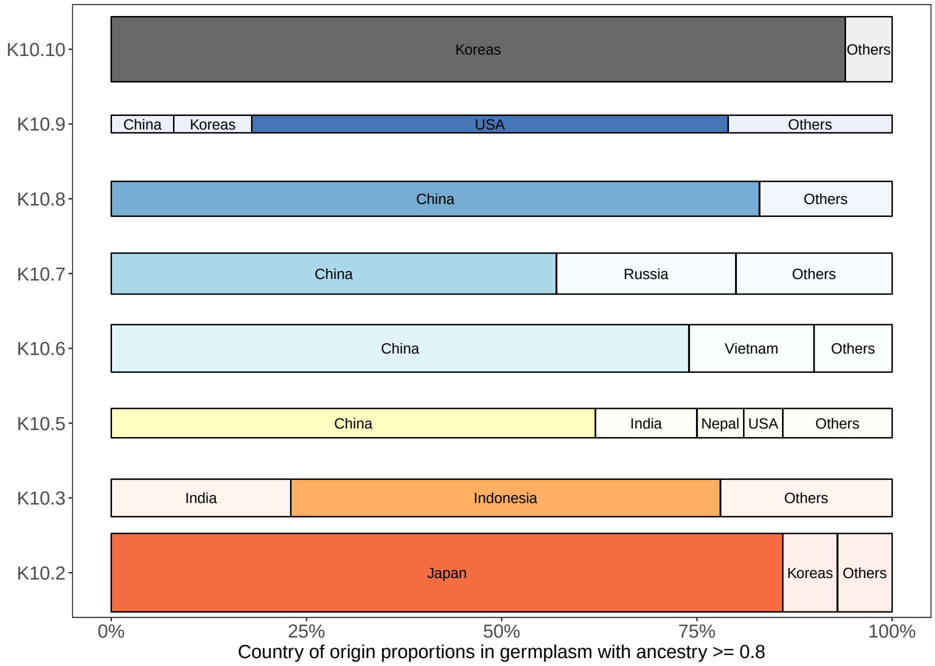


**Figure S 3:** Geographic origin of conserved accessions with ancestry fractions >= 80% with one of the inferred ancestral groups (K10.1 – K10.10) according to passport information. Bar width corresponds to group size.

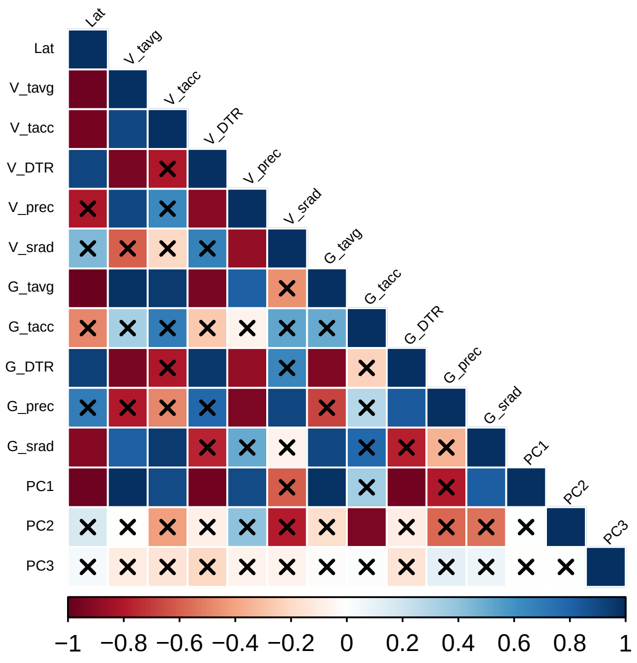


**Figure S 4:** Pearson correlation coefficients among environmental parameters characterizing six soybean growing regions in China. Black crosses indicate correlations that are not significant (p > 0.05).

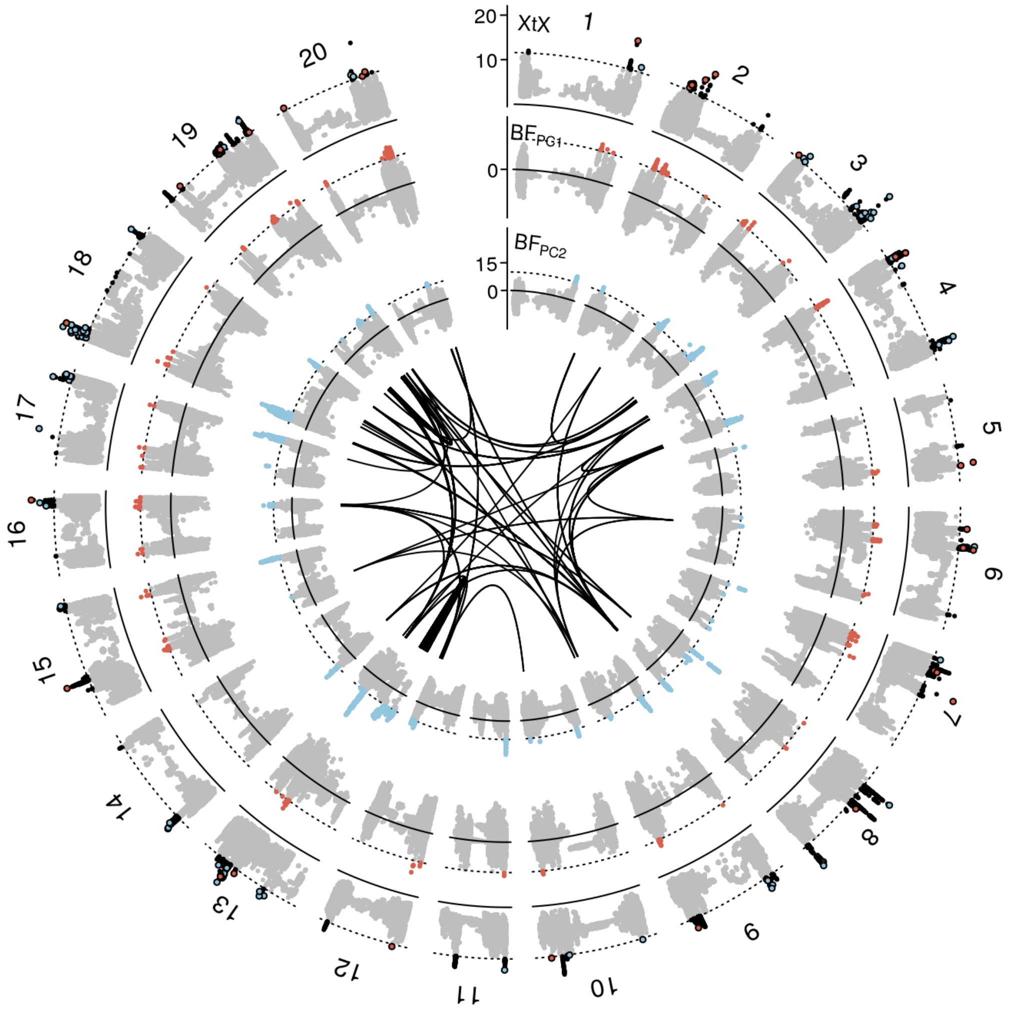


**Figure S 5:** Manhattan plot of the *XtX* statistic and of genotype-environment associations with the first two environmental principal components in Bayes factors for all 20 chromosomes in scenario B . Red and blue points in the *XtX* track represent overlaps between genetic differentiation and association signals with the first and second environmental principal components. The dotted horizontal lines represent the 1% POD significance threshold of the *XtX* statistic and the threshold of BF = 10 deciban. Black lines in the center represent regions with elevated LD exceeding the level of background LD 3-fold and a minimum physical distance of 5Mb between selection signatures.

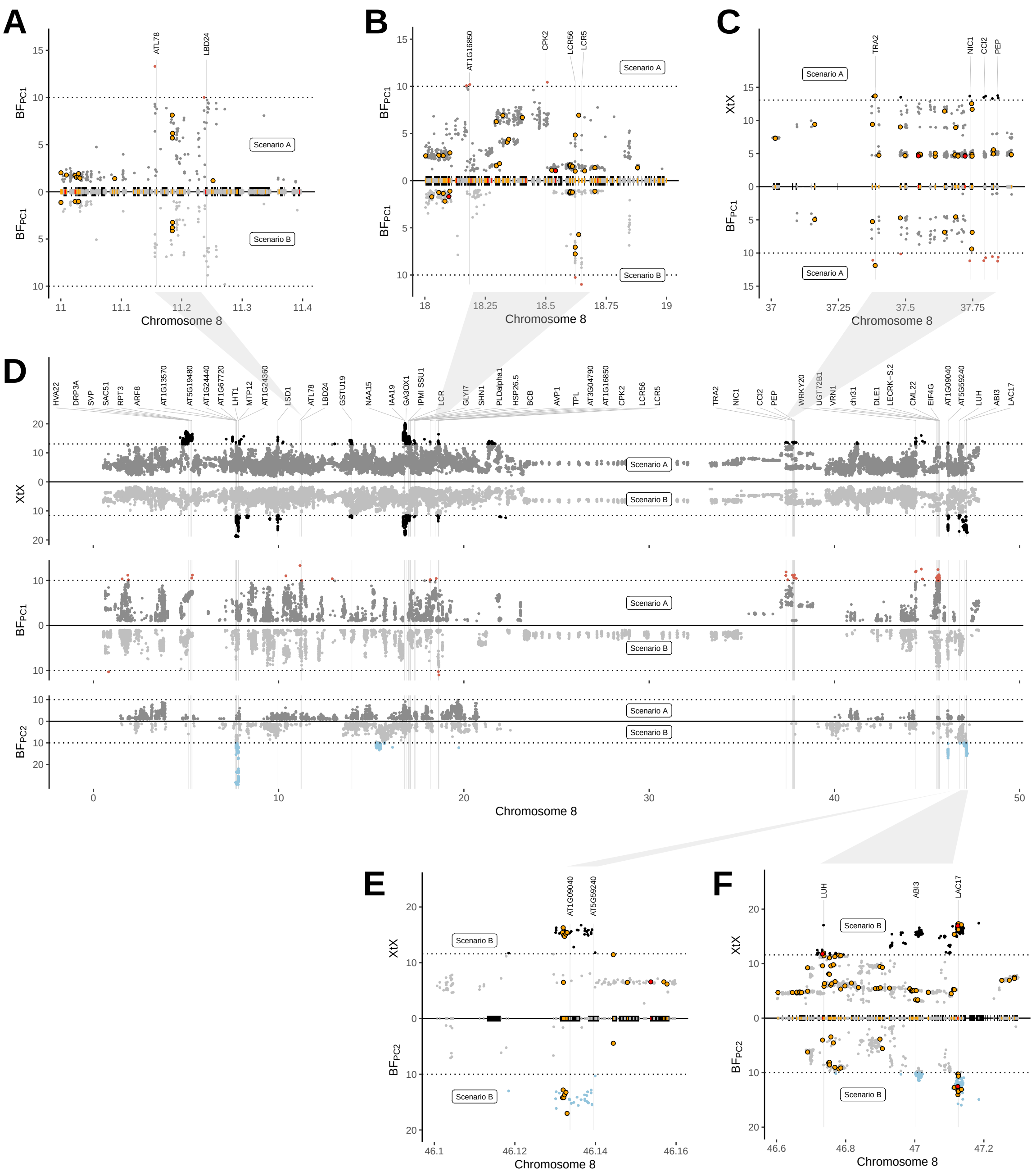

**Figure S 6:** Identification of candidate genes with a putative role in environmental adaptation on chromosome 8. (D) Manhattan plot of the *XtX* statistic and of genotype-environment associations with the first two environmental principal components in Bayes factors for scenario A and scenario B. Negative Bayes factors are omitted and dotted horizontal lines are analogous to Figure 2A. Labeled candidate genes are listed in Tab 3. Unit of x-axis is Mb. (A-C,E-F) Close-ups of genomic regions with selection signatures. Black blocks on the x-axis indicate the positions of predicted gene models. Red and yellow blocks and points indicate non-synonymous variants with high and moderate impacts on protein function.

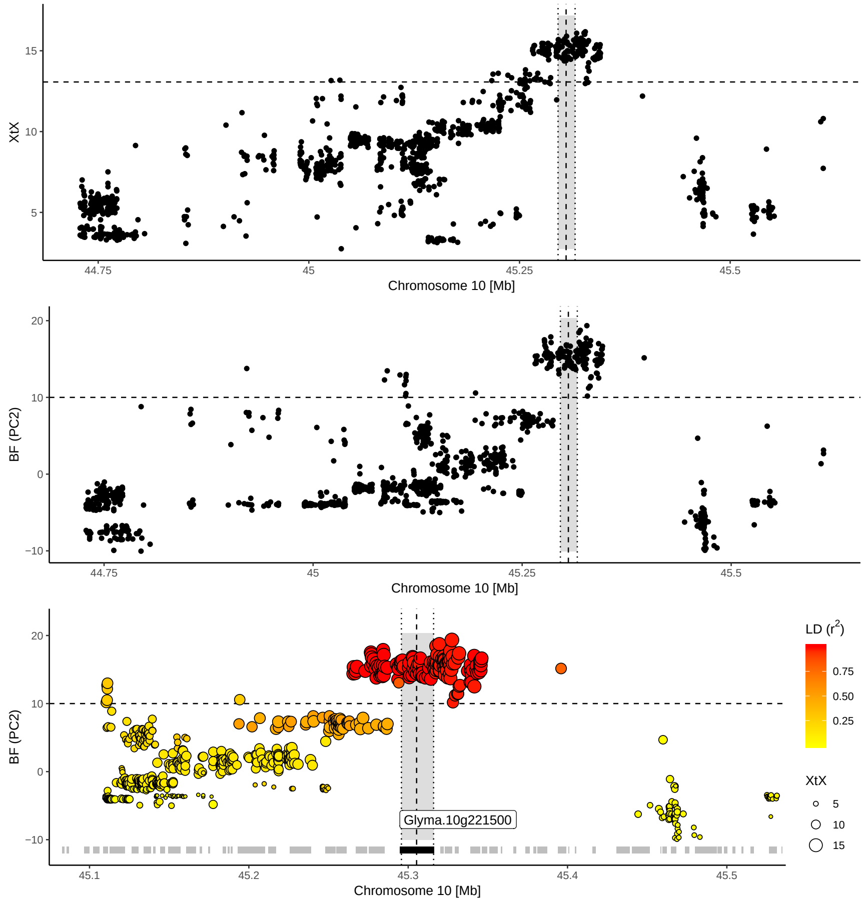


**Figure S 7:** Differentiation and association signature on chromosome 10 highlighting the region that harbors the *E2* locus (indicated by vertical dotted line) observed in scenario A. Bottom: Close-up of the region. Grey blocks on the x-axis indicate the position of predicted gene models, black block indicates the *E2* locus. Pairwise LD is reported with respect to a central marker in the *E2* locus.

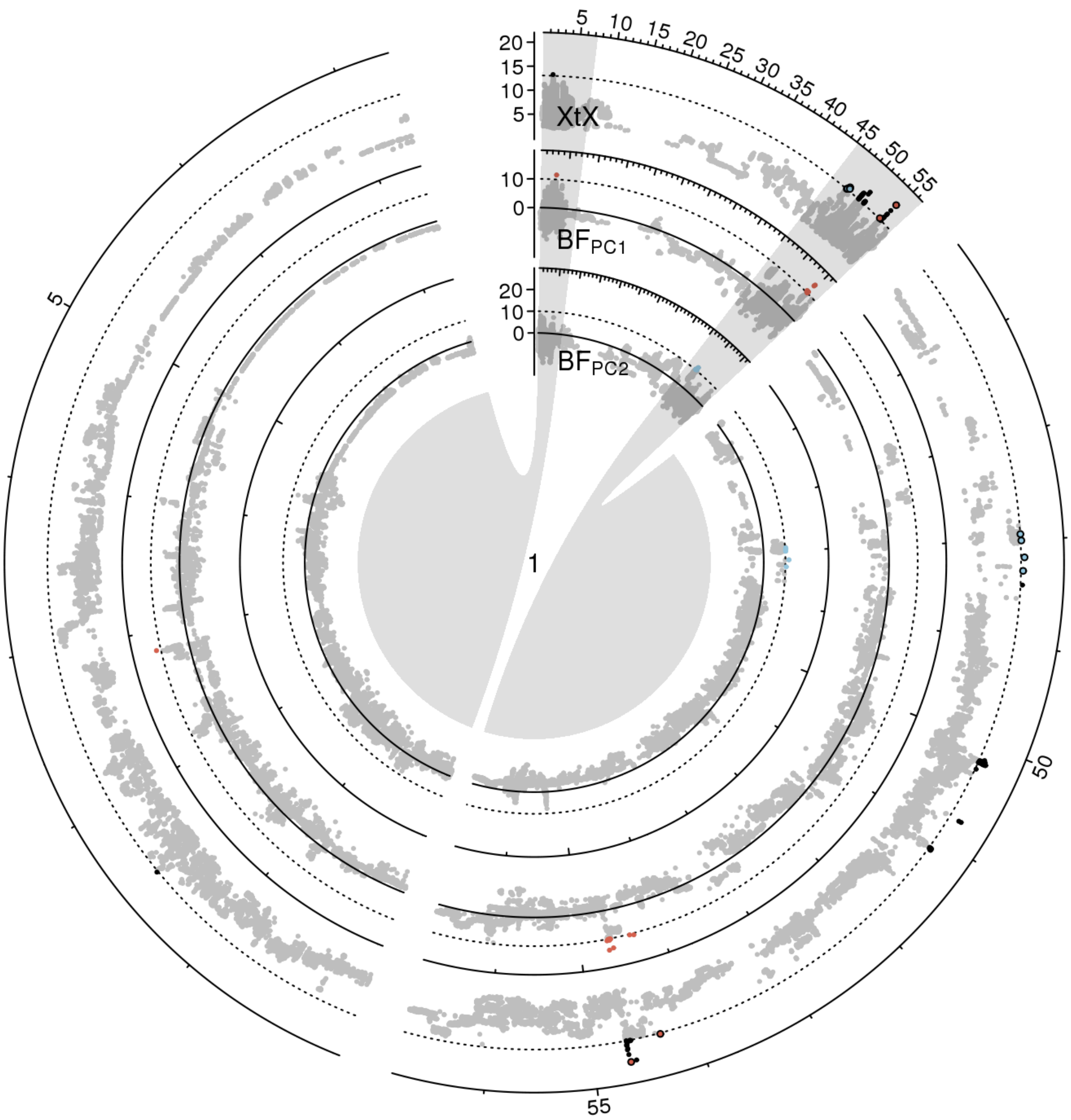


**Figure S 8:** Manhattan plot of the *XtX* statistic and of genotype-environment associations with the first two environmental principal components in Bayes factors for chromosome 1 in scenario A. Euchromatic regions on the chromosome arms are enlarged.

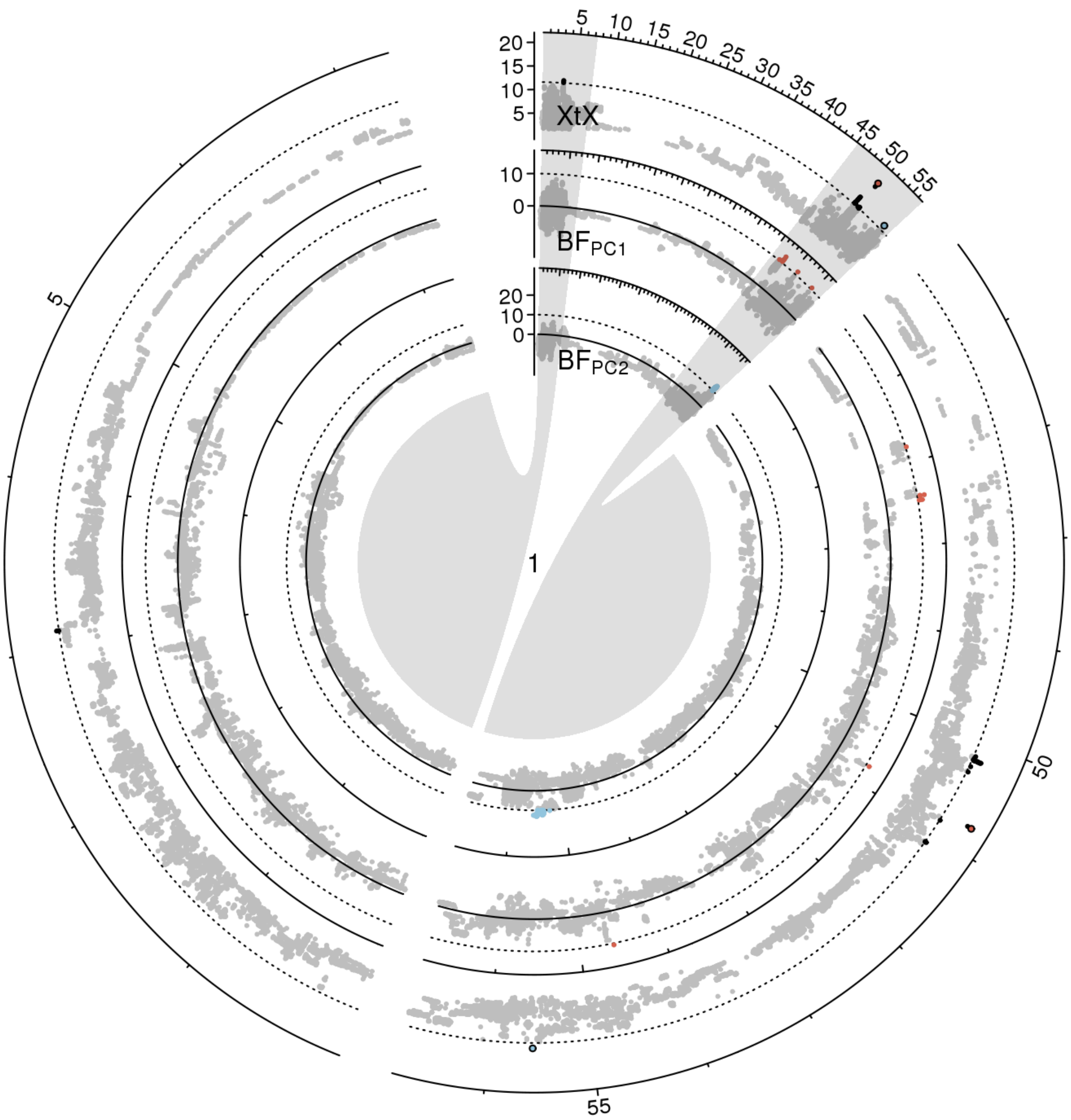


**Figure S 9:** Manhattan plot of the *XtX* statistic and of genotype-environment associations with the first two environmental principal components in Bayes factors for chromosome 1 in scenario B. Euchromatic regions on the chromosome arms are enlarged.

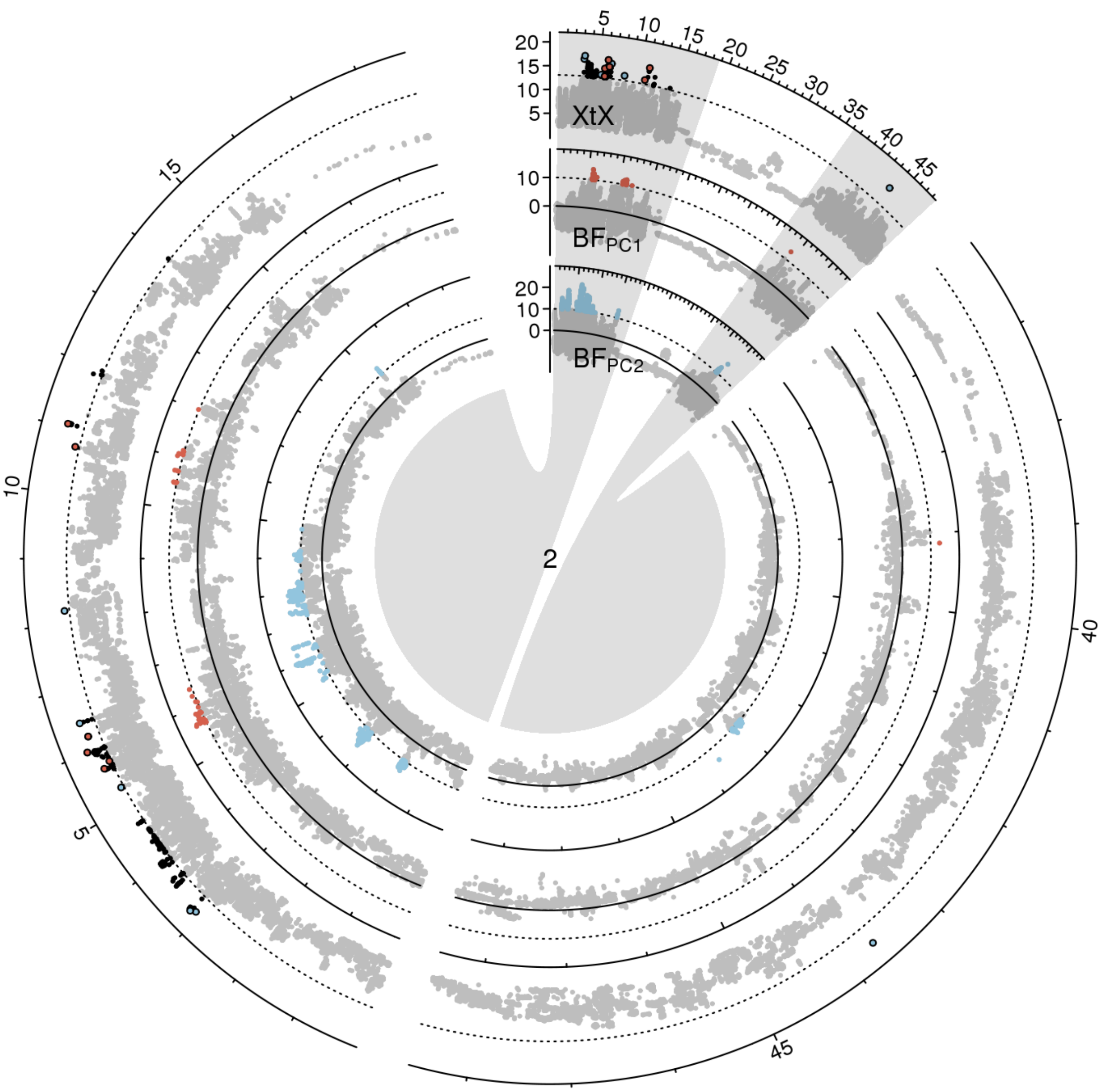


**Figure S 10:** Manhattan plot of the *XtX* statistic and of genotype-environment associations with the first two environmental principal components in Bayes factors for chromosome 2 in scenario A. Euchromatic regions on the chromosome arms are enlarged.

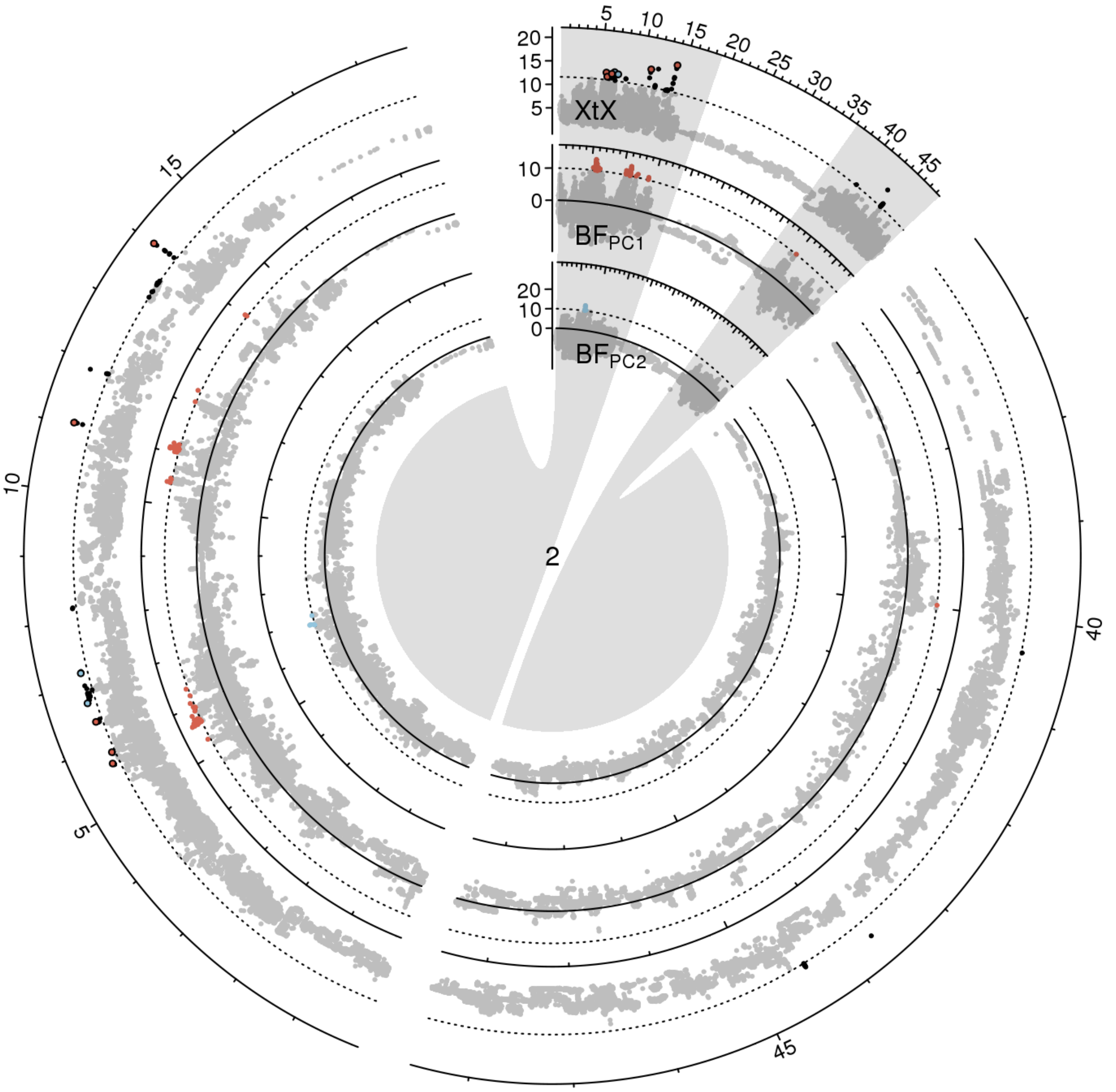


**Figure S 11:** Manhattan plot of the *XtX* statistic and of genotype-environment associations with the first two environmental principal components in Bayes factors for chromosome 2 in scenario B. Euchromatic regions on the chromosome arms are enlarged.

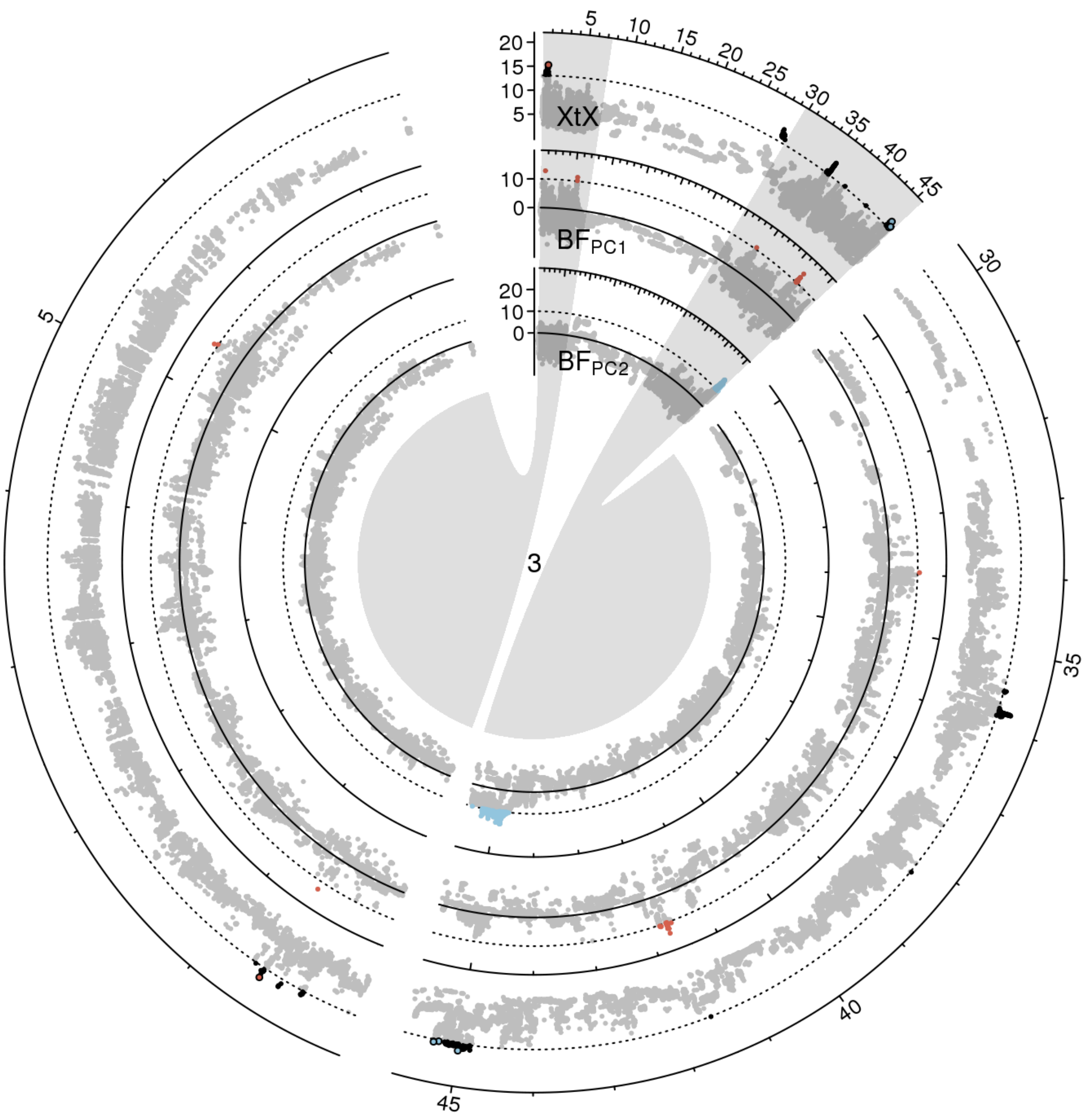


**Figure S 12:** Manhattan plot of the *XtX* statistic and of genotype-environment associations with the first two environmental principal components in Bayes factors for chromosome 3 in scenario A. Euchromatic regions on the chromosome arms are enlarged.

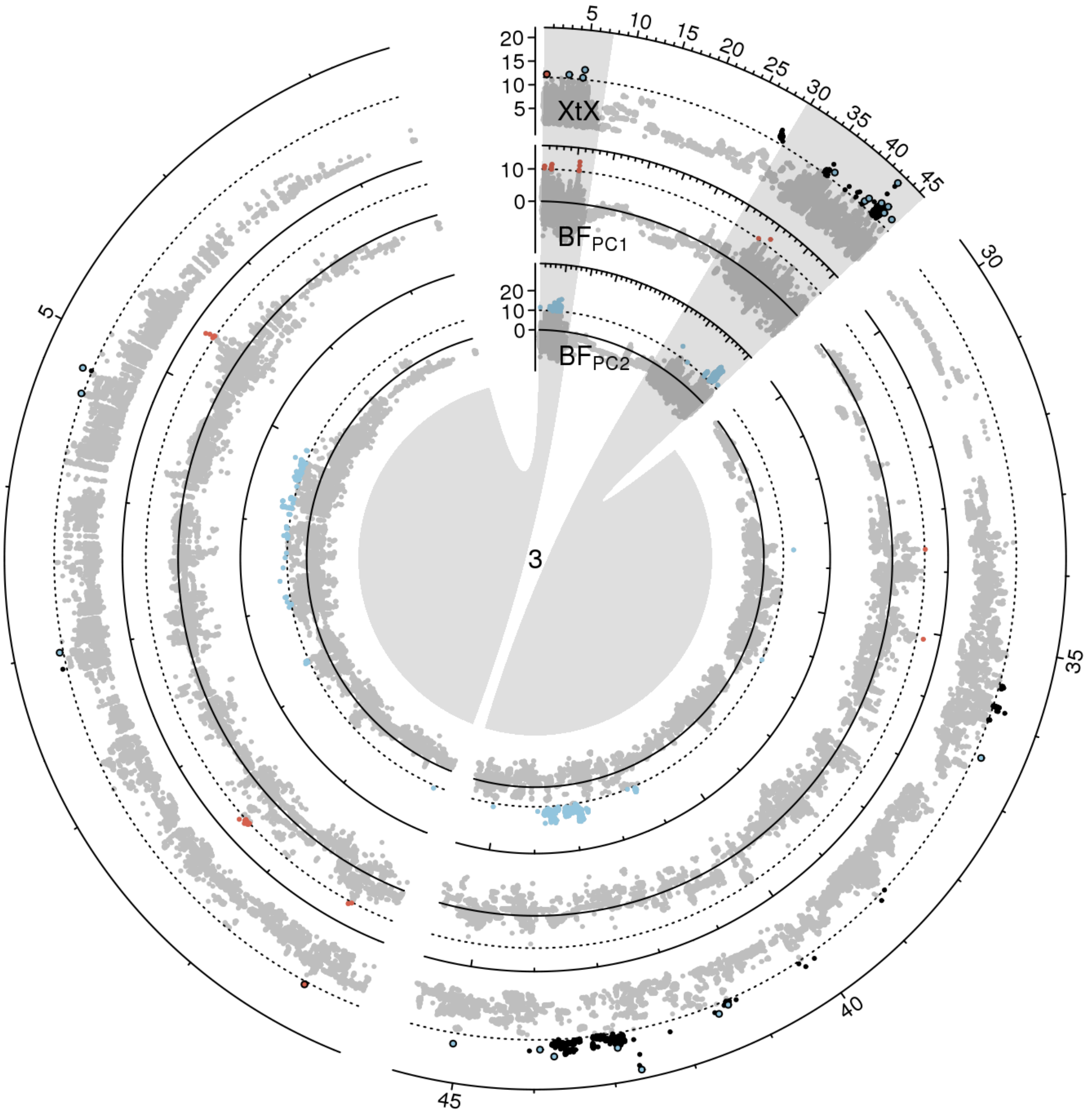


**Figure S 13:** Manhattan plot of the *XtX* statistic and of genotype-environment associations with the first two environmental principal components in Bayes factors for chromosome 3 in scenario B. Euchromatic regions on the chromosome arms are enlarged.

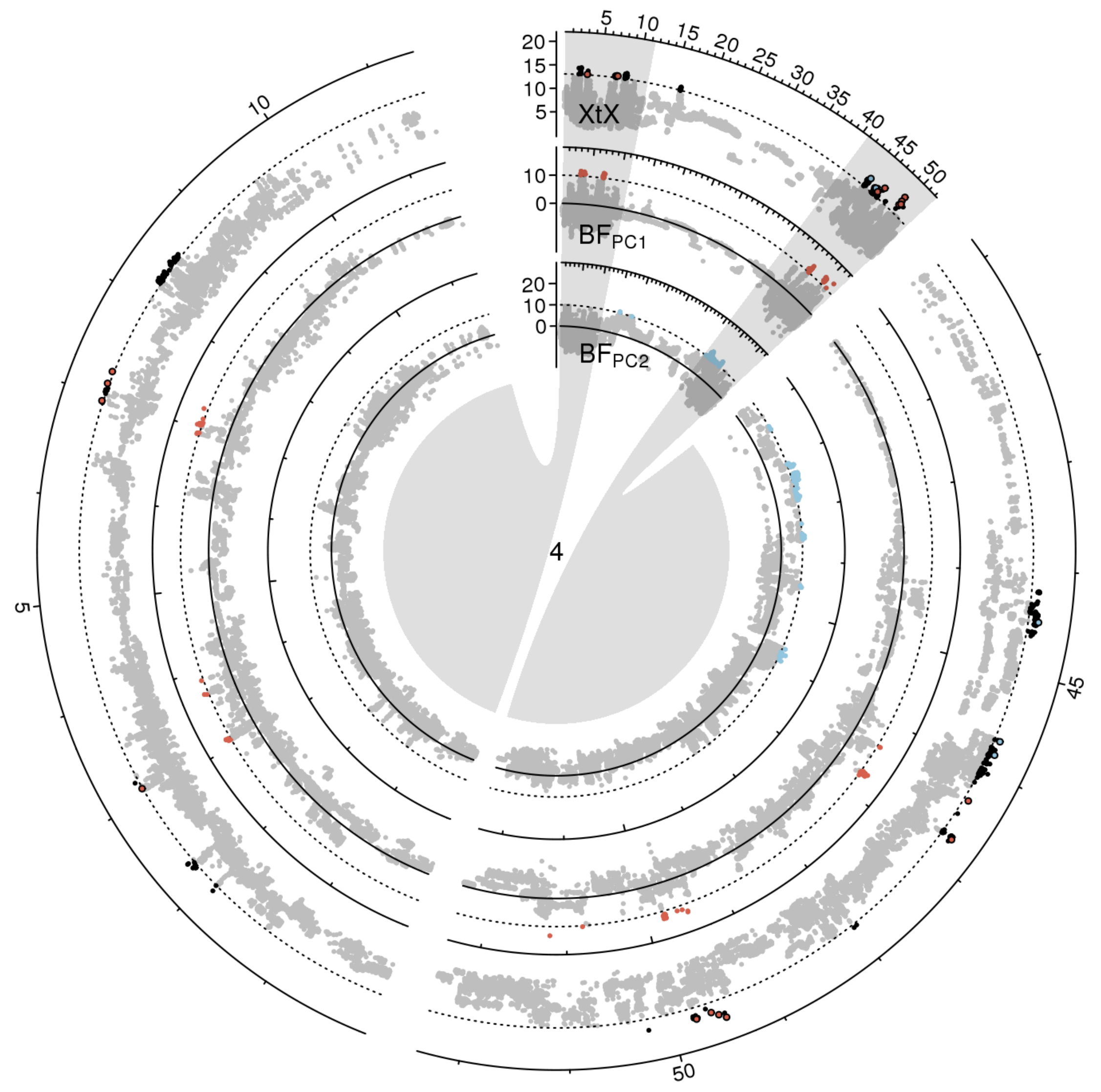


**Figure S 14:** Manhattan plot of the *XtX* statistic and of genotype-environment associations with the first two environmental principal components in Bayes factors for chromosome 4 in scenario A. Euchromatic regions on the chromosome arms are enlarged.

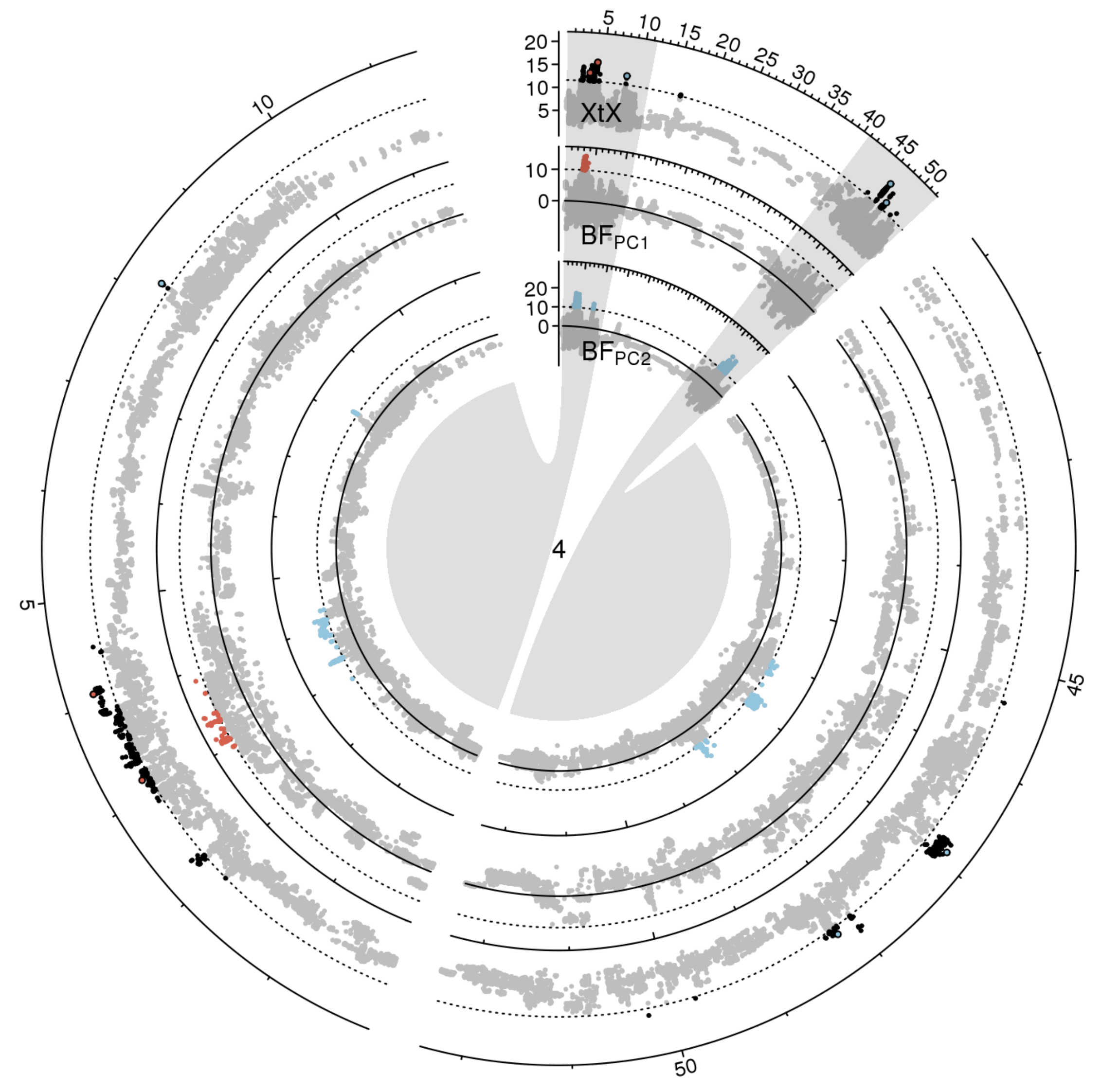


**Figure S 15:** Manhattan plot of the *XtX* statistic and of genotype-environment associations with the first two environmental principal components in Bayes factors for chromosome 4 in scenario B. Euchromatic regions on the chromosome arms are enlarged.

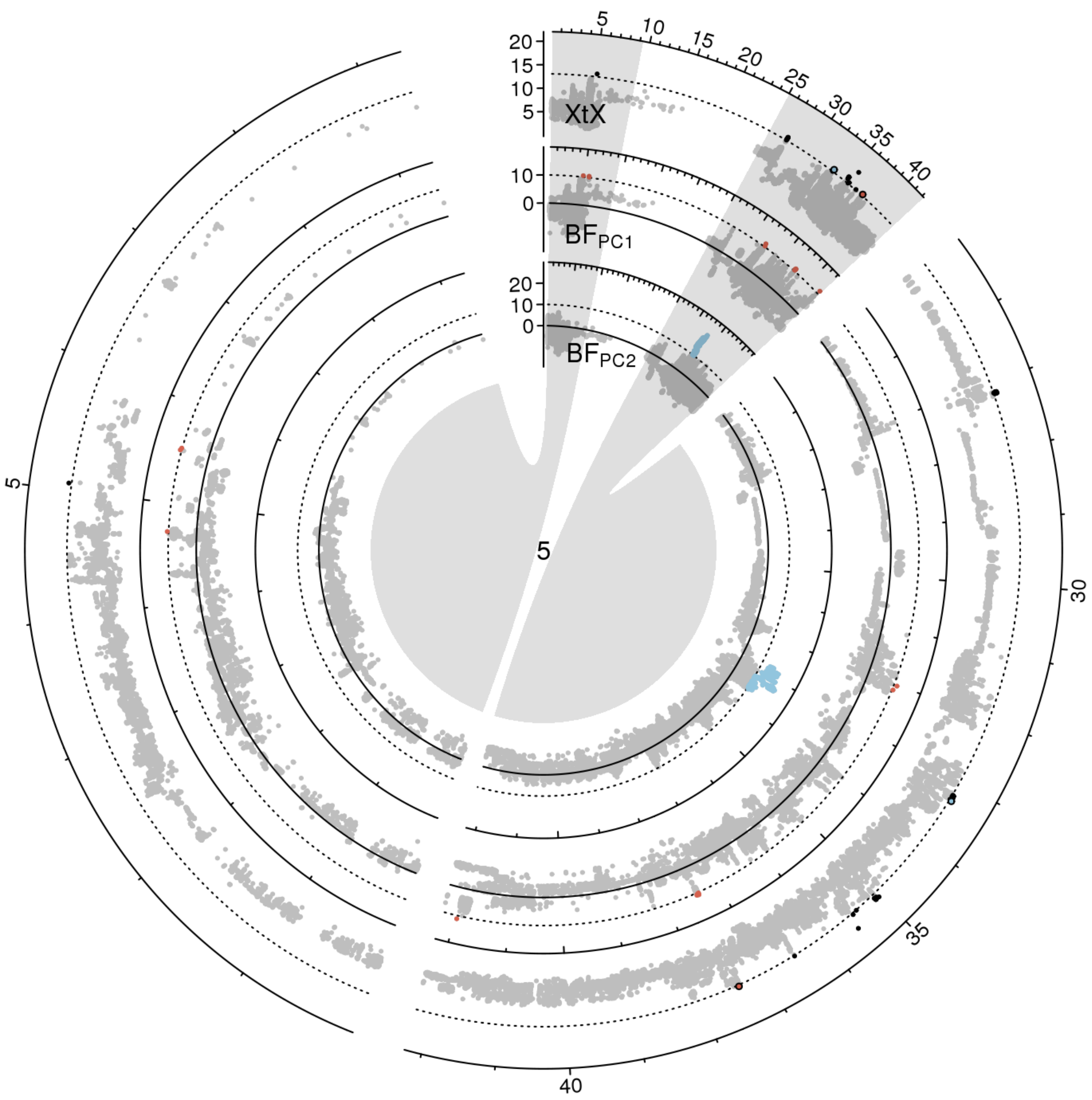


**Figure S 16:** Manhattan plot of the *XtX* statistic and of genotype-environment associations with the first two environmental principal components in Bayes factors for chromosome 5 in scenario A. Euchromatic regions on the chromosome arms are enlarged.

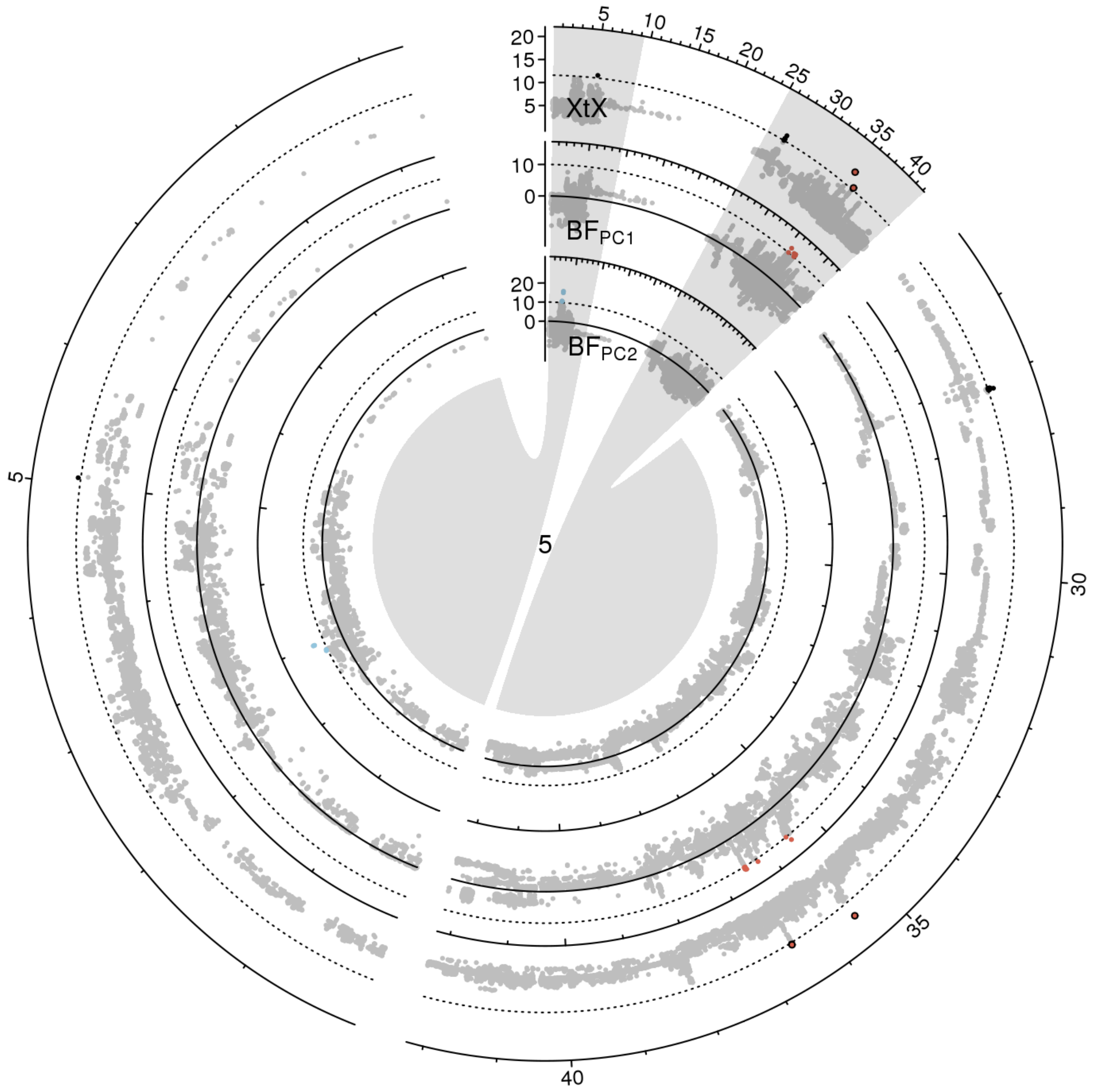


**Figure S 17:** Manhattan plot of the *XtX* statistic and of genotype-environment associations with the first two environmental principal components in Bayes factors for chromosome 5 in scenario B. Euchromatic regions on the chromosome arms are enlarged.

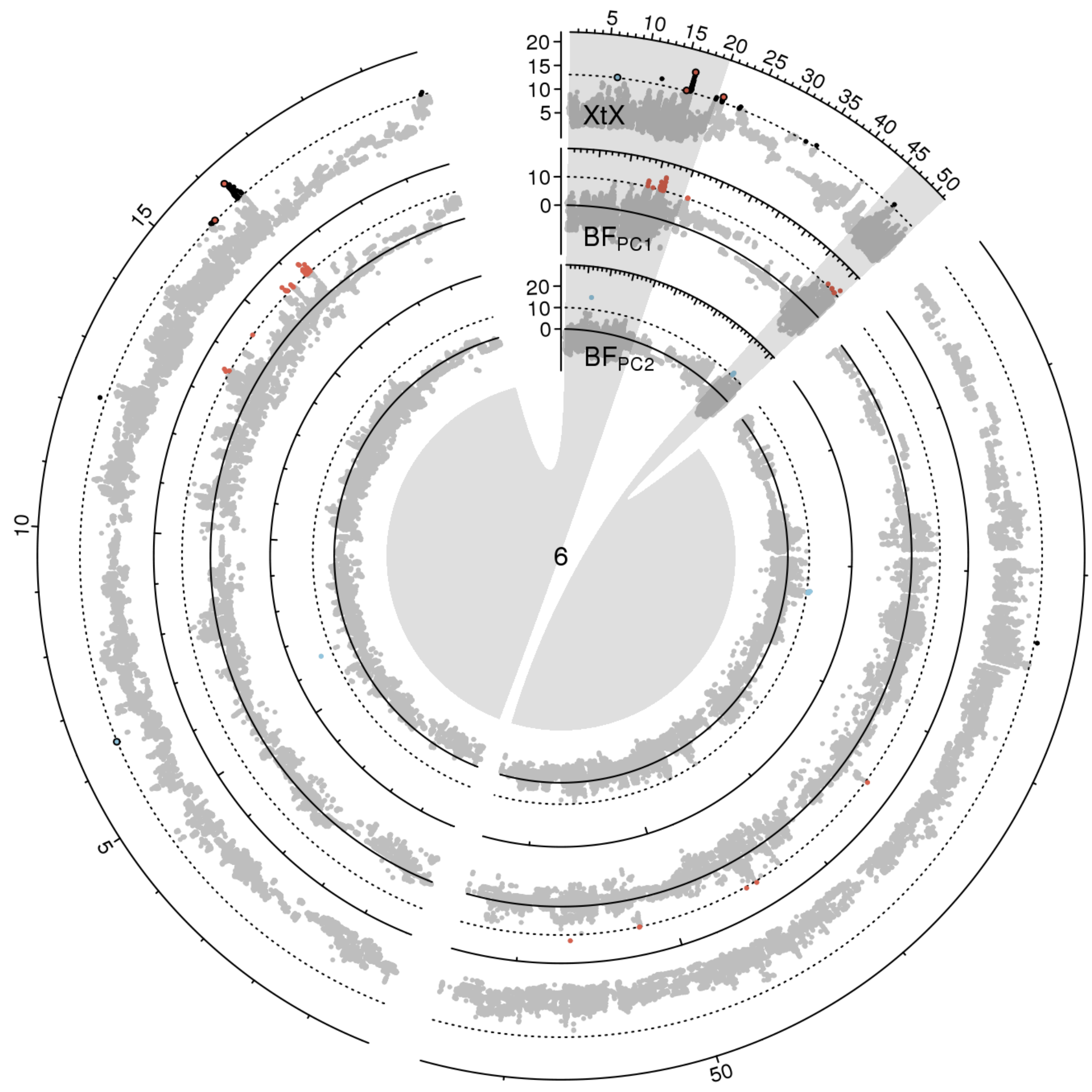


**Figure S 18:** Manhattan plot of the *XtX* statistic and of genotype-environment associations with the first two environmental principal components in Bayes factors for chromosome 6 in scenario A. Euchromatic regions on the chromosome arms are enlarged.

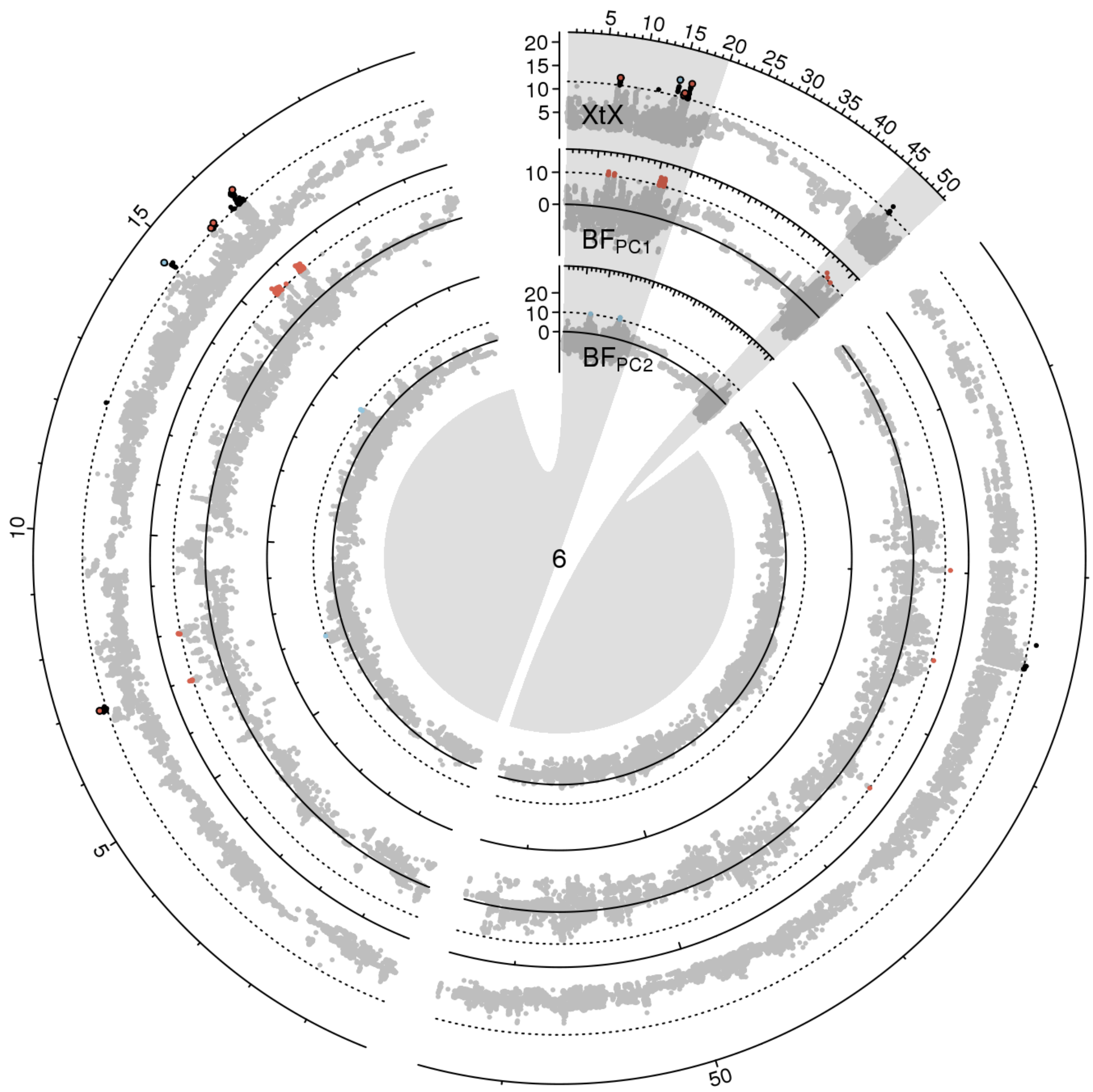


**Figure S 19:** Manhattan plot of the *XtX* statistic and of genotype-environment associations with the first two environmental principal components in Bayes factors for chromosome 6 in scenario B. Euchromatic regions on the chromosome arms are enlarged.

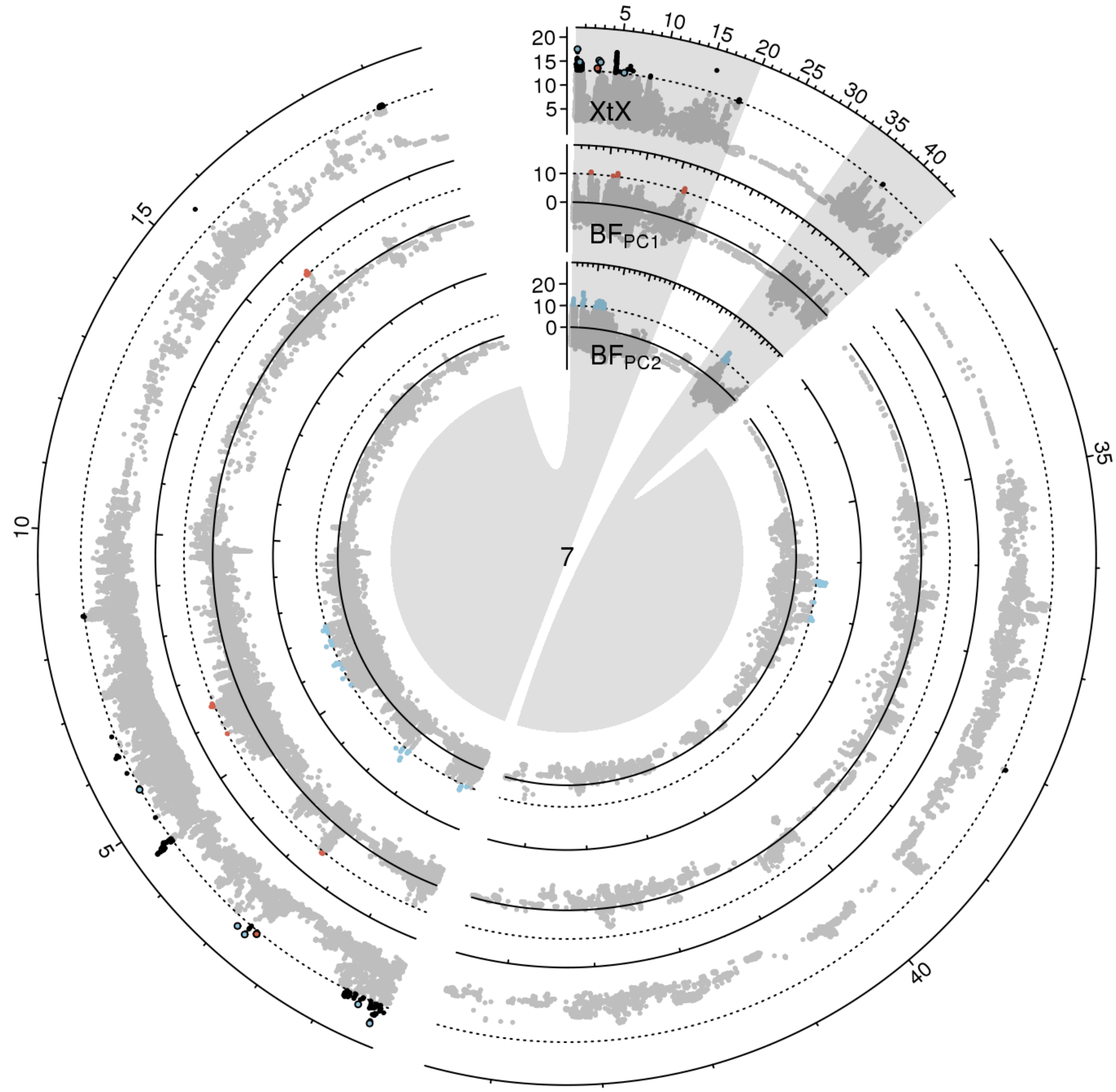


**Figure S 20:** Manhattan plot of the *XtX* statistic and of genotype-environment associations with the first two environmental principal components in Bayes factors for chromosome 7 in scenario A. Euchromatic regions on the chromosome arms are enlarged.

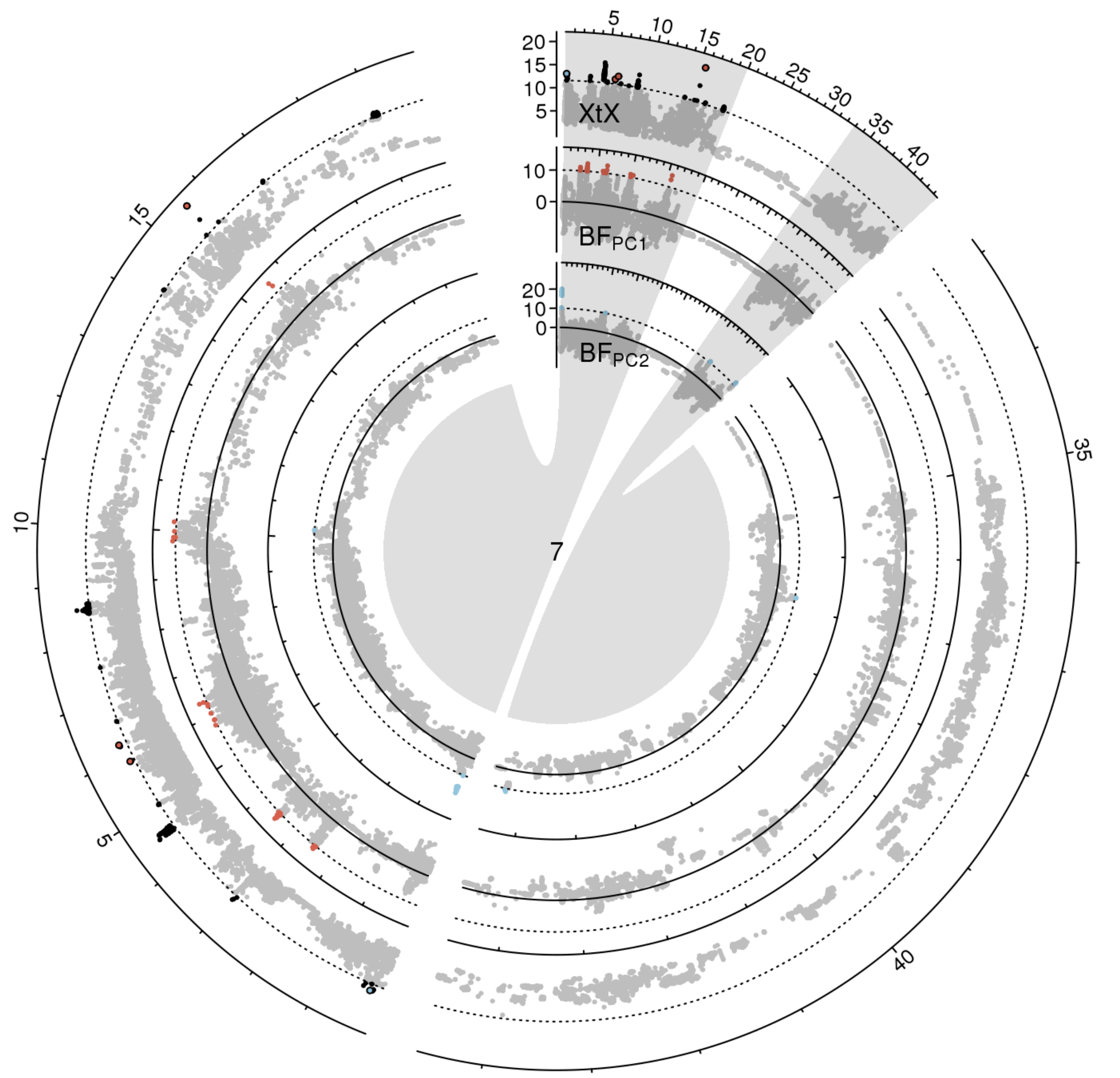


**Figure S 21:** Manhattan plot of the *XtX* statistic and of genotype-environment associations with the first two environmental principal components in Bayes factors for chromosome 7 in scenario B. Euchromatic regions on the chromosome arms are enlarged.

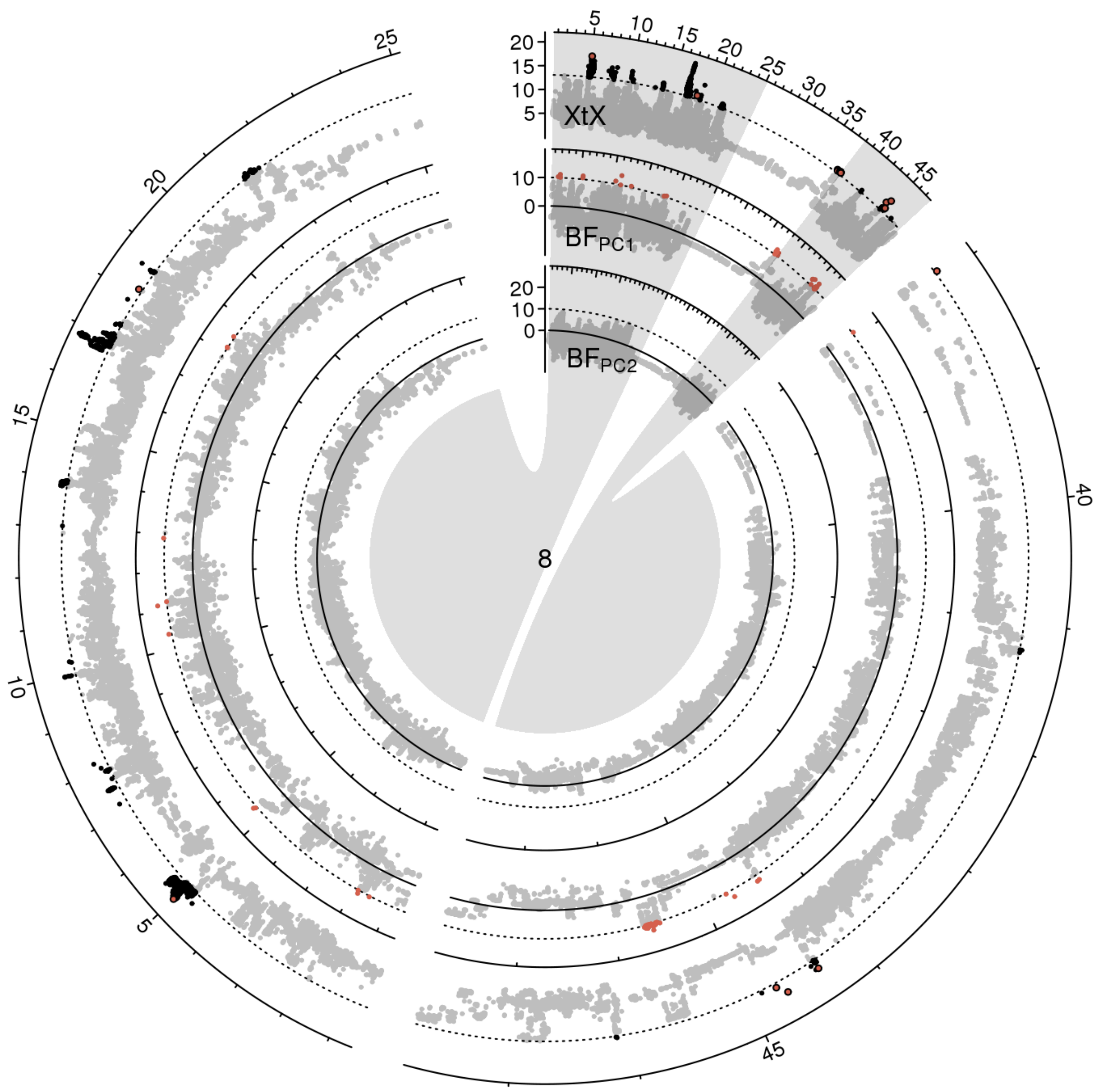


**Figure S 22:** Manhattan plot of the *XtX* statistic and of genotype-environment associations with the first two environmental principal components in Bayes factors for chromosome 8 in scenario A. Euchromatic regions on the chromosome arms are enlarged.

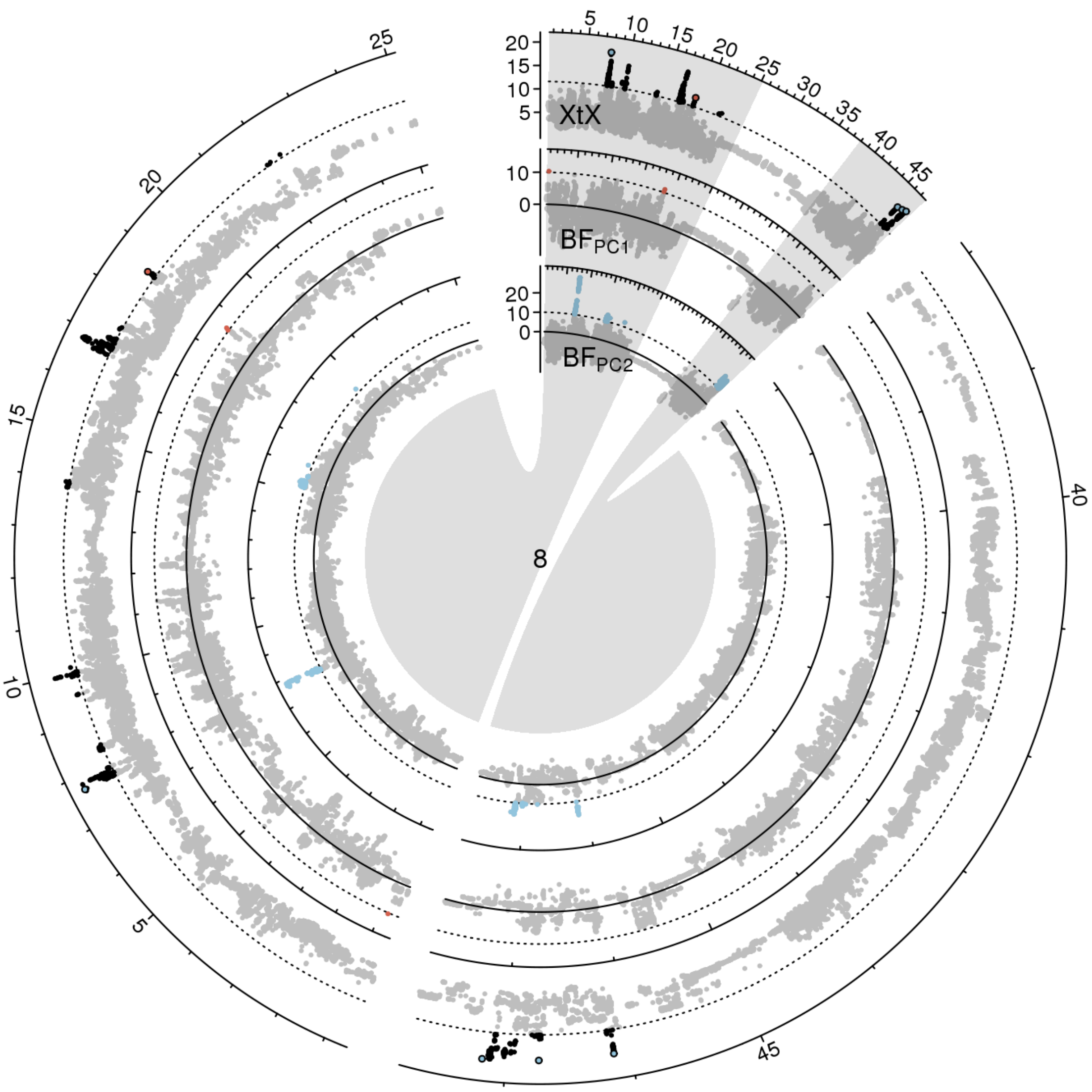


**Figure S 23:** Manhattan plot of the *XtX* statistic and of genotype-environment associations with the first two environmental principal components in Bayes factors for chromosome 8 in scenario B. Euchromatic regions on the chromosome arms are enlarged.

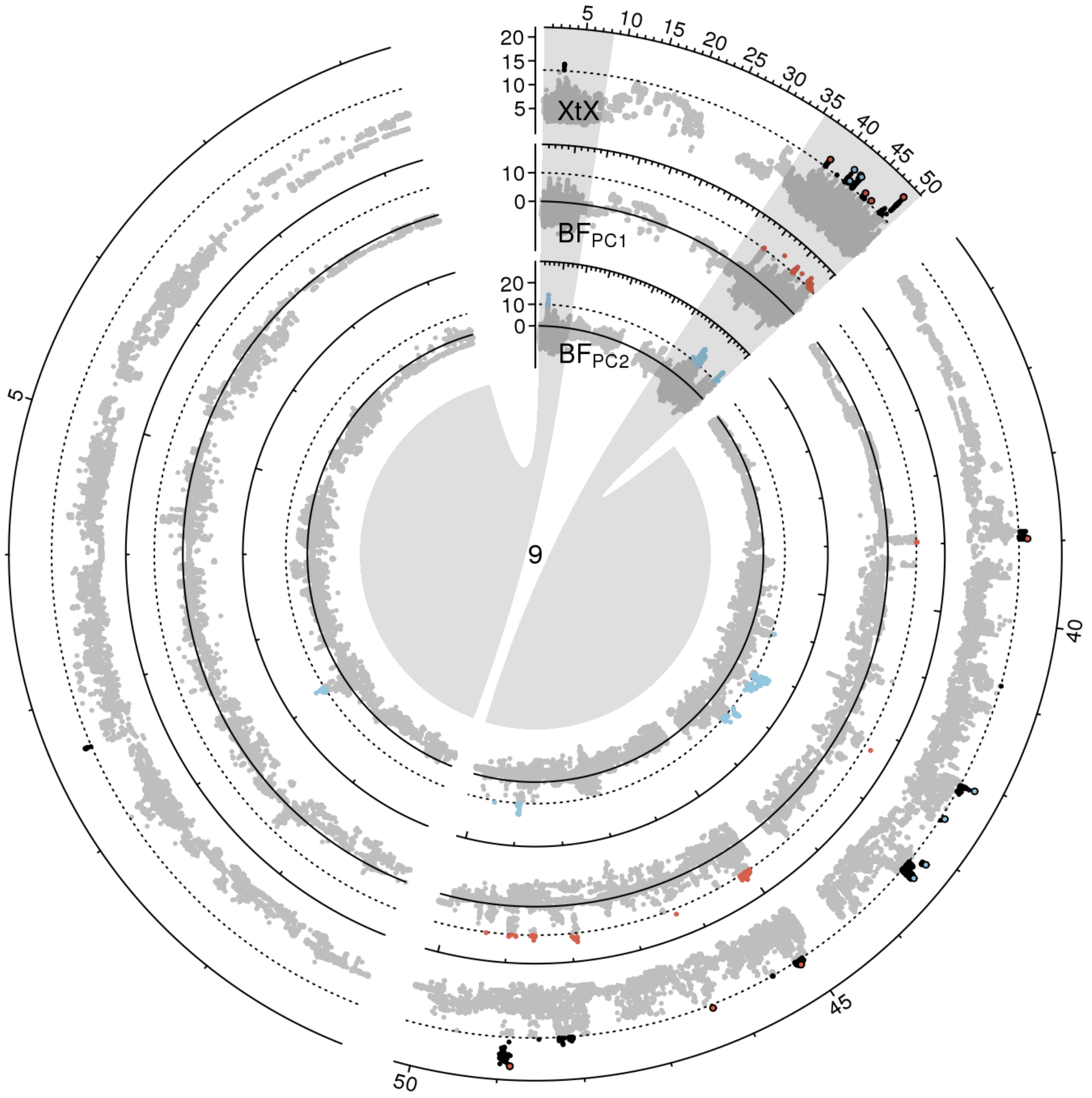


**Figure S 24:** Manhattan plot of the *XtX* statistic and of genotype-environment associations with the first two environmental principal components in Bayes factors for chromosome 9 in scenario A. Euchromatic regions on the chromosome arms are enlarged.

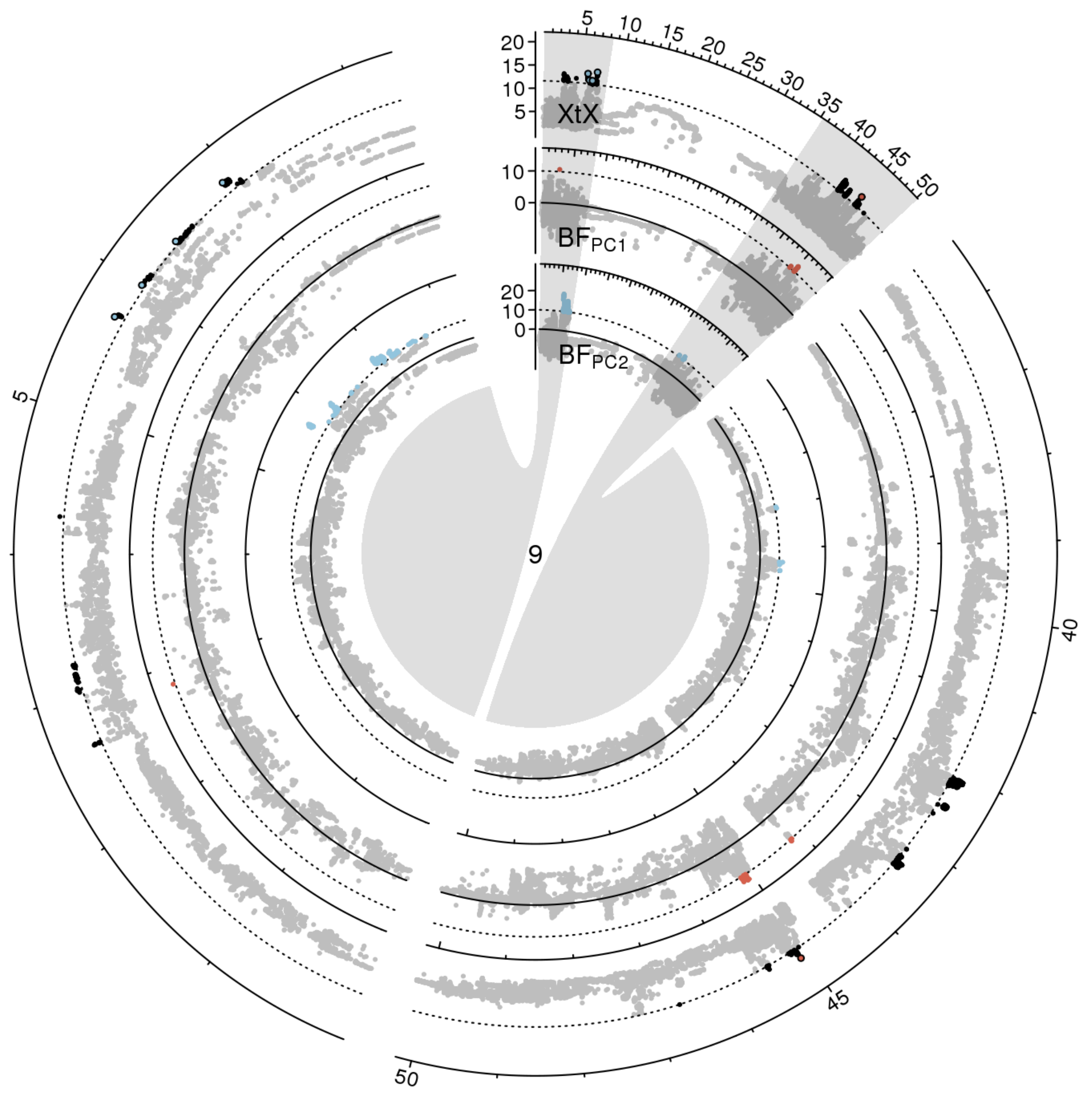


**Figure S 25:** Manhattan plot of the *XtX* statistic and of genotype-environment associations with the first two environmental principal components in Bayes factors for chromosome 9 in scenario B. Euchromatic regions on the chromosome arms are enlarged.

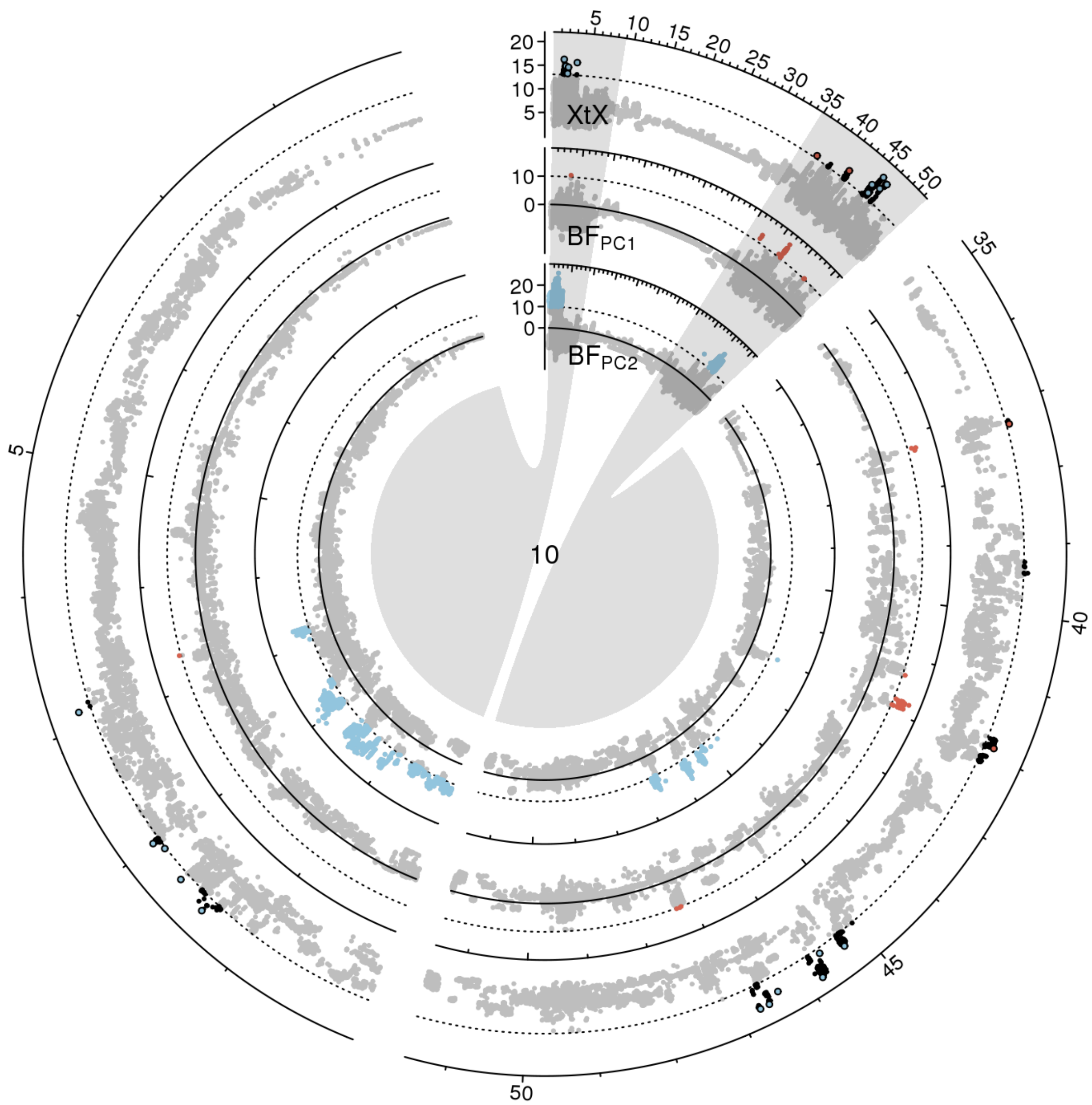


**Figure S 26:** Manhattan plot of the *XtX* statistic and of genotype-environment associations with the first two environmental principal components in Bayes factors for chromosome 10 in scenario A. Euchromatic regions on the chromosome arms are enlarged.

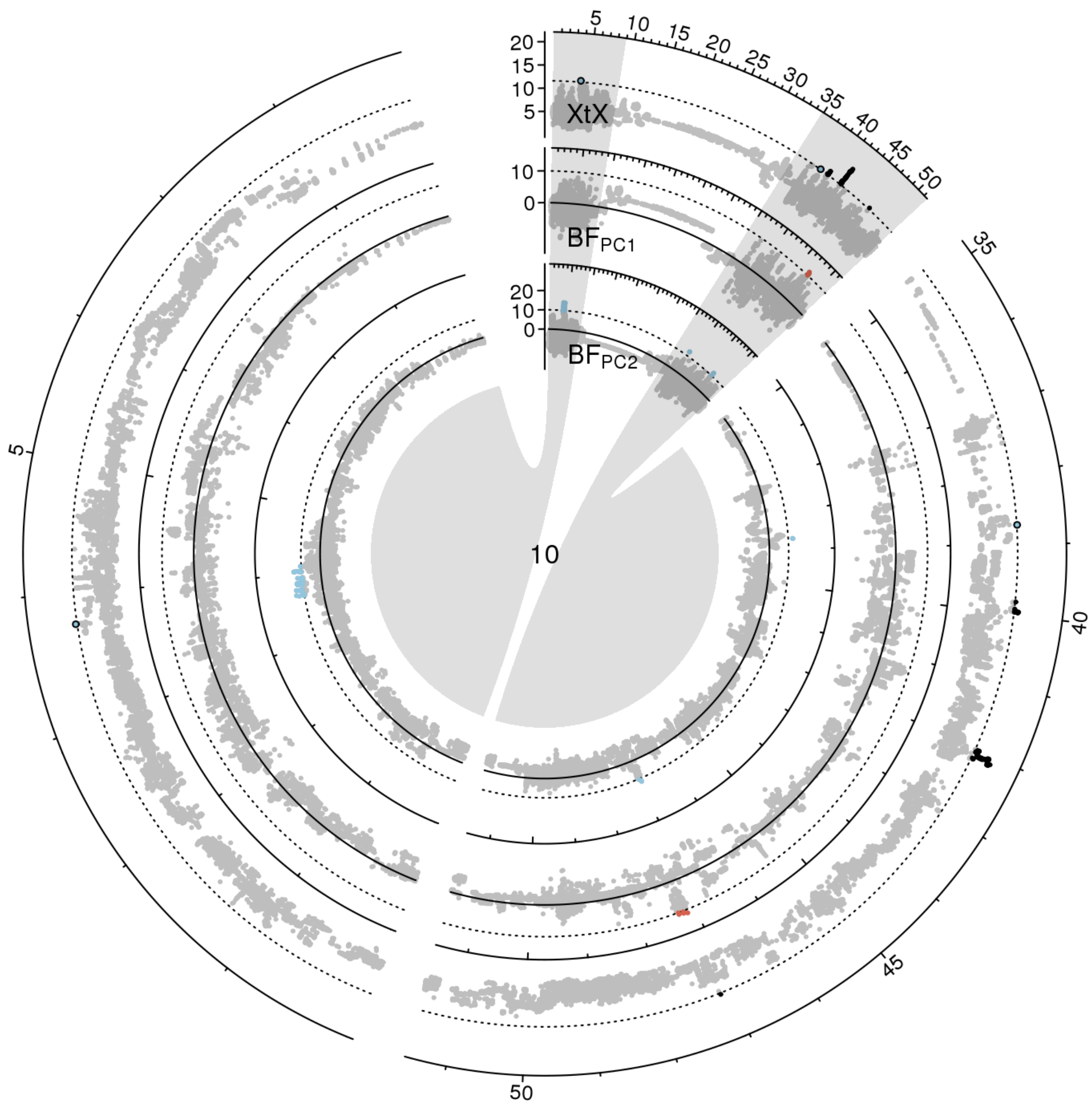


**Figure S 27:** Manhattan plot of the *XtX* statistic and of genotype-environment associations with the first two environmental principal components in Bayes factors for chromosome 10 in scenario B. Euchromatic regions on the chromosome arms are enlarged.

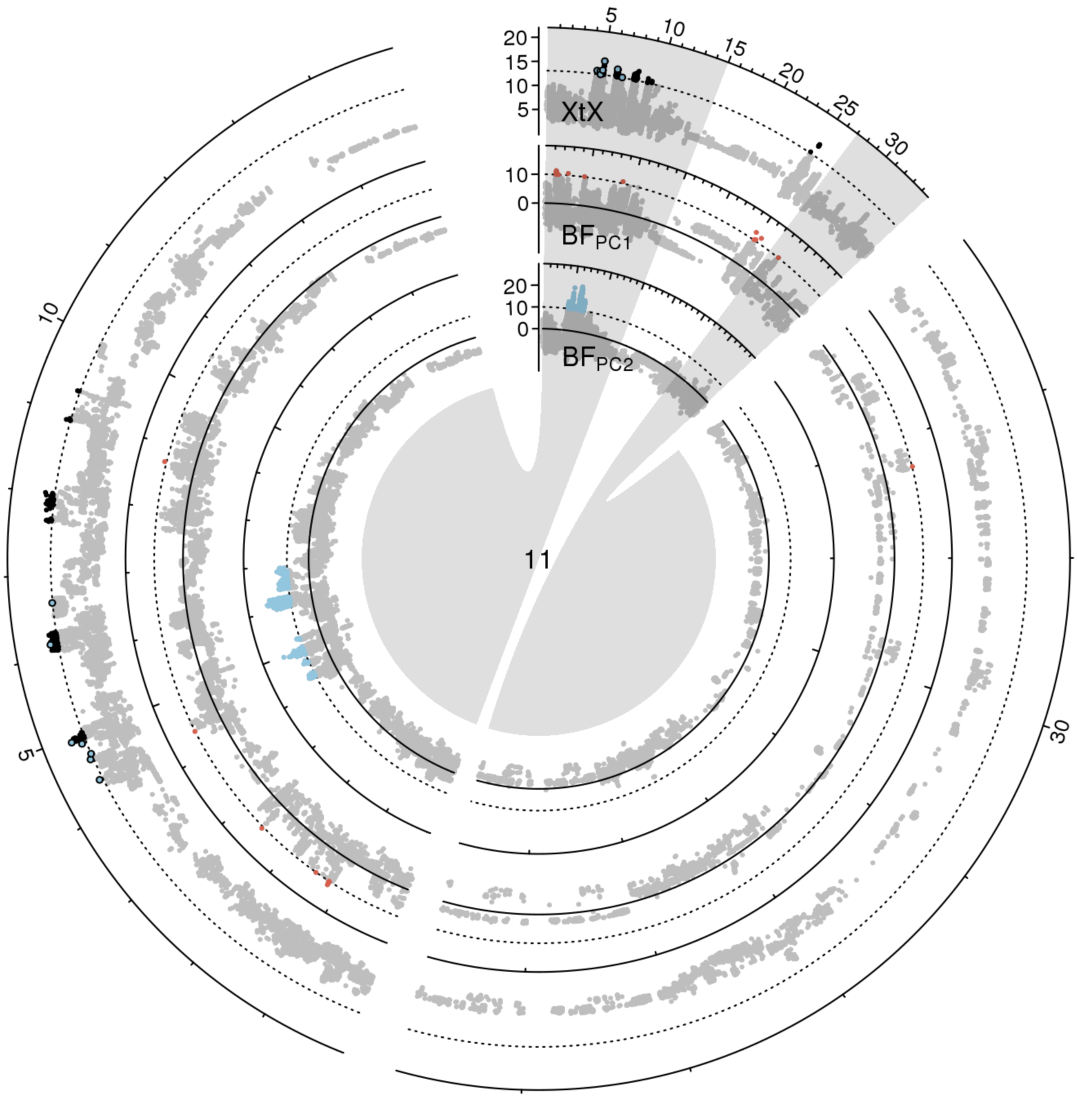


**Figure S 28:** Manhattan plot of the *XtX* statistic and of genotype-environment associations with the first two environmental principal components in Bayes factors for chromosome 11 in scenario A. Euchromatic regions on the chromosome arms are enlarged.

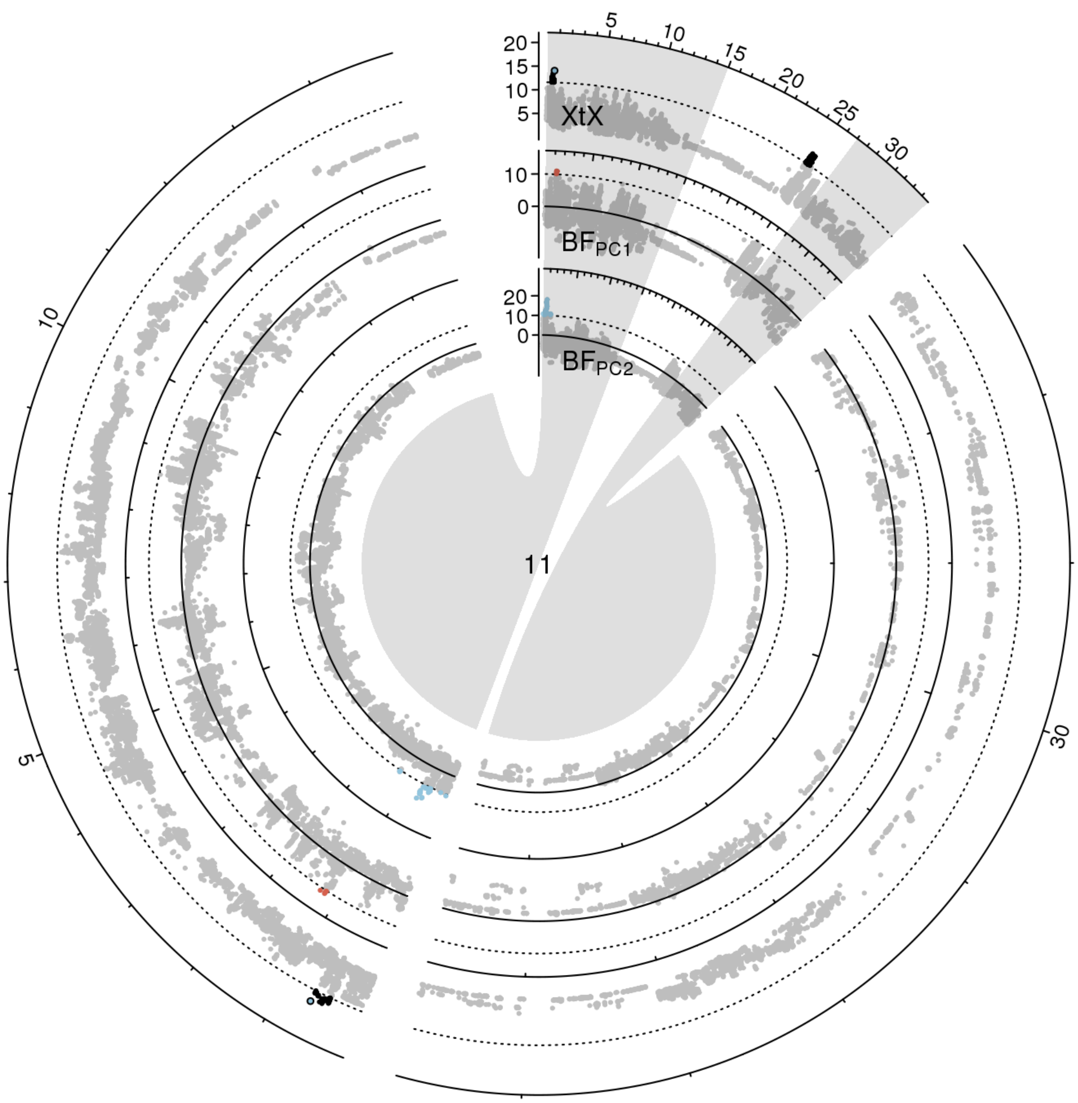


**Figure S 29:** Manhattan plot of the *XtX* statistic and of genotype-environment associations with the first two environmental principal components in Bayes factors for chromosome 11 in scenario B. Euchromatic regions on the chromosome arms are enlarged.

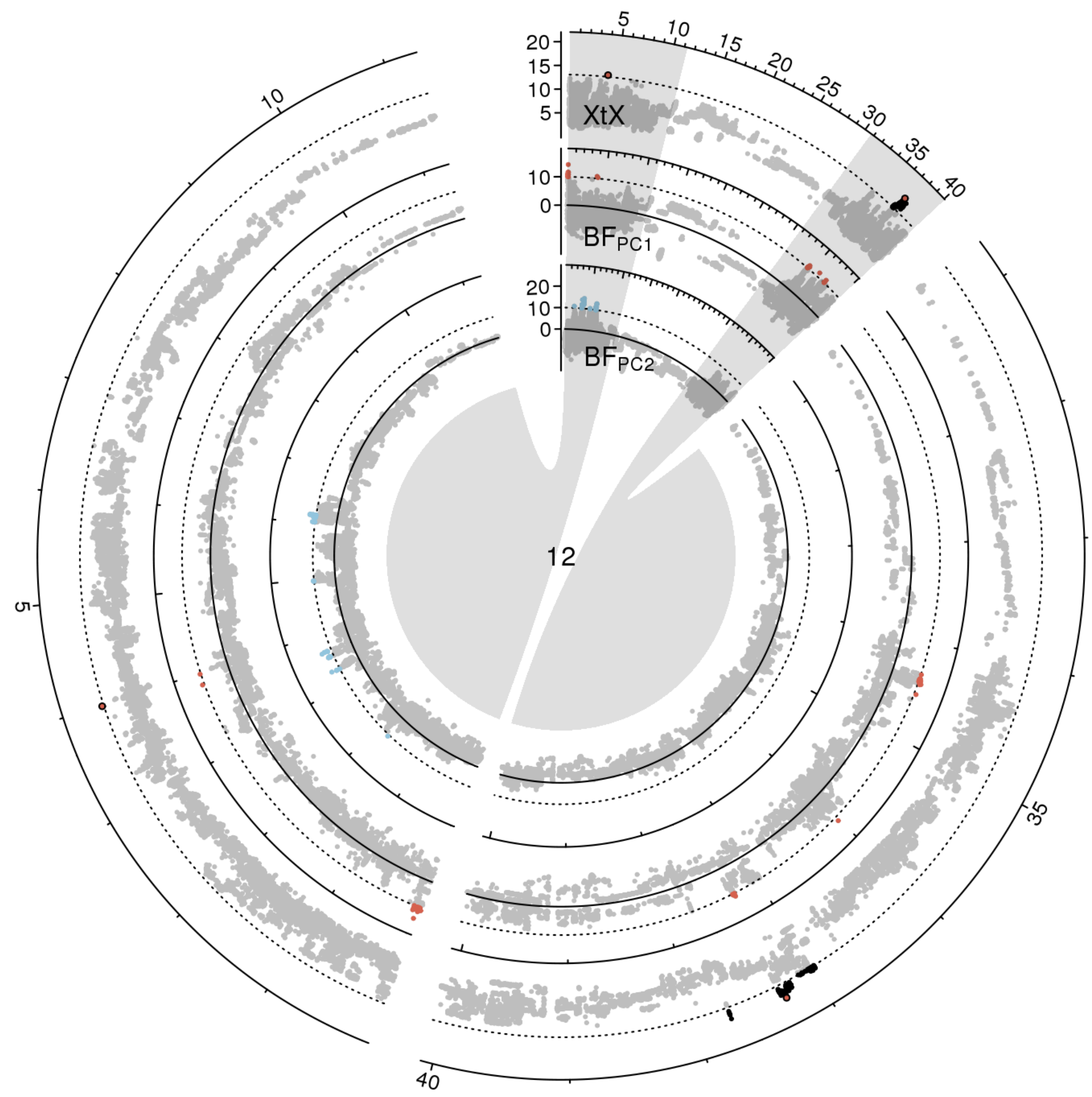


**Figure S 30:** Manhattan plot of the *XtX* statistic and of genotype-environment associations with the first two environmental principal components in Bayes factors for chromosome 12 in scenario A. Euchromatic regions on the chromosome arms are enlarged.

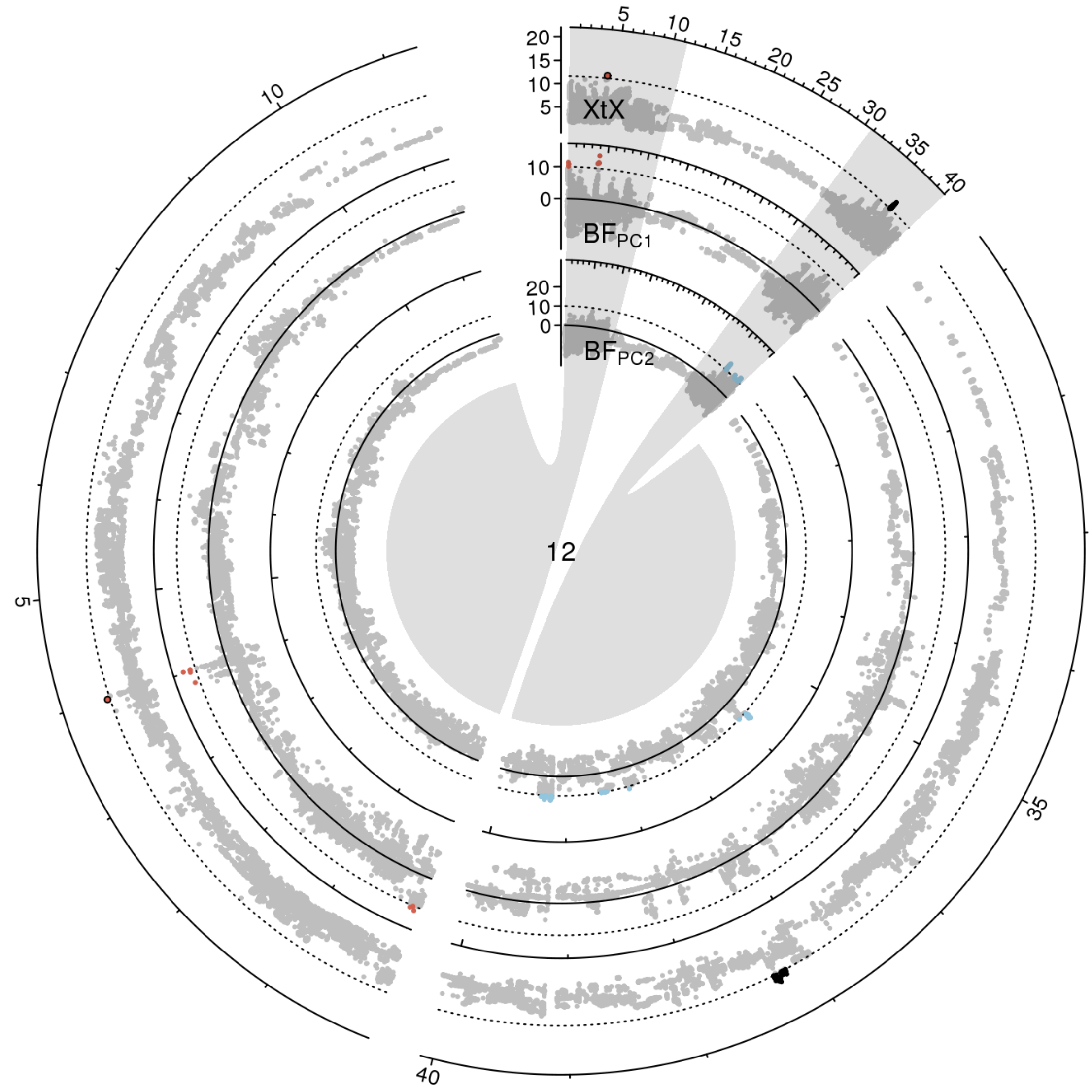


**Figure S 31:** Manhattan plot of the *XtX* statistic and of genotype-environment associations with the first two environmental principal components in Bayes factors for chromosome 12 in scenario B. Euchromatic regions on the chromosome arms are enlarged.

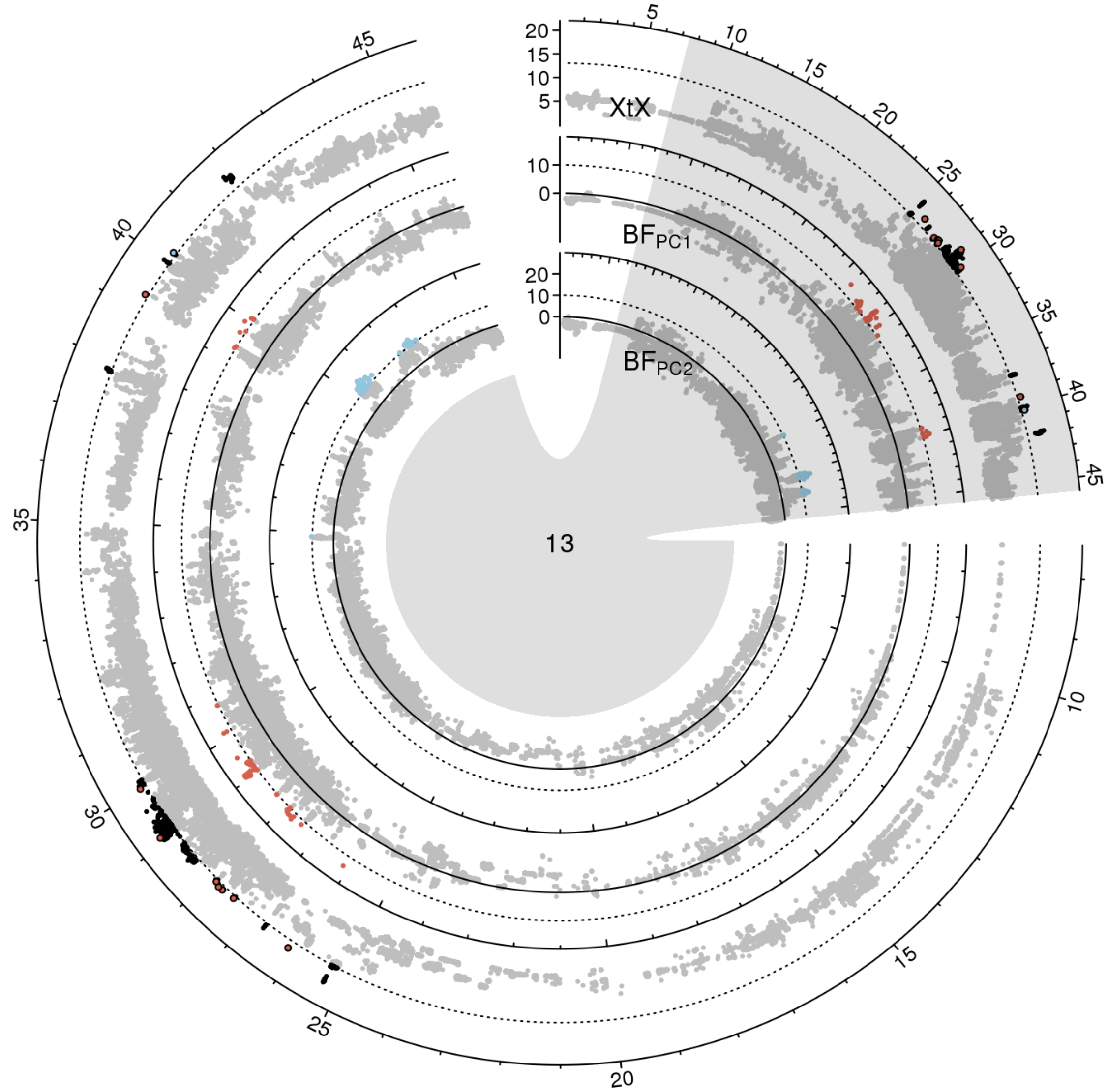


**Figure S 32:** Manhattan plot of the *XtX* statistic and of genotype-environment associations with the first two environmental principal components in Bayes factors for chromosome 13 in scenario A. Euchromatic regions on the chromosome arms are enlarged.

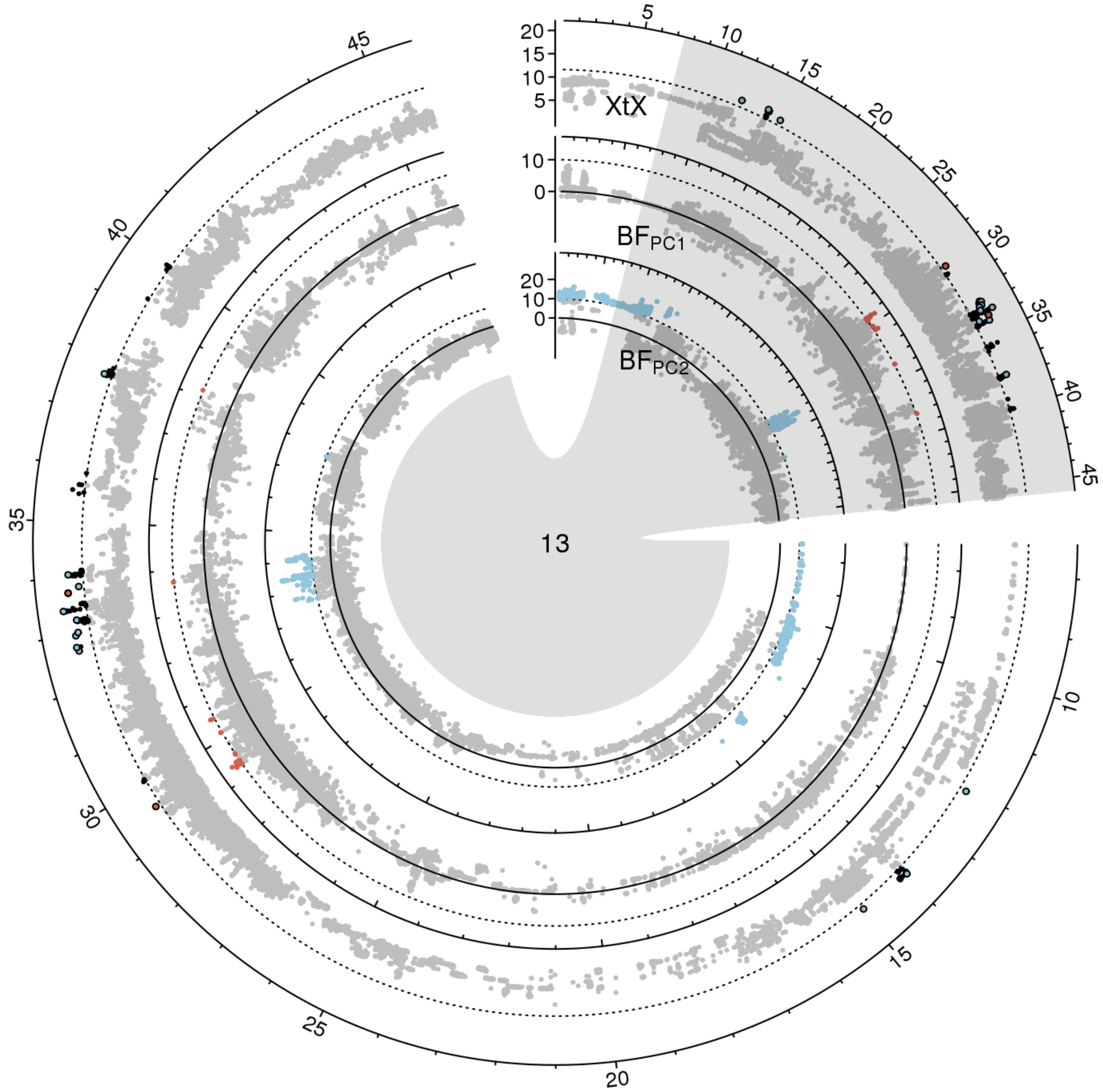


**Figure S 33:** Manhattan plot of the *XtX* statistic and of genotype-environment associations with the first two environmental principal components in Bayes factors for chromosome 13 in scenario B. Euchromatic regions on the chromosome arms are enlarged.

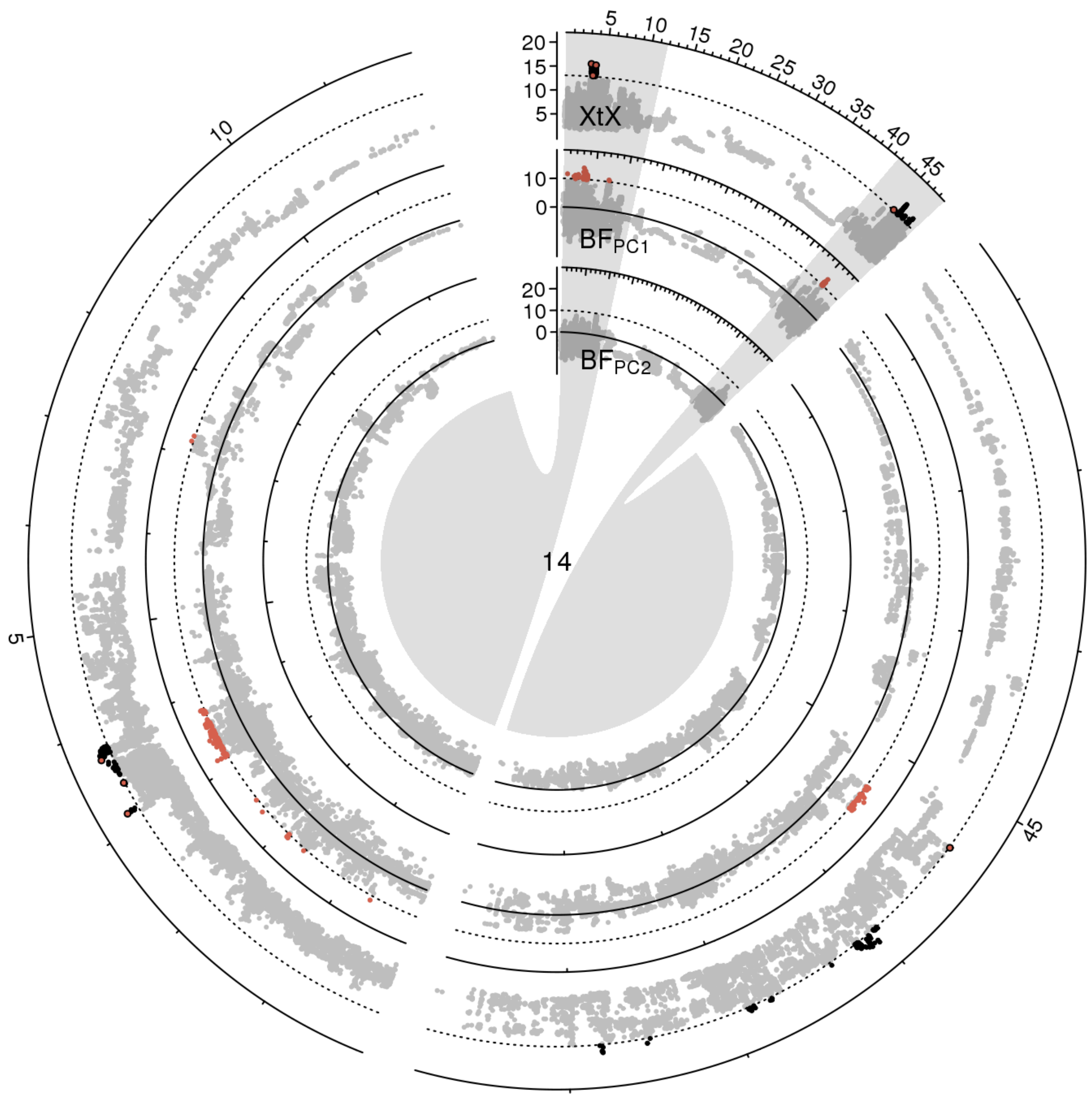


**Figure S 34:** Manhattan plot of the *XtX* statistic and of genotype-environment associations with the first two environmental principal components in Bayes factors for chromosome 14 in scenario A. Euchromatic regions on the chromosome arms are enlarged.

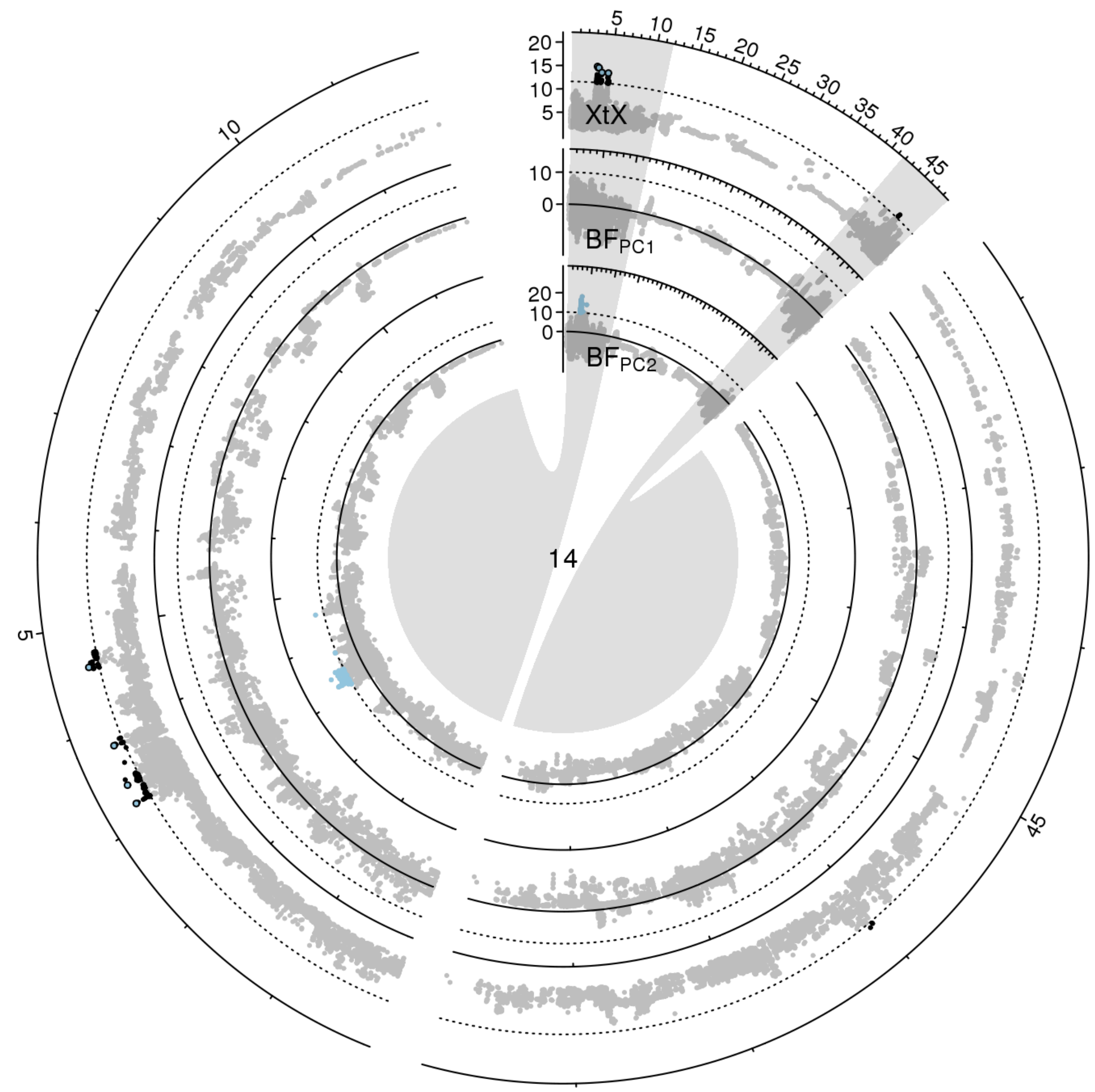


**Figure S 35:** Manhattan plot of the *XtX* statistic and of genotype-environment associations with the first two environmental principal components in Bayes factors for chromosome 14 in scenario B. Euchromatic regions on the chromosome arms are enlarged.

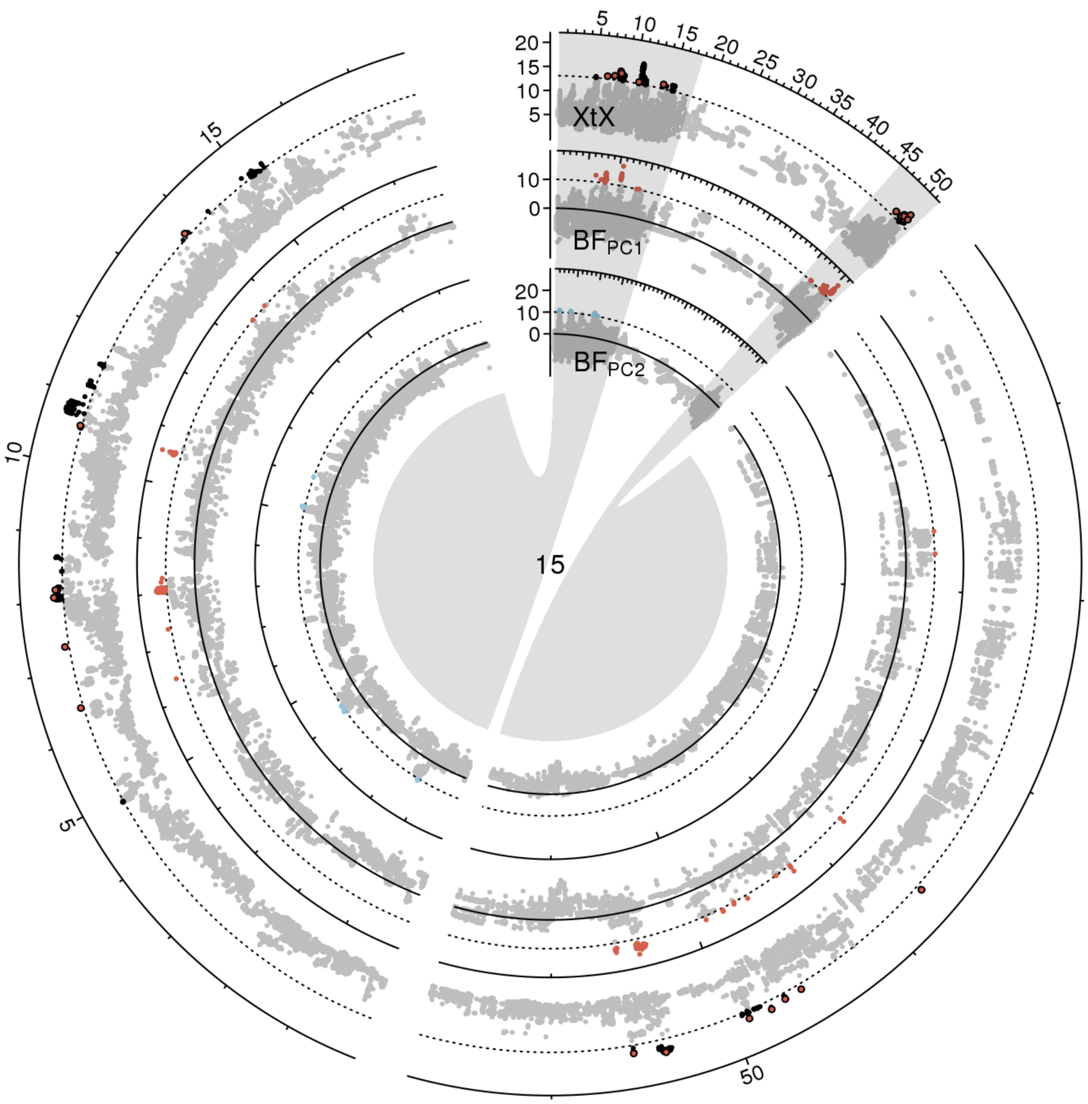


**Figure S 36:** Manhattan plot of the *XtX* statistic and of genotype-environment associations with the first two environmental principal components in Bayes factors for chromosome 15 in scenario A. Euchromatic regions on the chromosome arms are enlarged.

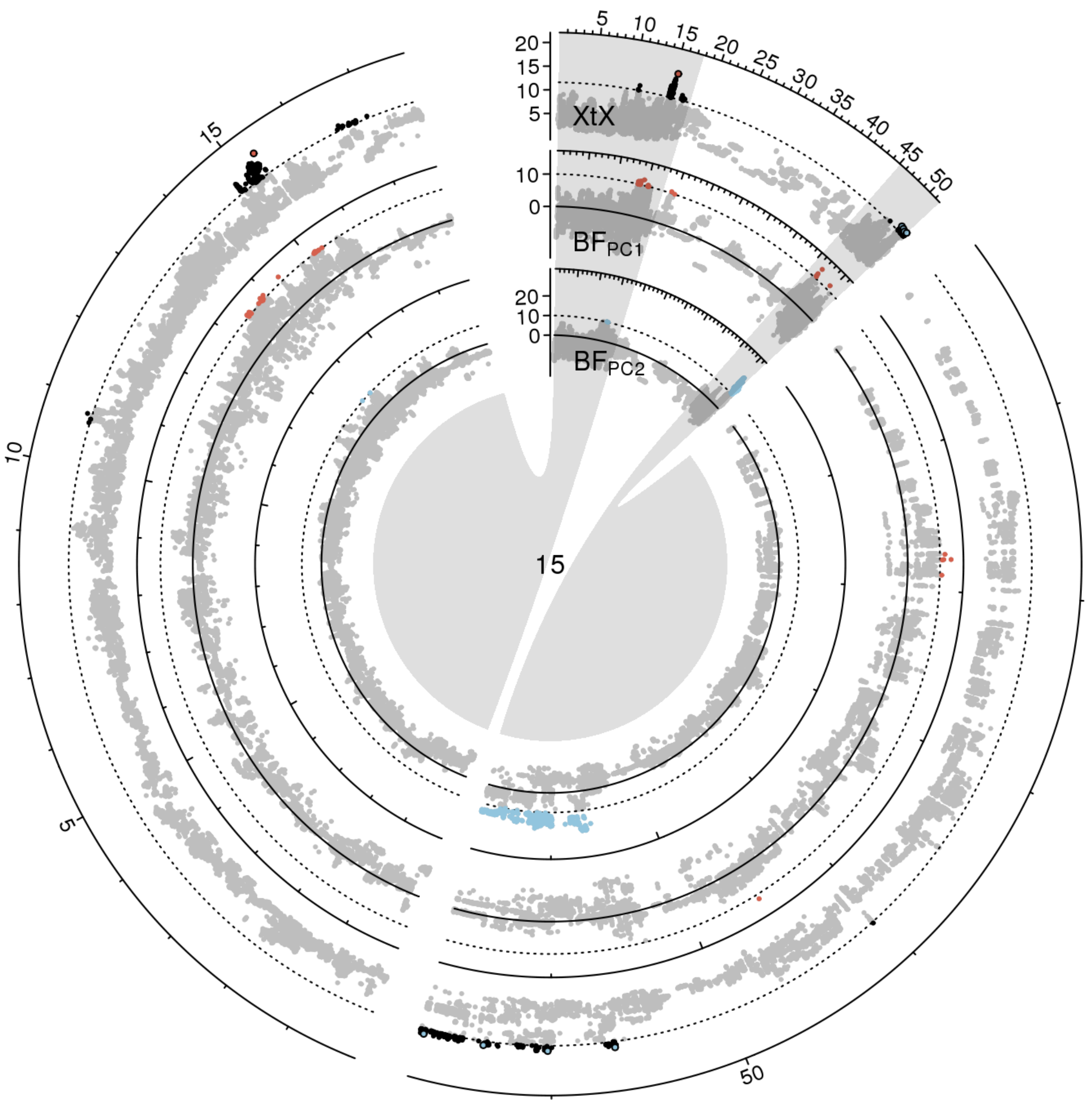


**Figure S 37:** Manhattan plot of the *XtX* statistic and of genotype-environment associations with the first two environmental principal components in Bayes factors for chromosome 15 in scenario B. Euchromatic regions on the chromosome arms are enlarged.

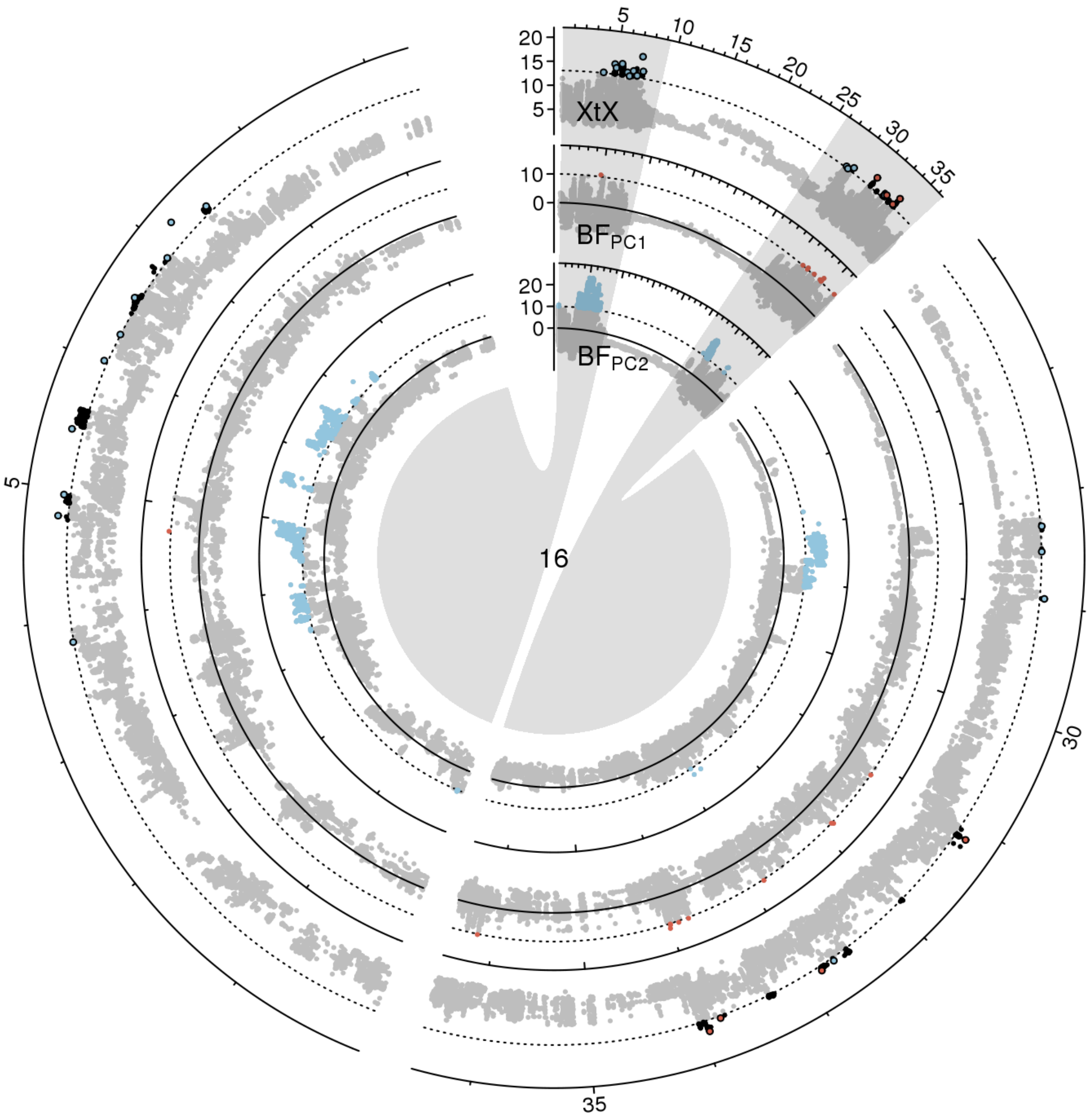


**Figure S 38:** Manhattan plot of the *XtX* statistic and of genotype-environment associations with the first two environmental principal components in Bayes factors for chromosome 16 in scenario A. Euchromatic regions on the chromosome arms are enlarged.

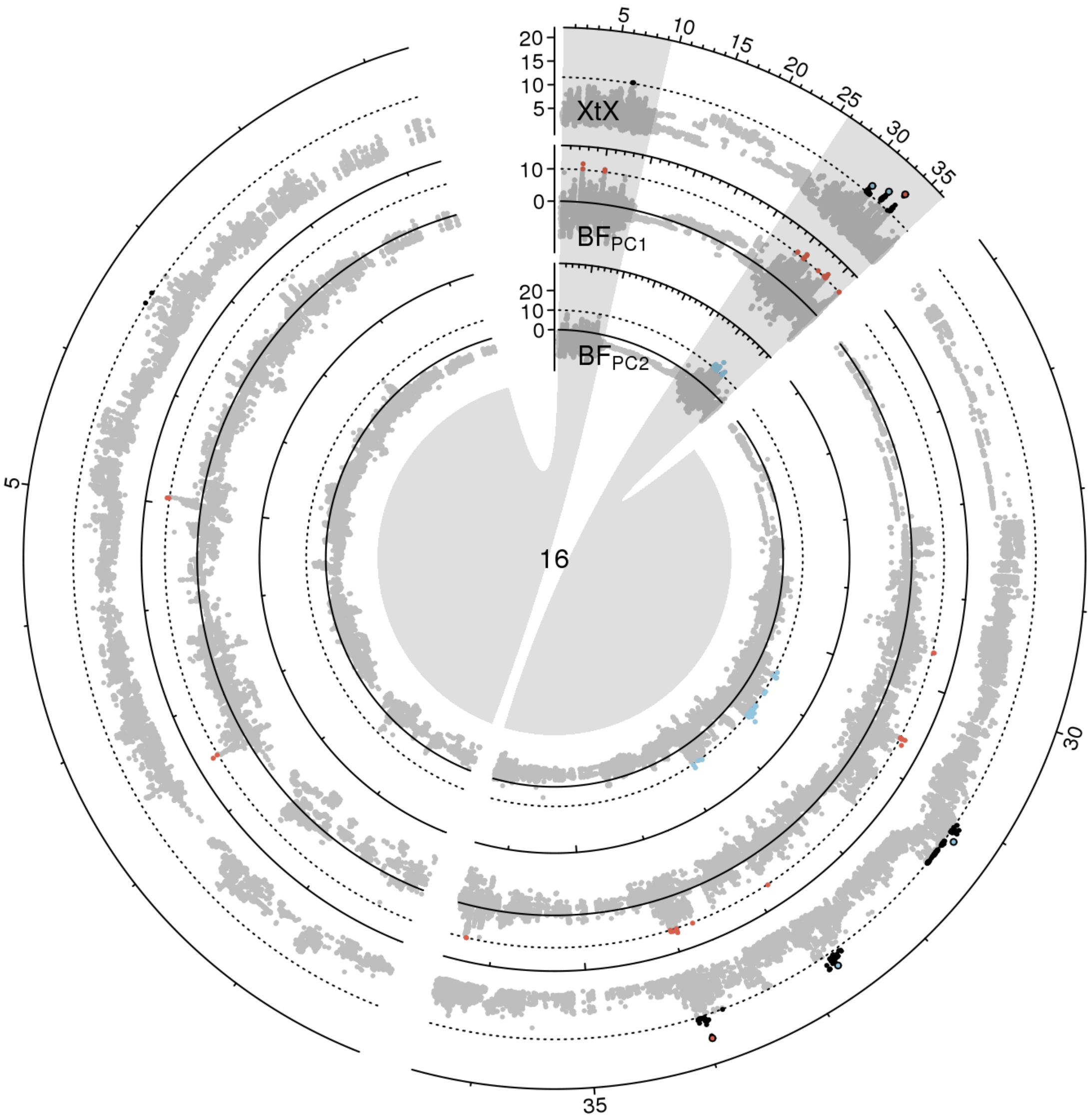


**Figure S 39:** Manhattan plot of the *XtX* statistic and of genotype-environment associations with the first two environmental principal components in Bayes factors for chromosome 16 in scenario B. Euchromatic regions on the chromosome arms are enlarged.

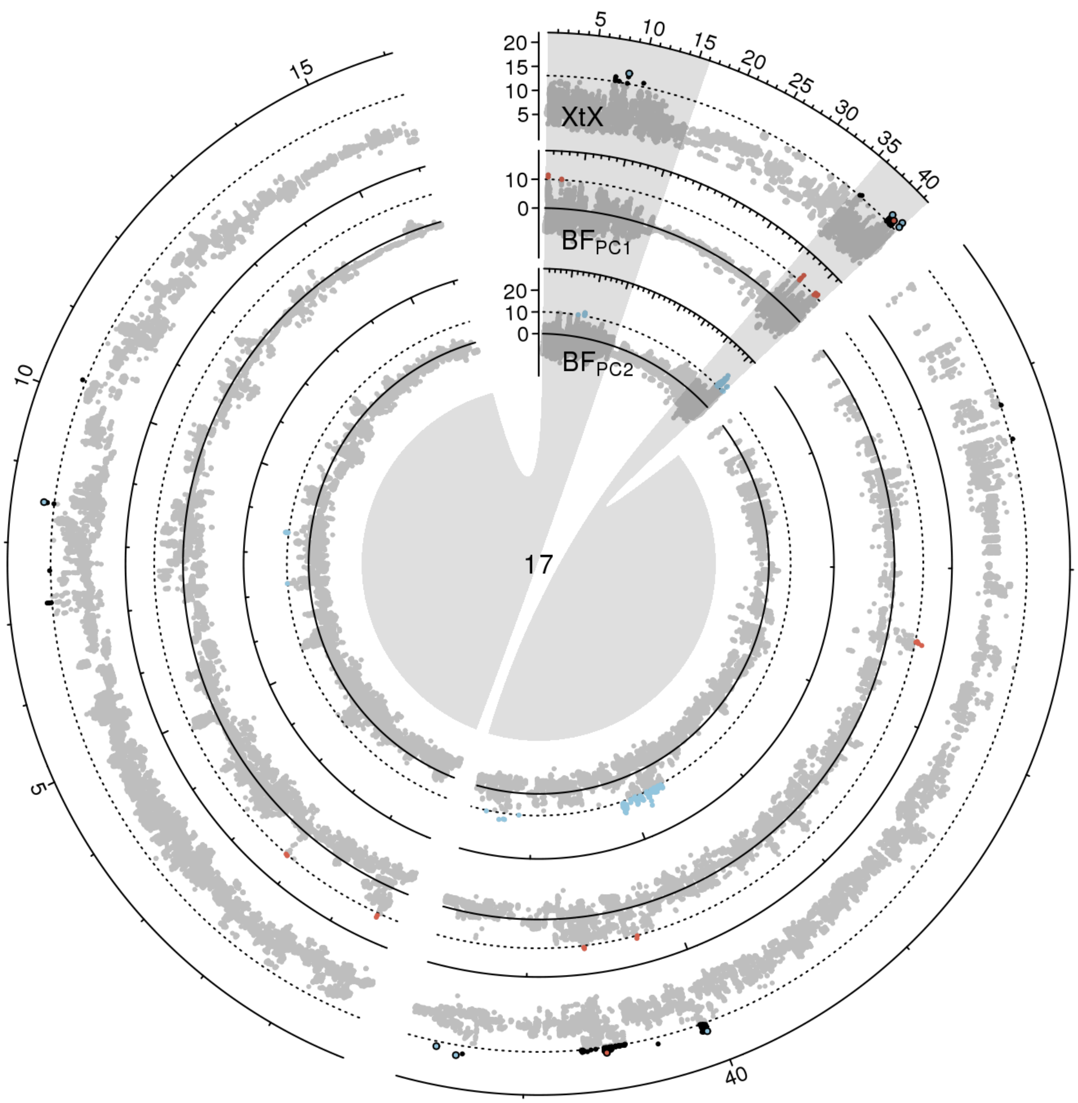


**Figure S 40:** Manhattan plot of the *XtX* statistic and of genotype-environment associations with the first two environmental principal components in Bayes factors for chromosome 17 in scenario A. Euchromatic regions on the chromosome arms are enlarged.

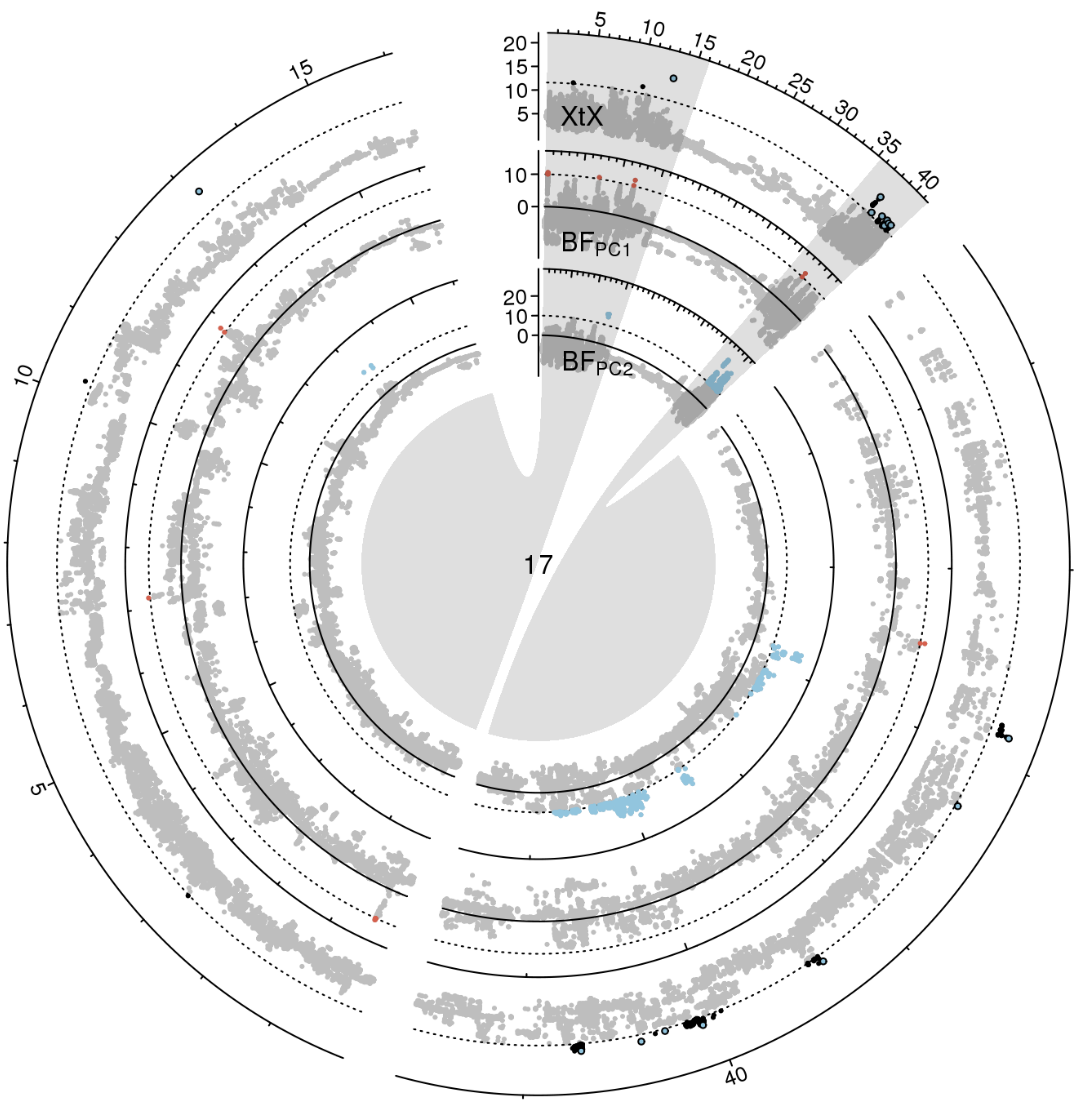


**Figure S 41:** Manhattan plot of the *XtX* statistic and of genotype-environment associations with the first two environmental principal components in Bayes factors for chromosome 17 in scenario B. Euchromatic regions on the chromosome arms are enlarged.

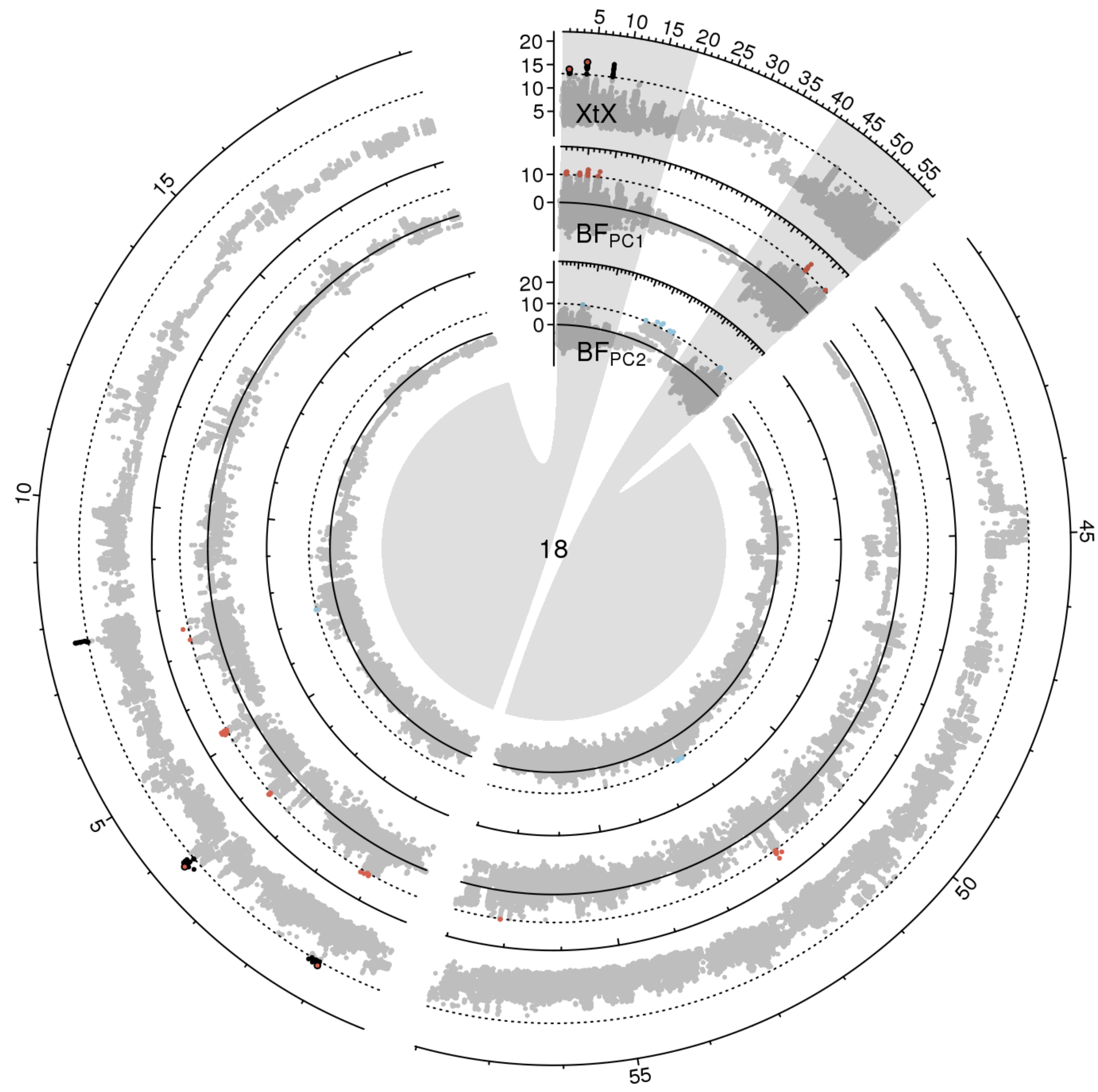


**Figure S 42:** Manhattan plot of the *XtX* statistic and of genotype-environment associations with the first two environmental principal components in Bayes factors for chromosome 18 in scenario A. Euchromatic regions on the chromosome arms are enlarged.

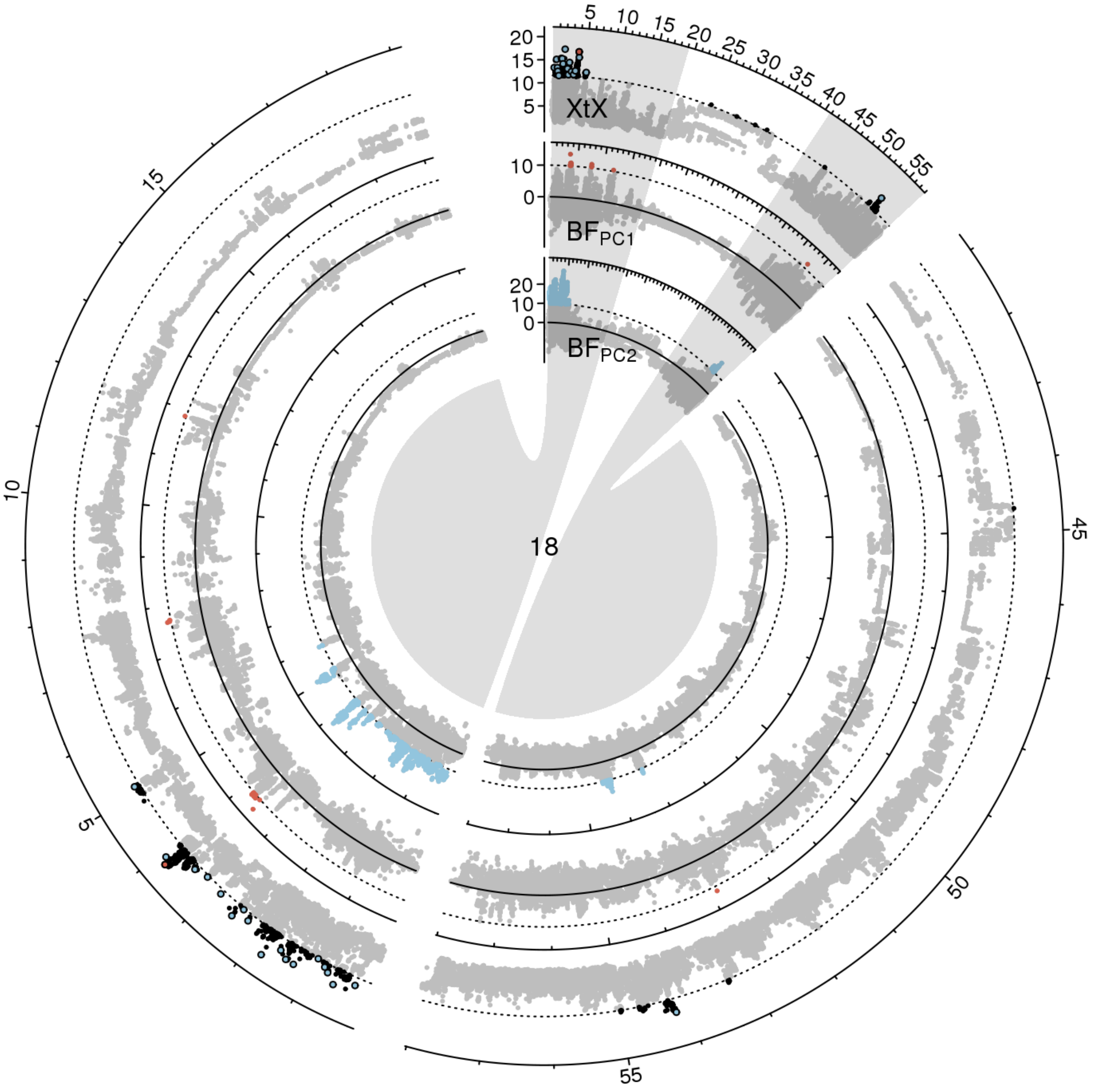


**Figure S 43:** Manhattan plot of the *XtX* statistic and of genotype-environment associations with the first two environmental principal components in Bayes factors for chromosome 18 in scenario B. Euchromatic regions on the chromosome arms are enlarged.

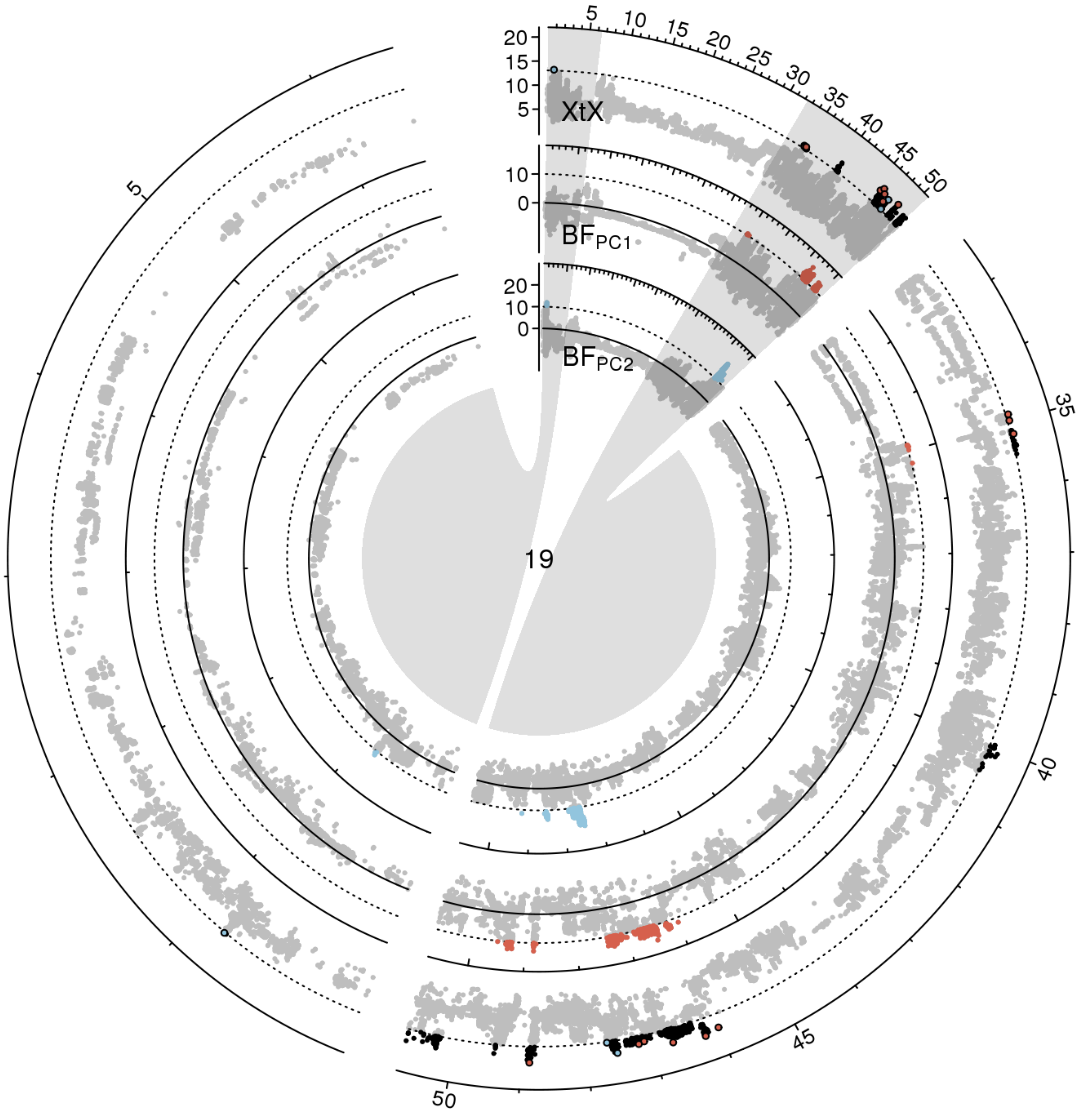


**Figure S 44:** Manhattan plot of the *XtX* statistic and of genotype-environment associations with the first two environmental principal components in Bayes factors for chromosome 19 in scenario A. Euchromatic regions on the chromosome arms are enlarged.

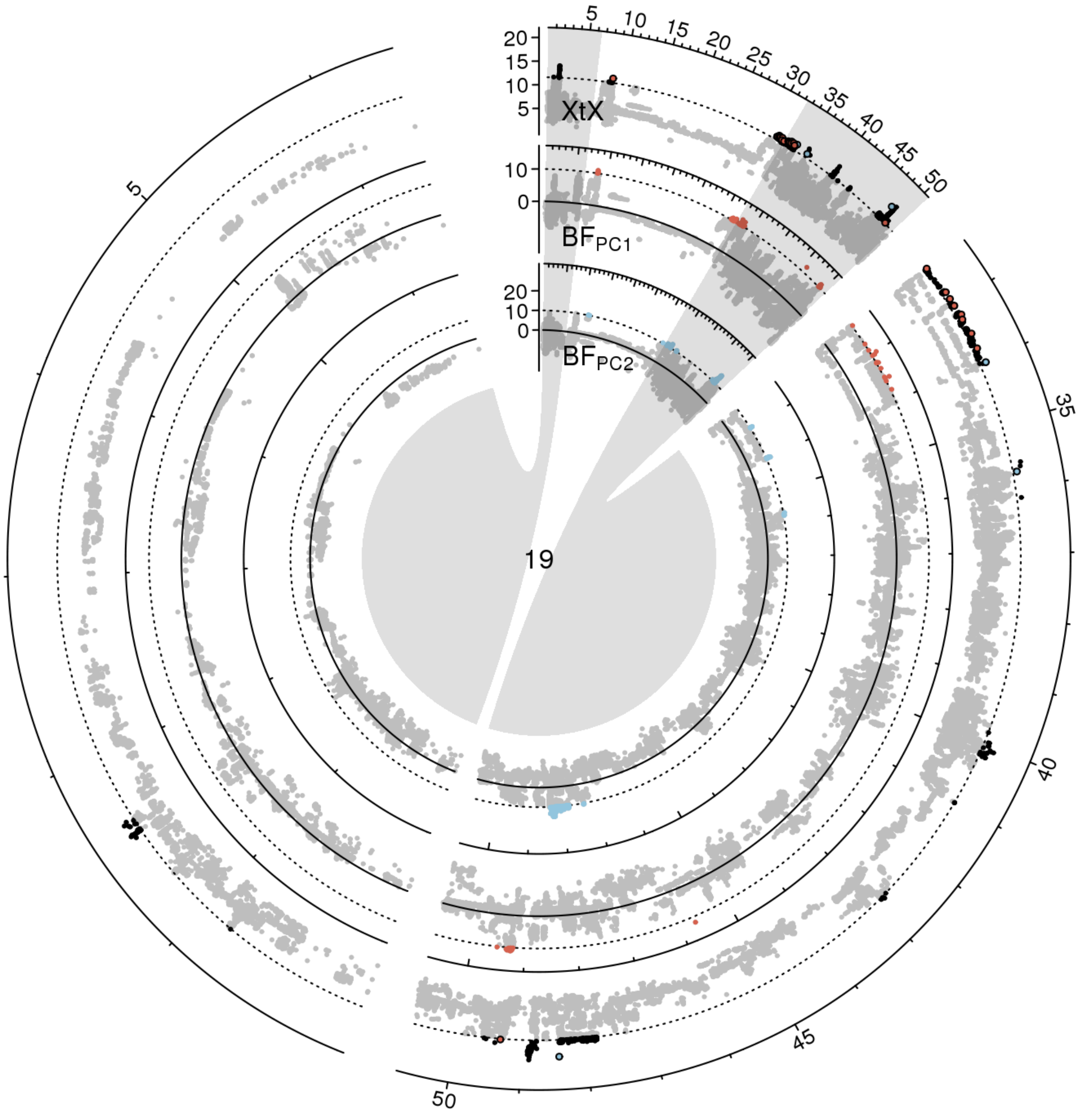


**Figure S 45:** Manhattan plot of the *XtX* statistic and of genotype-environment associations with the first two environmental principal components in Bayes factors for chromosome 19 in scenario B. Euchromatic regions on the chromosome arms are enlarged.

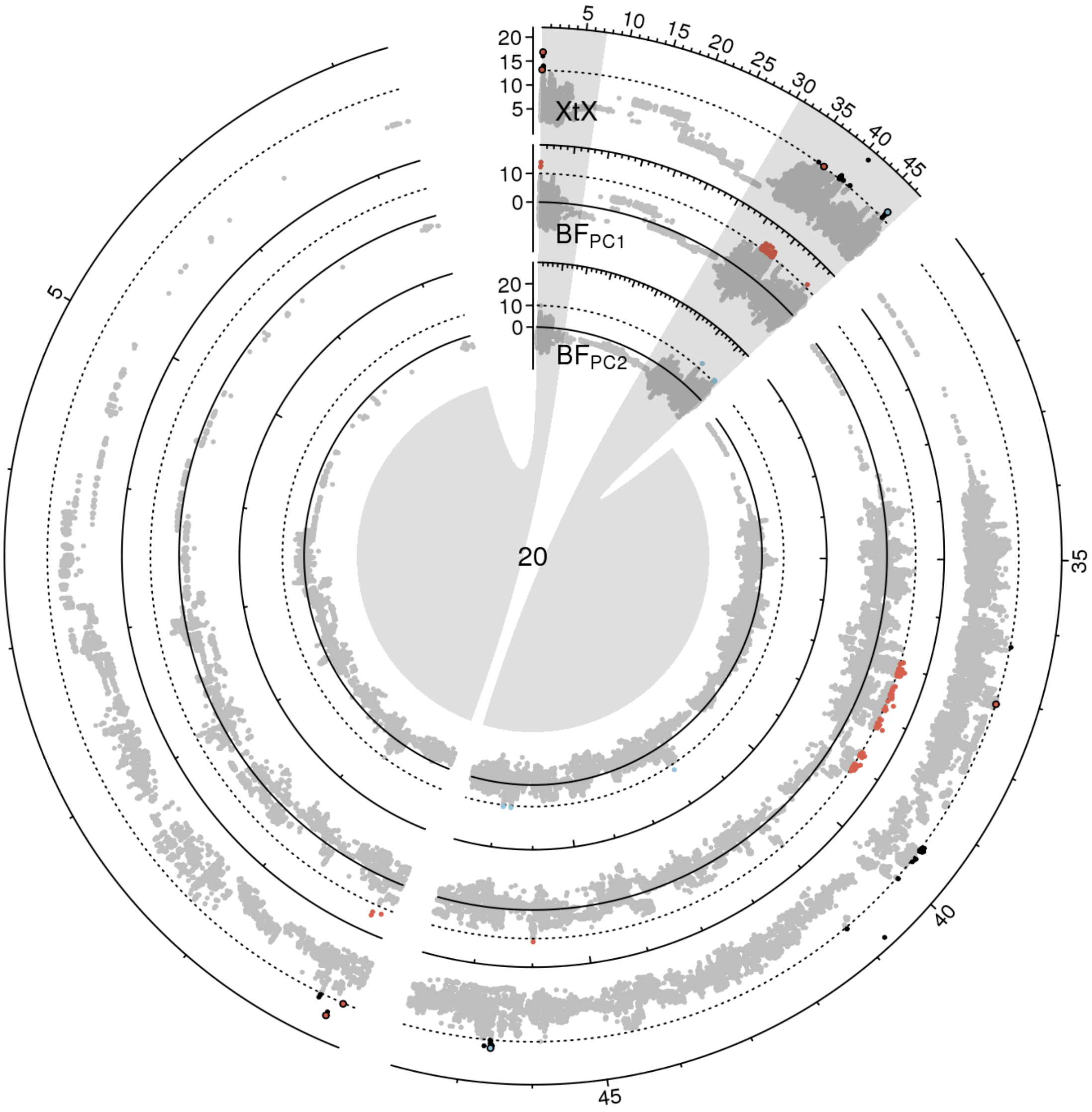


**Figure S 46:** Manhattan plot of the *XtX* statistic and of genotype-environment associations with the first two environmental principal components in Bayes factors for chromosome 20 in scenario A. Euchromatic regions on the chromosome arms are enlarged.

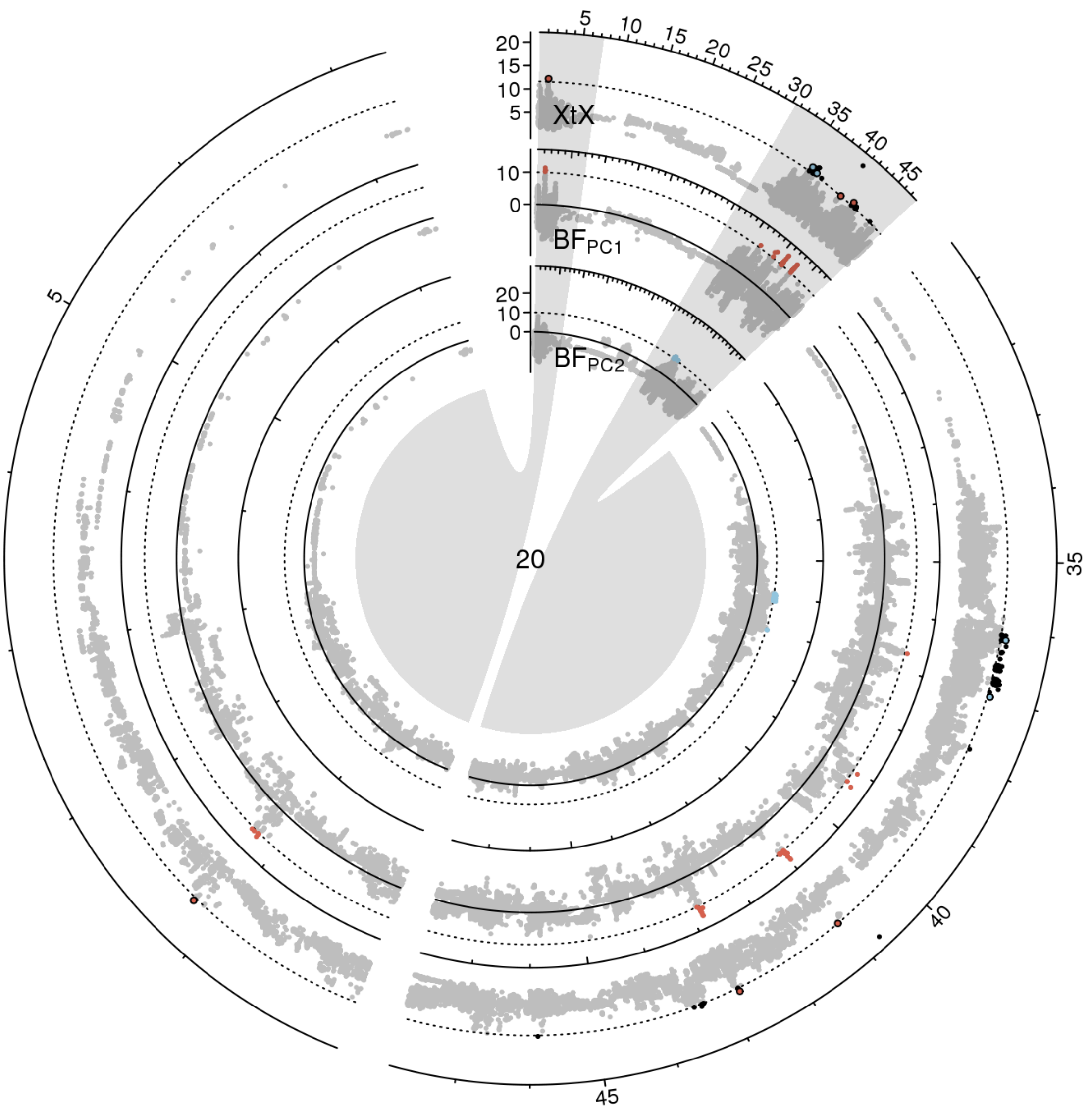


**Figure S 47:** Manhattan plot of the *XtX* statistic and of genotype-environment associations with the first two environmental principal components in Bayes factors for chromosome 20 in scenario B. Euchromatic regions on the chromosome arms are enlarged.

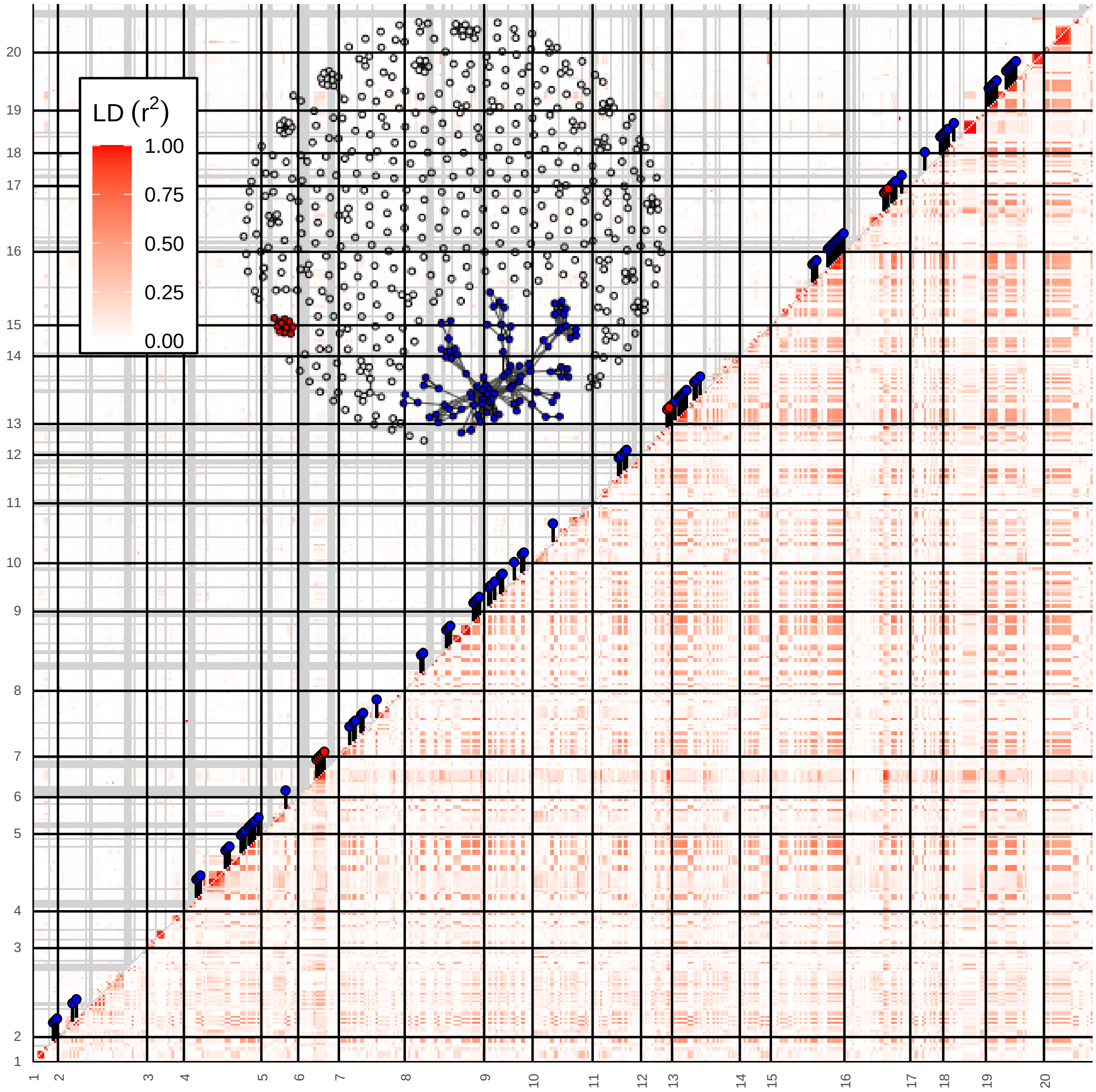


**Figure S 48:** LD estimates among 544 genomic regions with selection signatures observed in scenario A for germplasm groups from China (below diagonal) and in modern European soybean varieties (above diagonal). Red and blue pins on the diagnonal indicate regions that clustered in LD network analysis. Inset shows the LD network of the 544 genomic regions. Two clusters (92 and 10 regions, respectively) comprising LD connections exceeding the level of background LD 3-fold and a minimum physical distance of 5Mb are highlighted in blue and red.

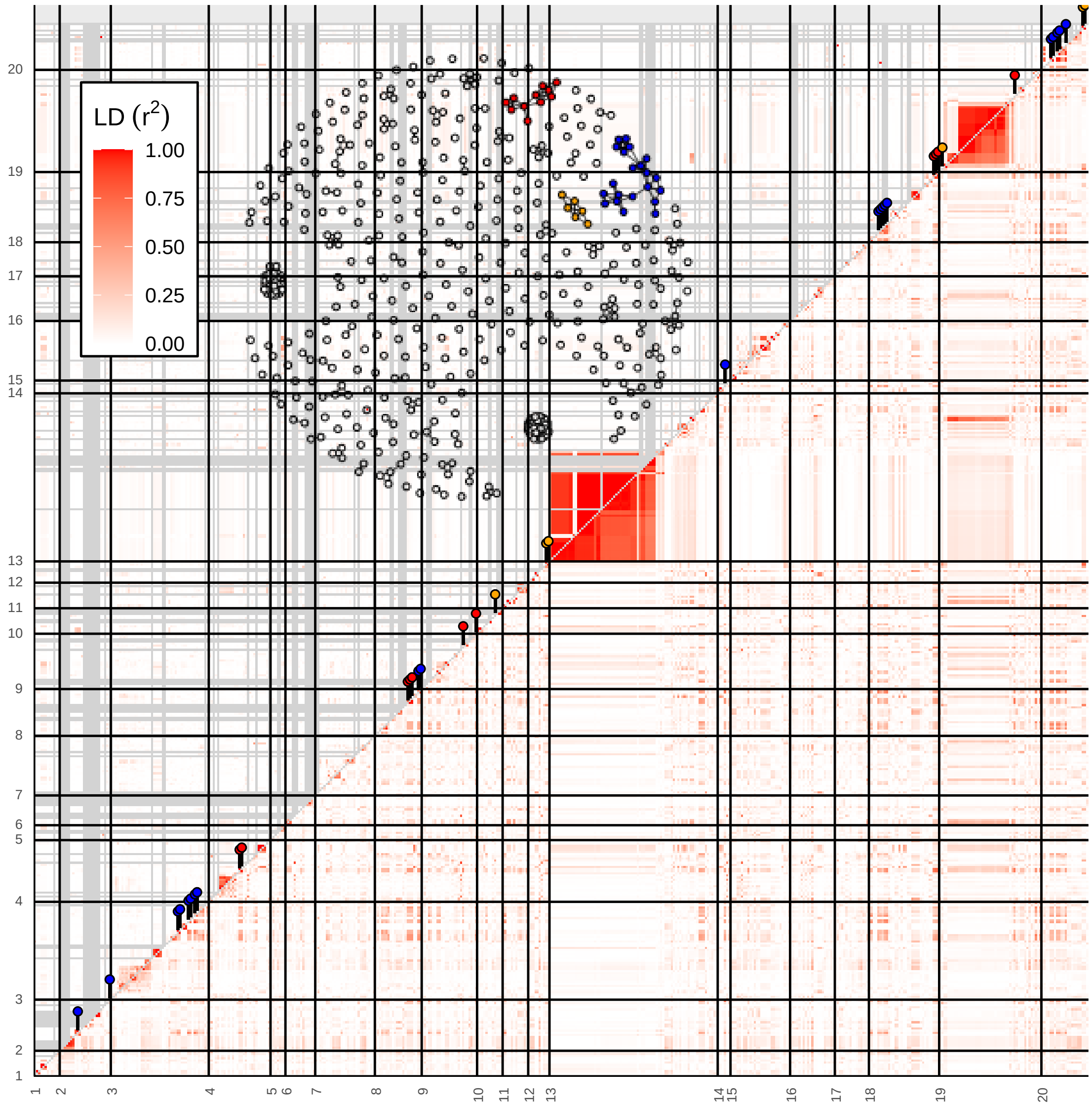


**Figure S 49:** LD estimates among 481 genomic regions with selection signatures observed in scenario B for germplasm groups from China (below diagonal) and in modern European soybean varieties (above diagonal). Red and blue pins on the diagnonal indicate regions that clustered in LD network analysis. Inset shows the LD network of the 481 genomic regions. Three clusters (21, 11 and 6 regions, respectively) comprising LD connections exceeding the level of background LD 3-fold and a minimum physical distance of 5Mb are highlighted in blue, red and orange.

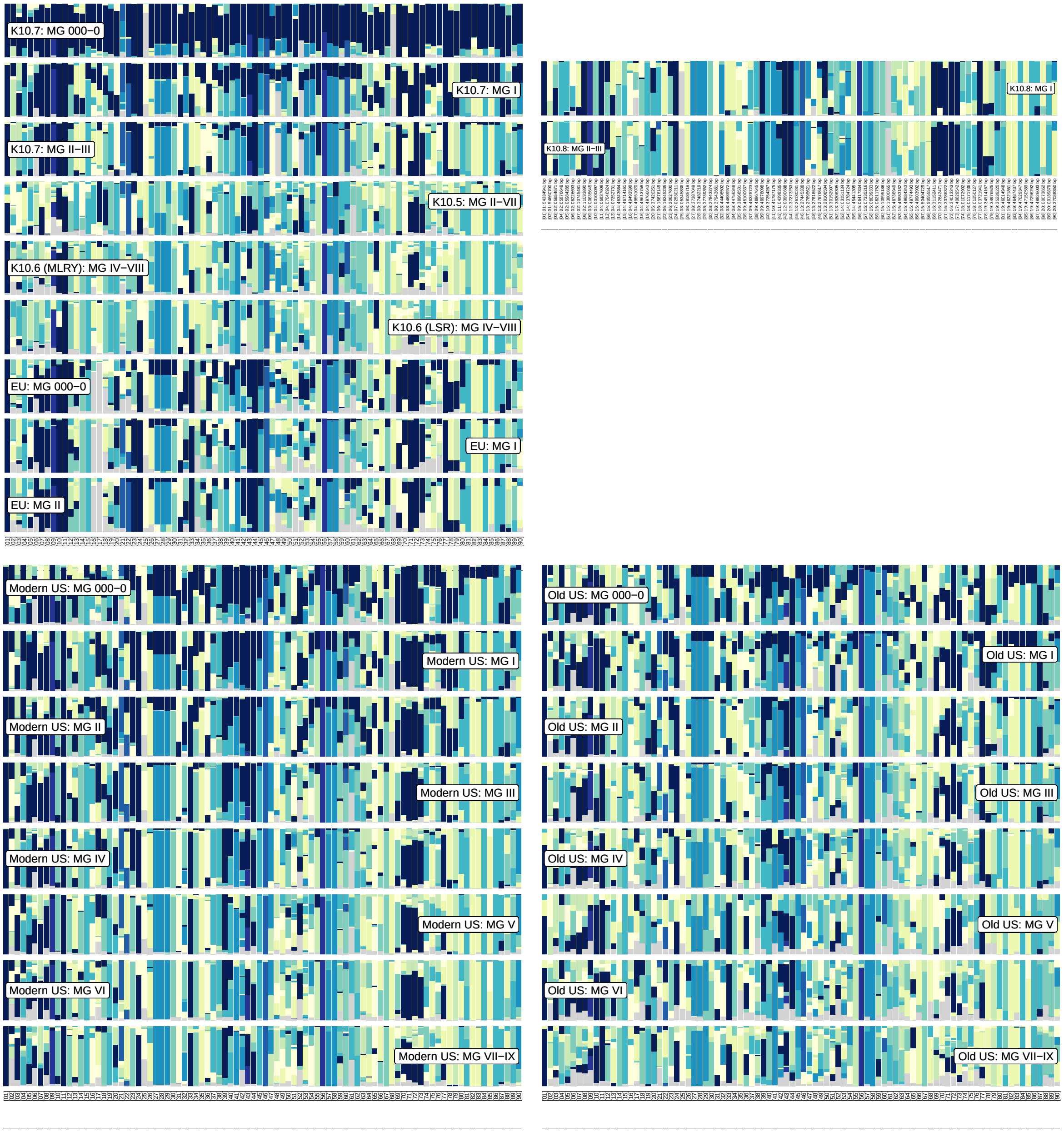


**Figure S 50:** Top left: Haplotype block proportions in germplasm groups from China (and modern European varieties) for 90 genomic regions with overlapping genetic differentiation and genotype-environment association signatures (PC1) observed in scenario A. Top right: Haplotype block proportions in the K10.8 subpopulation for the 90 genomic regions. Bottom left: Haplotype block proportions in modern US varieties for the 90 genomic regions. Bottom right: Haplotype block proportions in old US varieties for the 90 genomic regions.

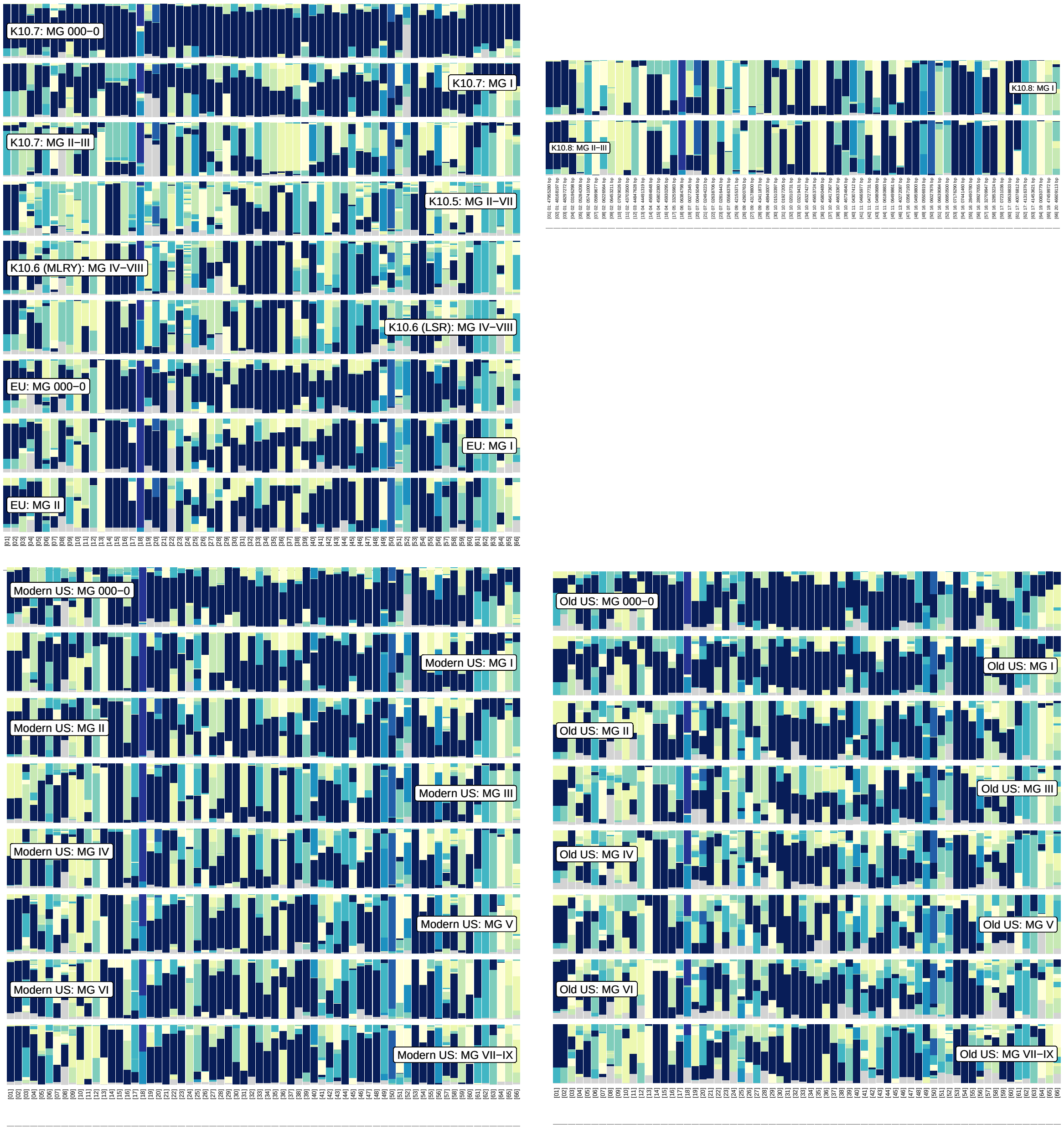


**Figure S 51:** Top left: Haplotype block proportions in germplasm groups from China (and modern European varieties) for 66 genomic regions with overlapping genetic differentiation and genotype-environment association signatures (PC2) observed in scenario A. Top right: Haplotype block proportions in the K10.8 subpopulation for the 66 genomic regions. Bottom left: Haplotype block proportions in modern US varieties for the 66 genomic regions. Bottom right: Haplotype block proportions in old US varieties for the 66 genomic regions.

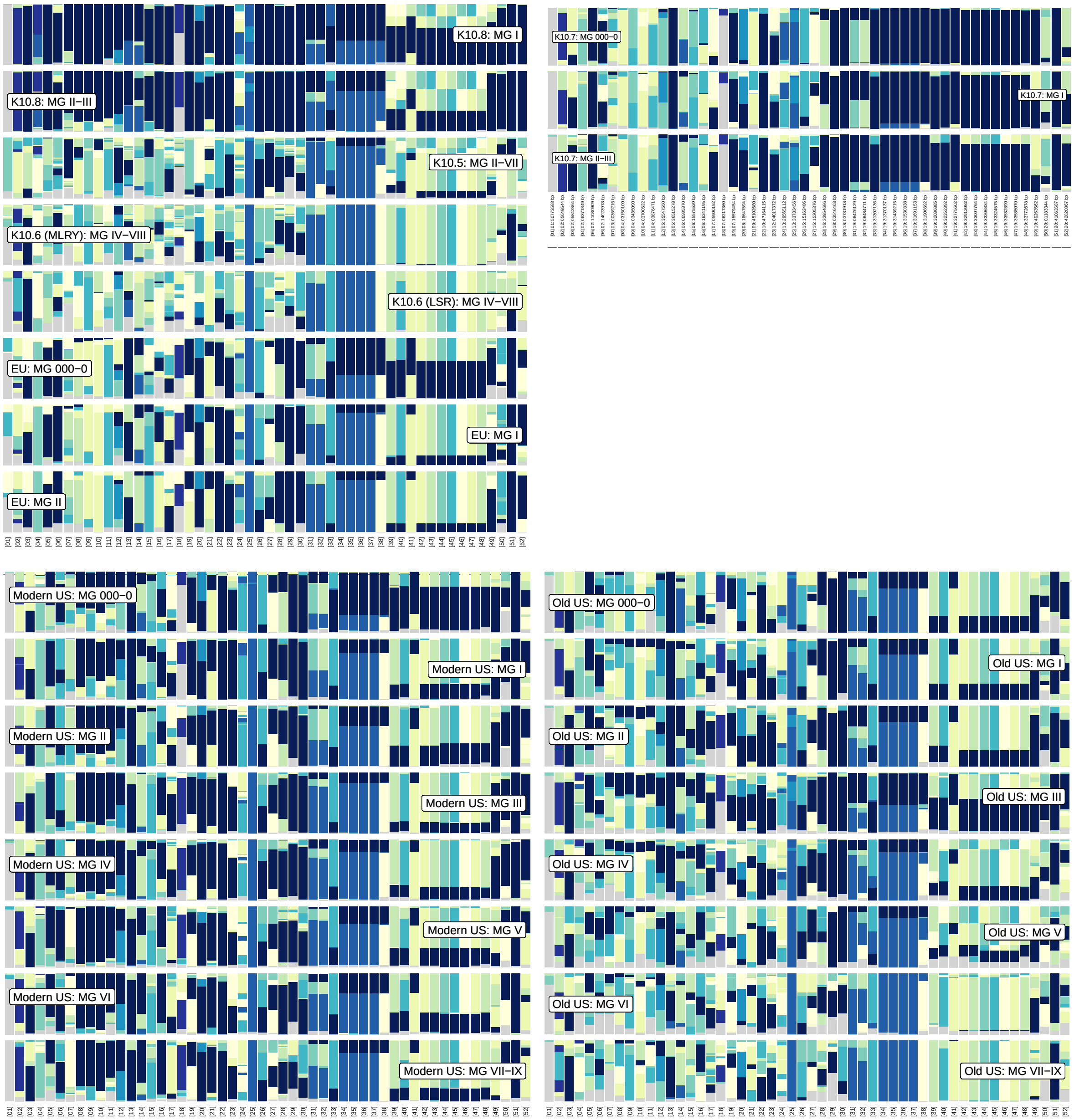


**Figure S 52:** Top left: Haplotype block proportions in germplasm groups from China (and modern European varieties) for 52 genomic regions with overlapping genetic differentiation and genotype-environment association signatures (PC1) observed in scenario B. Top right: Haplotype block proportions in the K10.7 subpopulation for the 52 genomic regions. Bottom left: Haplotype block proportions in modern US varieties for the 52 genomic regions. Bottom right: Haplotype block proportions in old US varieties for the 52 genomic regions.

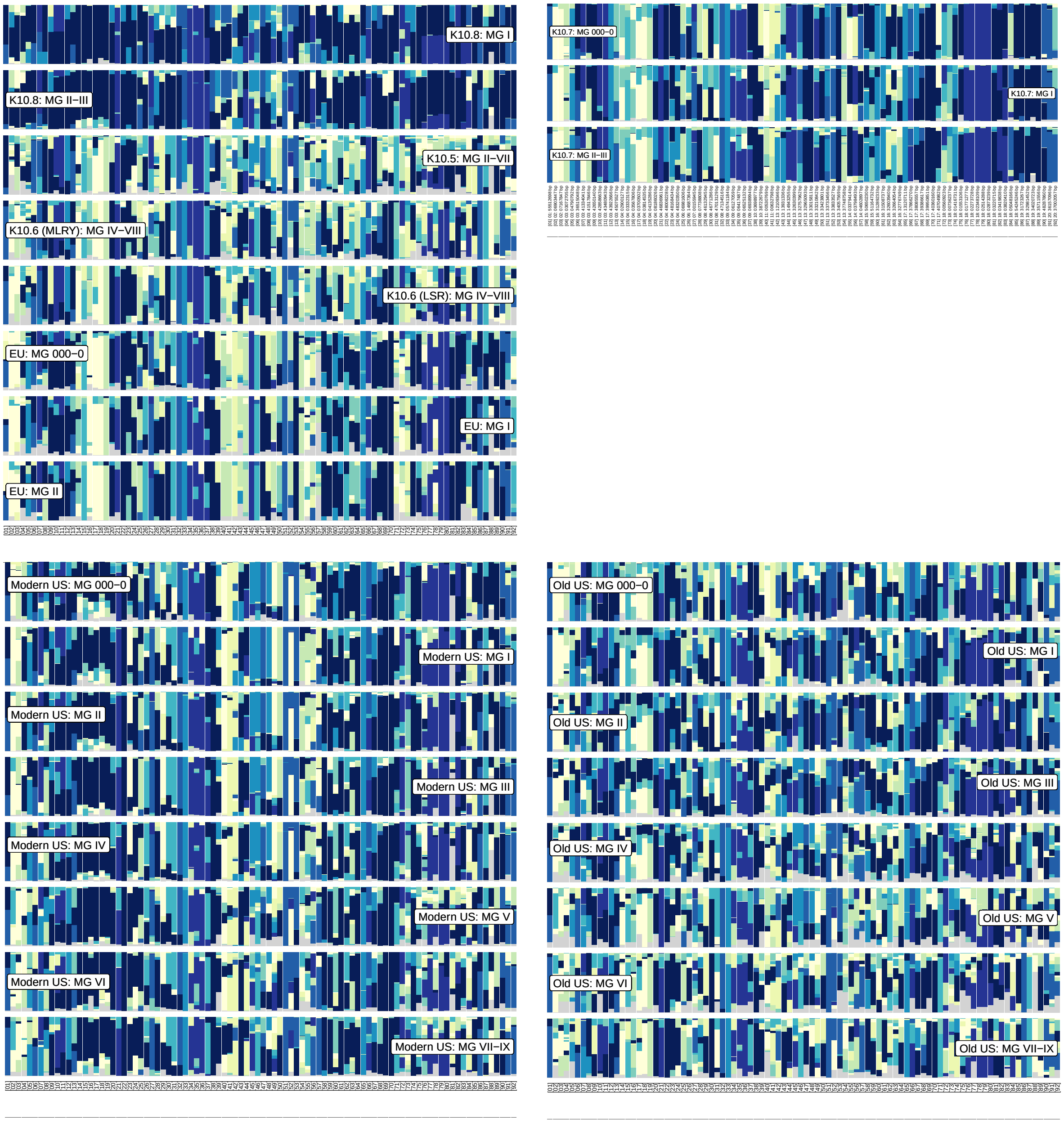


**Figure S 53:** Top left: Haplotype block proportions in germplasm groups from China (and modern European varieties) for 92 genomic regions with overlapping genetic differentiation and genotype-environment association signatures (PC2) observed in scenario B. Top right: Haplotype block proportions in the K10.8 subpopulation for the 92 genomic regions. Bottom left: Haplotype block proportions in modern US varieties for the 92 genomic regions. Bottom right: Haplotype block proportions in old US varieties for the 92 genomic regions.

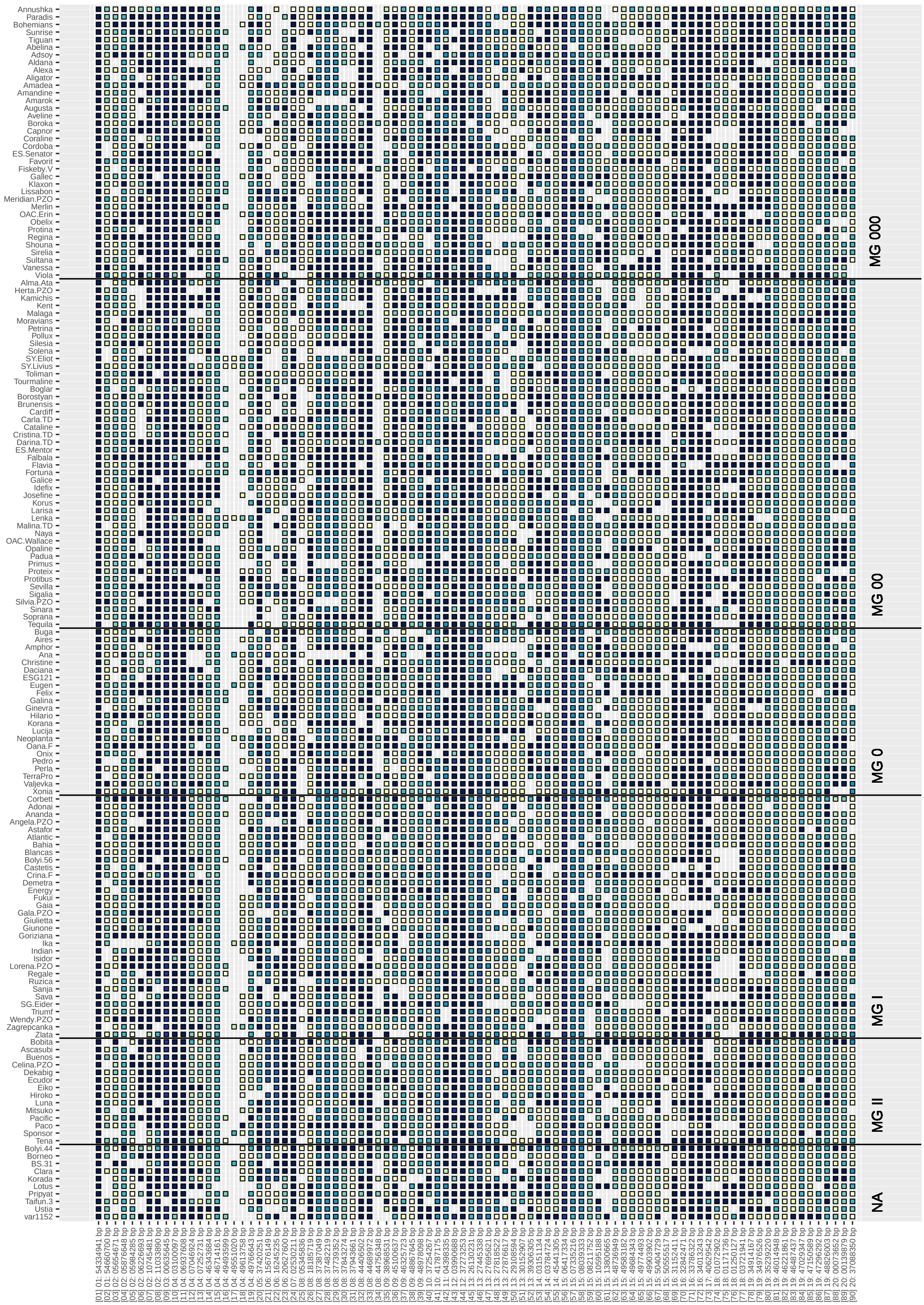

**Figure S 54:** Haplotype blocks in modern European varieties for 90 genomic regions with overlapping genetic differentiation and genotype-environment association signatures (PC1) observed in scenario A.

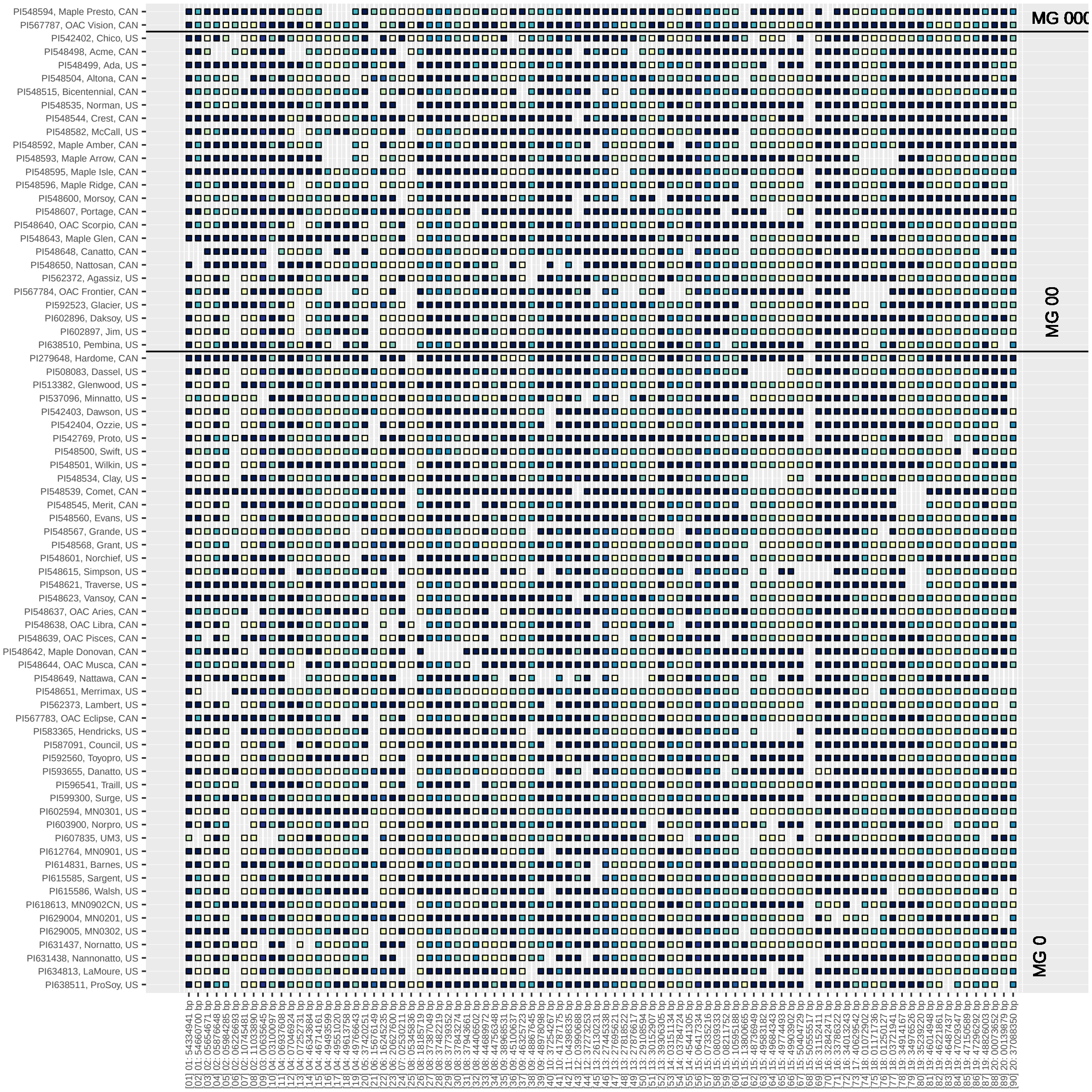

**Figure S 55:** Haplotype blocks in modern varieties from the USA and Canada for 90 genomic regions with overlapping genetic differentiation and genotype-environment association signatures (PC1) observed in scenario A.
